# TORC1 integrates metabolic state transitions during aging

**DOI:** 10.64898/2026.05.18.726092

**Authors:** Trishia Yi Ning Cheng, Arshia Naaz, Mingtong Gao, Wu Hao, Yizhong Zhang, Liang Cui, Jovian Jing Lin, Sonia Yogasundaram, Nashrul Afiq Faidzinn, Ong Yong Qing Victoria, Rajkumar Dorajoo, Brian K. Kennedy, Mohammad Alfatah

## Abstract

Target of rapamycin complex 1 (TORC1) coordinates nutrient availability with anabolic metabolism, yet how TORC1-linked metabolic states influence cellular aging remains unclear. Using genetic, transcriptomic, metabolomic, and pharmacological analyses in prototrophic *Saccharomyces cerevisiae*, we identify context-dependent metabolic state transitions that uncouple cellular proliferation from long-term survival during aging. Loss of the SEACIT complex, conserved with mammalian GATOR1, establishes a low-flux metabolic state characterized by coordinated remodeling of nitrogen, nucleotide, and central carbon metabolism during stationary phase. Pharmacological restraint of nucleotide, glycolytic, or sterol metabolism converges on similar adaptive metabolic programs, whereas perturbations that impair mitochondrial respiratory capacity destabilize these states and reduce survival. Integrative analyses in human cancer and primary cells further reveal that diverse metabolic perturbations converge on mTORC1 suppression but generate distinct mitochondrial and stress responses depending on nutrient availability. Together, these findings demonstrate that TORC1 integrates metabolic state transitions in which nutrient availability and mitochondrial function determine cellular survival during aging.

## INTRODUCTION

Cellular aging is fundamentally shaped by nutrient availability and the ability of cells to dynamically allocate metabolic resources between anabolic growth, metabolic maintenance, and stress adaptation ^1–4^. Central to this coordination is target of rapamycin complex 1 (TORC1), a conserved nutrient-sensing kinase complex that integrates amino acid, energy, and growth-factor signals to regulate protein synthesis, central carbon metabolism, lipid biosynthesis, and mitochondrial activity ^5–13^. Through these outputs, TORC1 coordinates adaptive metabolic responses to fluctuating nutrient conditions. Dysregulated TORC1 signaling has been strongly linked to aging and age-related diseases, including cancer, where elevated TORC1 activity promotes anabolic metabolism and uncontrolled cellular growth ^10,14–19^. Understanding how TORC1 integrates metabolic state transitions during aging is therefore central to both aging biology and cancer metabolism.

Cells frequently encounter nutrient limitation in physiological environments. Aging tissues, ischemic regions, and solid tumors often experience restricted nutrient and oxygen availability, requiring cells to adopt metabolic programs that sustain survival under metabolic stress ^2,3,20–22^. Cancer cells in particular persist in nutrient-poor microenvironments through metabolic rewiring that supports continued adaptation despite limited resources ^20–25^. Consistent with this view, TORC1 signaling exerts broad effects on organismal physiology. Reduced TORC1 activity has been associated with delayed aging and lifespan extension across species, whereas dysregulated or hyperactive TORC1 signaling contributes to age-associated diseases, including cancer ^7,10,15,26–28^. Importantly, responses to dietary or metabolic interventions are highly context dependent. Dietary restriction can extend lifespan in some genetic backgrounds while producing neutral or adverse outcomes in others, indicating that cellular metabolic state, rather than nutrient availability alone, determines survival outcomes ^29–31^. These observations suggest that nutrient-sensing pathways coordinate adaptive metabolic states that determine cellular survival during aging.

TORC1 signaling operates through a modular regulatory architecture composed of multiple upstream nutrient-sensing inputs and downstream effector branches ^5–7,9,12,13^. In *Saccharomyces cerevisiae*, amino acid availability is transmitted to TORC1 through the Rag-family GTPases Gtr1 and Gtr2, which are anchored to the vacuolar membrane by the EGO complex and regulated by upstream modules including Lst4/Lst7, Vam6, and the SEA protein complexes ^12,32–37^. Downstream of TORC1, signaling is conveyed through discrete effector branches including the AGC-family kinase Sch9 and phosphatase-associated modules involving Sit4 and its regulatory subunits Rrd1 and Rrd2 ^11,12,38,39^. Importantly, this regulatory architecture is broadly conserved across eukaryotes: yeast Gtr1/Gtr2 correspond to mammalian RagA/B and RagC/D, the EGO complex is functionally analogous to the lysosomal Ragulator complex, and several upstream regulatory modules have conserved counterparts in higher eukaryotes ^12,40–43^.

The presence of multiple upstream regulators and downstream effector branches suggests that TORC1 integrates interconnected metabolic programs rather than functioning as a single linear signaling pathway. Distinct modules within this network influence nutrient sensing, anabolic metabolism, mitochondrial activity, and metabolic flux through partially overlapping mechanisms, raising the possibility that perturbations at different regulatory nodes generate qualitatively distinct metabolic states rather than simply scaling TORC1 activity. However, how specific TORC1 regulatory modules coordinate metabolic state transitions during aging remains unclear, as genetic perturbations within the TORC1 network frequently generate divergent physiological outcomes that cannot be explained solely by reduced growth or anabolic output.

Here, we systematically dissect upstream regulators and downstream effectors of TORC1 in prototrophic *Saccharomyces cerevisiae* under nutrient-limited conditions. By integrating genetic perturbations with transcriptomic, metabolomic, and pharmacological analyses across proliferative and stationary phases, we identify context-dependent metabolic state transitions that uncouple cellular proliferation from long-term survival during aging. We show that loss of the SEACIT complex establishes a low-flux metabolic state characterized by coordinated remodeling of nitrogen, nucleotide, and central carbon metabolism during stationary phase. Pharmacological restraint of nucleotide, glycolytic, or sterol metabolism converges on similar adaptive metabolic programs, whereas perturbations that impair mitochondrial respiratory capacity destabilize these states and reduce survival. Complementary analyses in human cancer and primary cells further reveal that diverse metabolic perturbations converge on mTORC1 suppression but generate distinct mitochondrial and stress responses depending on nutrient availability. Together, these findings establish a framework in which TORC1 integrates context-dependent metabolic state transitions that determine cellular adaptation and survival during aging.

## RESULTS

### TORC1 network perturbations uncouple cellular growth from long-term survival during aging

To examine how TORC1 regulatory modules influence cellular proliferation and survival under nutrient limitation, we analyzed upstream regulators and downstream effectors of TORC1 **(Figure 1A)** in prototrophic *Saccharomyces cerevisiae* grown in synthetic defined (SD) medium. SD medium imposes constrained nutrient availability, enabling resolution of metabolic-state differences beyond simple growth effects. Early growth and long-term survival were quantified using high-throughput liquid culture assays.

**Figure 1.**
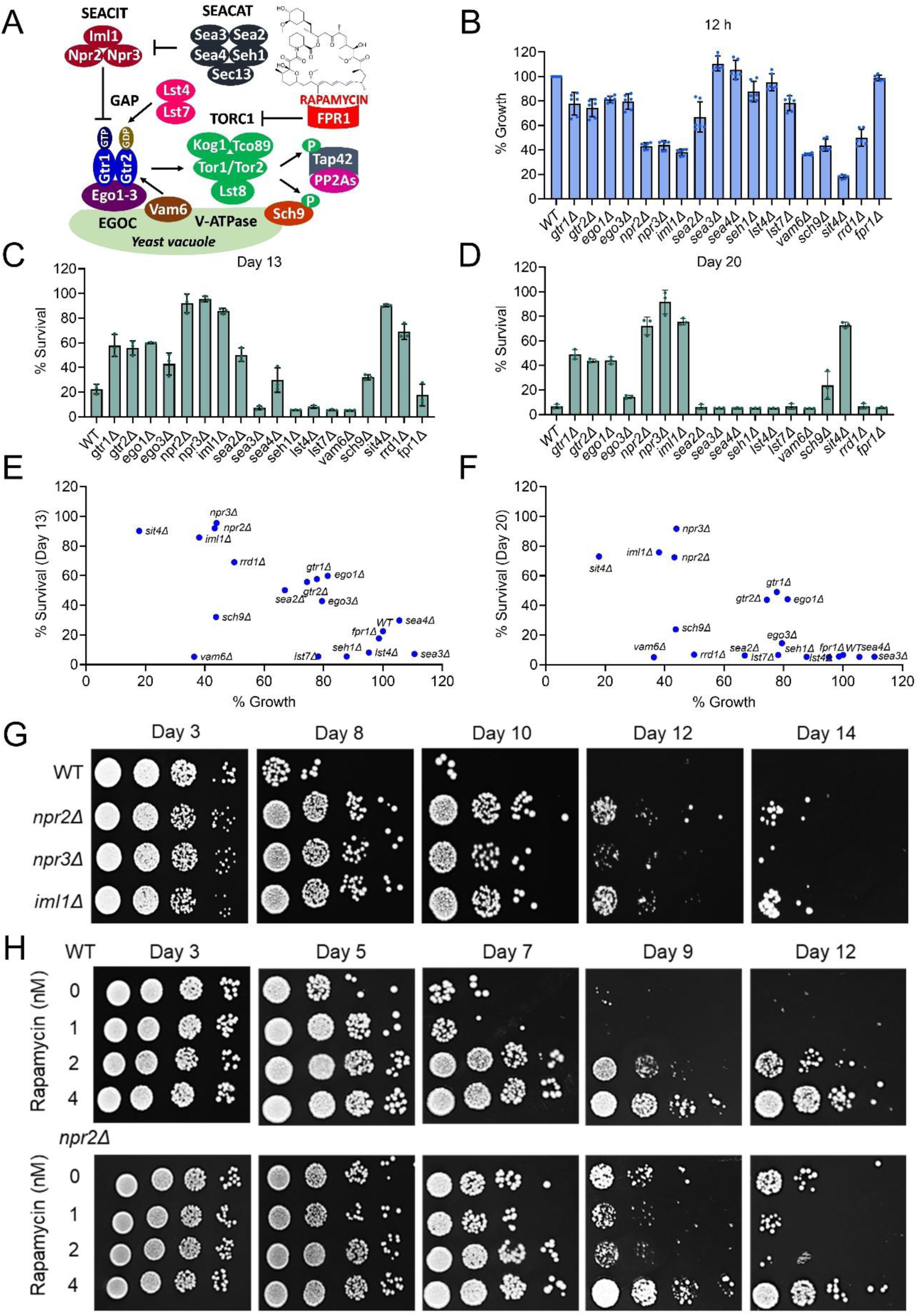
Distinct TORC1 regulatory modules uncouple growth from long-term cellular survival during cellular aging. (A) Schematic representation of TORC1 regulation in Saccharomyces cerevisiae, highlighting upstream regulatory complexes (SEACIT, SEACAT, EGOC, Lst4–Lst7, Vam6), core TORC1 components, downstream effectors, and inhibition by rapamycin via Fpr1. (B) Relative early growth of wild-type and indicated TORC1 pathway mutants measured after 12 h of culture in synthetic defined (SD) medium. Values are normalized to wild-type (mean ± SD, n = 6). (C–D) Long-term cellular survival assessed during stationary phase, shown as the percentage of viable cells on day 13 (C) and day 20 (D) of culture in SD medium (mean ± SD, n = 3). (E–F) Relationship between early growth (12 h) and long-term survival at day 13 (E) and day 20 (F). (G) Survival of wild-type and SEACIT-deficient strains (*npr2Δ*, *npr3Δ*, *iml1Δ*) over time assessed by spot dilution assays at the indicated days of stationary phase. (H) Survival analysis of wild-type and *npr2Δ* cells treated with increasing concentrations of rapamycin, assessed by spot dilution assays during stationary phase.

Deletion of several upstream positive regulators of TORC1 ^11,12^, including *gtr1Δ*, *gtr2Δ*, *ego1Δ*, and *ego3Δ*, resulted in reduced early growth accompanied by increased long-term survival **(Figures 1B–1D and S1A–S1D)**. Similarly, deletion of downstream TORC1 effectors (*sch9Δ*, *sit4Δ*, *rrd1Δ*) impaired growth while enhancing survival. These phenotypes are consistent with reduced TORC1 signaling output under nutrient-limited conditions and indicate that attenuation of specific TORC1 inputs or outputs can favor maintenance-oriented states.

The SEACAT complex also functions as a positive regulator of TORC1 ^11,12^. Deletion of *SEA2* and *SEA4* resulted in modest increases in long-term survival **(Figures 1B–1D and S1A–S1D)**, whereas deletion of other SEACAT components (*SEA3*, *SEH1*) did not enhance survival, indicating component-specific effects. Similarly, deletion of *LST4* or *LST7* produced variable growth and survival phenotypes and did not promote survival **(Figures 1B–1D and S1A–S1D)**. These observations suggest that reduced TORC1 signaling does not uniformly impose a survival-favorable metabolic state.

The SEACIT complex (*NPR2*, *NPR3*, *IML1*) functions as a negative regulator of TORC1 ^11,12^. Based on this role, loss of SEACIT would be expected to enhance TORC1 activity and promote growth. Instead, *npr2Δ*, *npr3Δ*, and *iml1Δ* cells exhibited reduced early growth in SD medium **(Figures 1B and S1A–S1C)**, consistent with previous observations in glucose-based minimal conditions ^44,45^. Strikingly, despite acting as negative regulators of TORC1, deletion of *NPR2*, *NPR3*, or *IML1* robustly increased long-term survival while simultaneously reducing growth **(Figures 1C**, **1D and S1D)**.

To quantify the relationship between proliferation and survival, we plotted long-term survival against early growth across TORC1 mutants **(Figures 1E and 1F)**. Overall, growth capacity showed a poor correlation with late-life survival. SEACIT mutants clustered in a regime characterized by reduced growth yet high survival, whereas other upstream TORC1 regulators, including *vam6Δ*, exhibited reduced growth accompanied by reduced survival.

These data indicate that uncoupling of growth and survival is module-specific within the TORC1 network rather than a general consequence of slowed proliferation.

The reduced growth of SEACIT mutants in SD medium was TORC1-dependent, as low-dose rapamycin partially restored growth, consistent with previous reports **(Figures S1E and S1F)** ^44,45^. We next examined whether this altered growth–survival relationship extended to pharmacological TORC1 inhibition. Spot dilution assays independently validated survival phenotypes observed in high-throughput liquid culture assays across all deletion strains **(Figures 1G and S1G)**.

Rapamycin treatment increased long-term survival in wild-type cells grown in SD medium **(Figures 1H and S1H)**, consistent with its established effects on TORC1-regulated metabolism ^17–19,46^. Unexpectedly, under identical nutrient-limited conditions, low concentrations of rapamycin reduced survival in the SEACIT-deficient *npr2Δ* background. These findings demonstrate that the survival consequences of TORC1 inhibition are strongly context dependent.

### SEACIT governs state-dependent transcriptional remodeling across growth and stationary phases

The growth–survival uncoupling observed in SEACIT-deficient cells **(Figure 1)** suggests that loss of SEACIT imposes a distinct metabolic state rather than uniformly attenuating TORC1 output. To define how SEACIT influences gene regulation across physiological states, we performed transcriptomic profiling of *npr2Δ* and wild-type cells during exponential growth (5 h) and stationary phase (72 h) in SD medium. Principal component analysis revealed clear separation by both genotype and physiological state **(Figures S2A and S2B)**, indicating that loss of *NPR2* drives pronounced, state-dependent transcriptional remodeling.

Differential expression analysis identified extensive and largely non-overlapping transcriptional changes between exponential growth and stationary phase. During exponential growth, 283 genes were differentially expressed in *npr2Δ* cells, whereas *583* genes were differentially expressed during stationary phase, with only 58 genes shared between conditions **(Figures 2A, 2B and Table S1)**. Of the growth-phase–specific genes, 201 were upregulated and 82 were downregulated, whereas during stationary phase 90 genes were uniquely upregulated and 494 were downregulated, indicating a substantial shift in transcriptional output as cells transition from proliferative to non-proliferative states.

**Figure 2.**
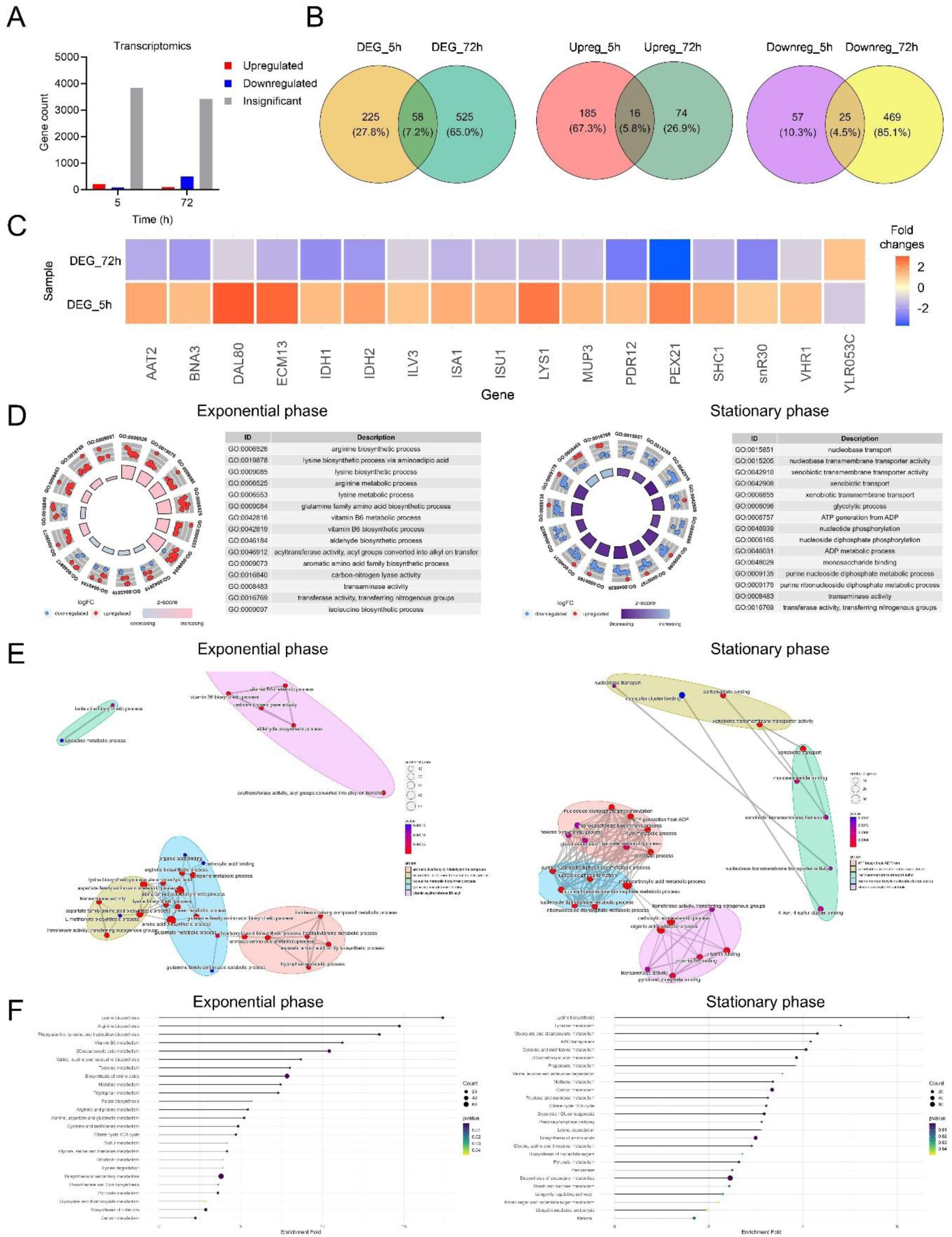
NPR2-dependent transcriptional remodeling is state dependent across growth and stationary phases. (A) Bar plot showing the number of differentially expressed genes (DEGs) in *npr2Δ* relative to wild-type during exponential growth (5 h) and stationary phase (72 h) in synthetic defined (SD) medium. (B) Venn diagrams illustrating overlap and phase-specific DEGs between exponential and stationary phases. DEGs are separated into upregulated and downregulated gene sets, highlighting limited transcriptional overlap and extensive state-specific *NPR2*-dependent regulation. (C) Heatmap of genes significantly regulated by *NPR2* in both exponential and stationary phases but exhibiting divergent expression patterns across physiological states, illustrating phase-dependent rewiring rather than uniform transcriptional effects. (D) Circular Gene Ontology (GO) enrichment plots for DEGs identified during exponential growth (left) and stationary phase (right), highlighting distinct biological processes associated with NPR2-dependent transcriptional programs in proliferative versus non-proliferative states. (E) Network representation of enriched GO terms clustered by functional similarity for exponential growth (left) and stationary phase (right). Nodes represent GO terms, with edge connectivity reflecting shared gene membership. (F) Dot plots showing significantly enriched KEGG pathways among DEGs during exponential growth (left) and stationary phase (right). Dot size corresponds to gene count and color indicates enrichment significance.

Among the 58 shared differentially expressed genes, expression patterns diverged across physiological states. 16 genes were consistently upregulated, and 25 genes were consistently downregulated across both conditions, while 17 genes exhibited opposing regulation between exponential growth and stationary phase **(Figure 2C)**. Notably, genes involved in nitrogen regulation and amino acid metabolism, including *DAL80*, *ECM30*, *LYS1*, and *AAT2*, were upregulated during growth but downregulated during stationary phase, highlighting dynamic rewiring of nitrogen-associated programs upon loss of SEACIT.

Gene ontology analysis revealed that during exponential growth, *npr2Δ* cells strongly upregulated anabolic pathways centered on amino acid biosynthesis and nitrogen metabolism, including arginine, lysine, glutamine-family, branched-chain, and aromatic amino acid biosynthetic processes, as well as vitamin B6 metabolism and transaminase activity **(Figures 2D–2F and Extended Data 1)**. Gene-level analysis confirmed coordinated induction of these pathways **(Figure S3A)**, consistent with a compensatory anabolic transcriptional program that supports growth despite impaired TORC1 regulatory balance. In contrast, several genes linking mitochondrial metabolism and carbon–nitrogen integration, including *CIT1*, *CIT2*, and *GDH1*, exhibited reduced expression, suggesting altered mitochondrial–nuclear metabolic coordination rather than activation of a canonical mitochondrial stress response.

In stationary phase, the transcriptional landscape of *npr2Δ* cells shifted markedly. Gene ontology analysis revealed enrichment of transport- and maintenance-associated processes, including nucleobase and nucleoside transport, xenobiotic transport, glycolysis, ATP generation, and nucleotide phosphorylation, with the majority of associated genes exhibiting reduced expression **(Figures 2D–2F and Extended Data 2)**. Gene-level analysis confirmed broad downregulation of central carbon metabolism, energy production, and nucleotide metabolism genes, accompanied by selective preservation of transport-related functions **(Figure S3B)**. This pattern is consistent with global transcriptional dampening and stabilization of a low-flux, maintenance-oriented metabolic configuration during the non-proliferative state.

Together, these data indicate that SEACIT loss does not impose a uniform transcriptional effect but instead establishes a state-dependent metabolic program, characterized by anabolic compensation during growth and transition into a low-flux maintenance configuration during stationary phase (**Extended Data 3)**. This transcriptional remodeling provides a mechanistic basis for the growth–survival uncoupling observed in SEACIT mutants **(Figure 1)**.

### Mitochondrial stress signaling modulates SEACIT-dependent metabolic states and survival

The state-dependent transcriptional remodeling observed in SEACIT-deficient cells **(Figure 2)** revealed a low-flux, maintenance-oriented metabolic configuration during stationary phase without activation of canonical mitochondrial stress programs. We therefore asked whether the stability of this SEACIT-dependent metabolic state is influenced by mitochondrial stress signal transmission.

In yeast, mitochondrial stress signals are transmitted to the nucleus primarily through the retrograde (RTG) pathway **(Figure 3A)**. Deletion of *RTG1*, *RTG2*, or *RTG3* significantly increased long-term survival in SD medium **(Figure S4A)**, with *rtg1Δ* exhibiting a stronger effect than SEACIT mutants **(Figure 3B)**, indicating that mitochondrial stress signaling itself can limit survival under nutrient-limited conditions.

**Figure 3.**
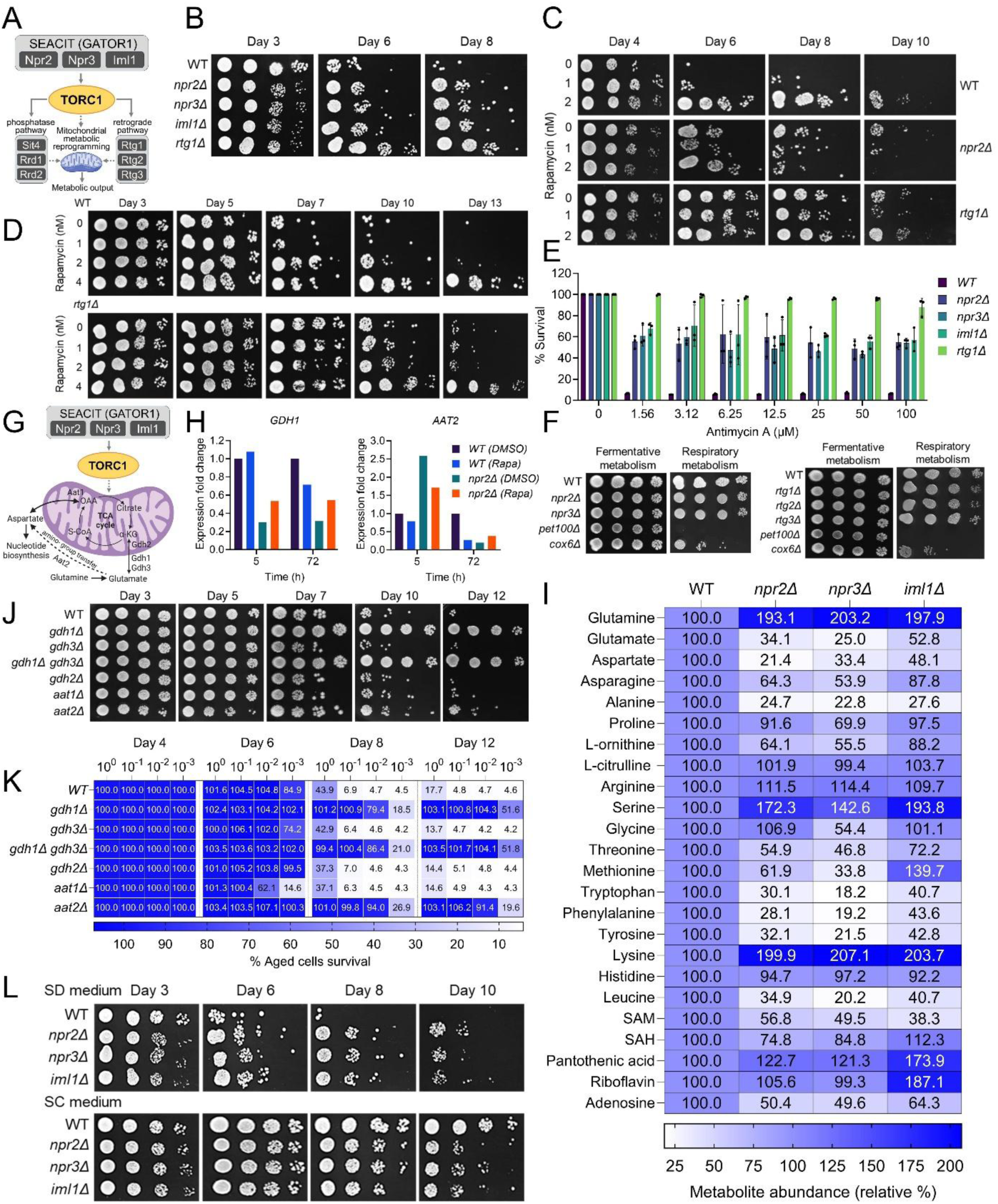
Mitochondrial stress signaling and nitrogen metabolism modulate the SEACIT-dependent long-term survival state. (A) Schematic representation of SEACIT–TORC1 regulation and downstream effector branches linking nutrient sensing to mitochondrial metabolic signaling in *Saccharomyces cerevisiae*, including phosphatase-dependent and retrograde (RTG) pathways. (B) Chronological survival assessed by spot dilution assays for wild-type, *npr2Δ*, *npr3Δ*, *iml1Δ*, and *rtg1Δ* cells at the indicated days during stationary phase. (C) Chronological survival of wild-type, *npr2Δ*, and *rtg1Δ* cells treated with increasing concentrations of rapamycin, monitored by spot dilution assays at the indicated days. (D) Rapamycin dose–response survival analysis of wild-type and *npr2Δ* cells during stationary phase, assessed by spot dilution assays. (E) Quantification of cell survival following treatment with increasing concentrations of antimycin A. Survival at day 3 is expressed as a percentage relative to untreated controls for wild-type and indicated mutants (*npr2Δ*, *npr3Δ*, *iml1Δ*, *rtg1Δ*). Data represent mean ± SD (n = 3). (F) Growth of SEACIT mutants (*npr2Δ*, *npr3Δ*), RTG mutants (*rtg1Δ*, *rtg2Δ, rtg3Δ*), and mitochondrial-deficient (*pet100*, *cox6Δ*) strains under fermentative (YP + glucose) and respiratory (YP + glycerol) conditions, assessed by spot dilution assays. (G) Schematic summary illustrating SEACIT–TORC1 control of mitochondrial-linked nitrogen metabolism, including glutamate–glutamine interconversion, aspartate transamination, and nucleotide-associated metabolic outputs. (H) Relative expression of *GDH1* and *AAT2* in wild-type and *npr2Δ* cells treated with DMSO or rapamycin, measured during exponential growth (5 h) and stationary phase (72 h). (I) Heatmap showing relative intracellular metabolite abundance in *npr2Δ*, *npr3Δ*, and *iml1Δ* cells compared with wild-type (set to 100%). Metabolites associated with amino acid, nitrogen, and one-carbon metabolism are shown. (J) Survival of wild-type and indicated amino acid metabolism mutants (*gdh1Δ*, *gdh3Δ*, *gdh1Δ gdh3Δ*, *gdh2Δ*, *aat1Δ*, *aat2Δ*) assessed by spot dilution assays at the indicated days. (K) Quantification of survival for strains shown in (J), expressed as percentage relative to wild-type, measured by outgrowth-based viability assays in YPD medium. (L) Spot dilution–based survival analysis of wild-type, *npr2Δ*, *npr3Δ*, and *iml1Δ* cells aged in synthetic defined (SD) or synthetic complete (SC) medium, demonstrating nutrient-context dependence of SEACIT-associated survival.

We next examined whether TORC1 inhibition perturbs the SEACIT-associated metabolic state through mitochondrial signaling. During exponential growth, rapamycin exerted only modest effects on mitochondrial TCA and anaplerotic gene expression in both wild-type and *npr2Δ* cells **(Figure S4B)**. In contrast, during stationary phase, rapamycin induced genotype-dependent remodeling of mitochondrial metabolism: wild-type cells showed broad repression of oxidative and anaplerotic genes, whereas *npr2Δ* cells exhibited selective restoration of oxidative components (*ACO1*, *IDH1*) together with reduced expression of anaplerotic *PYC2* and downstream TCA component *KGD1* **(Figure S4C)**. Consistent with destabilization of the SEACIT-associated metabolic configuration, rapamycin reduced survival in *npr2Δ* cells but not in *rtg1Δ* cells **(Figures 3C and 3D)**.

To directly test whether mitochondrial respiratory stress similarly depends on stress signal transmission, we treated cells with antimycin A (AMA) ^47^. AMA markedly reduced survival in wild-type cells, whereas SEACIT mutants were largely resistant **(Figure 3E)**. Strikingly, *rtg1Δ* cells were completely resistant to AMA-induced survival loss **(Figure 3E)**. Deletion of *SIT4*, *RRD1*, or *RRD2* also conferred resistance to AMA **(Figure S3E)**, supporting a role for mitochondrial stress signaling downstream of TORC1 in modulating survival.

To distinguish regulated metabolic maintenance from intrinsic mitochondrial dysfunction, we compared *npr2Δ* cells with respiratory-deficient *cox6Δ* cells, which lack a core subunit of mitochondrial complex IV ^48^. *cox6Δ* cells exhibited transcriptional signatures of acute mitochondrial dysfunction during growth and stress-associated repression of ribosome biogenesis and translation during stationary phase, whereas *npr2Δ* cells selectively induced amino acid biosynthesis programs and maintained metabolic and transport-related gene expression **(Figures S4F–S4I and Extended Data 4 and 5)**. Consistently, deletion of *COX6* or *PET100,* a mitochondrial assembly factor required for cytochrome c oxidase function ^48^, resulted in markedly shortened survival **(Figure S4J)** ^49^.

Finally, assessment of growth under fermentative (glucose) and respiratory (glycerol) conditions showed that SEACIT and RTG mutants were capable of growth under both conditions, indicating preserved mitochondrial respiratory competence, whereas *pet100Δ* and *cox6Δ* cells were defective for growth on glycerol **(Figure 3F)**.

Together, these results demonstrate that SEACIT-dependent survival reflects stabilization of a regulated low-flux metabolic state supported by intact mitochondrial function, whereas mitochondrial stress signaling, rather than mitochondrial dysfunction per se, acts as a conditional destabilizer of this state under nutrient-limited conditions.

### SEACIT loss rewires nitrogen and nucleotide metabolism under nutrient limitation

The stabilization of a SEACIT-dependent low-flux metabolic state prompted us to identify the metabolic pathways that define this configuration. Transcriptomic analyses revealed prominent remodeling of nitrogen-associated programs in SEACIT-deficient cells **(Figure 2)**, leading us to examine nitrogen and nucleotide metabolism in detail.

Across growth and stationary phases, SEACIT loss induced coordinated but selective rewiring of amino acid biosynthesis, nitrogen transfer, and nucleoside transport pathways **(Figure 2)**. *GDH1* and *GDH3* were consistently downregulated, whereas *GDH2* was upregulated **(Figures 3G, S5A and Table S1)** ^50,51^. Genes in the aspartate/asparagine branch (*AAT1*, *AAT2*) ^51–53^ were elevated during growth but reduced during stationary phase, while amino acid transporters ^54^ showed divergent regulation, with *GAP1* remaining elevated and *DIP5* declining at later stages **(Figures 3G, S5A and Table S1)**. Notably, rapamycin reversed several of these expression patterns in *npr2Δ* cells, particularly restoring *GDH1* and *AAT2* expression at 72 h **(Figures 3H and S5B–S5D)**, indicating TORC1-dependent modulation of nitrogen assimilation.

Targeted metabolomics at 72 h revealed pronounced accumulation of glutamine in *npr2Δ*, *npr3Δ*, and *iml1Δ* cells, accompanied by reduced glutamate and downstream amino acids, as well as decreased adenosine, a purine-derived nucleoside **(Figure 3I)**. Similar metabolic signatures were observed in *rtg1Δ* and *sit4Δ* cells **(Figure S5E)**, linking nitrogen and nucleotide remodeling to survival-promoting metabolic states downstream of TORC1.

Genetic perturbation of nitrogen assimilation enzymes revealed pathway specificity. Deletion of *GDH1*, but not *GDH2* or *GDH3*, significantly enhanced long-term survival, while *aat2Δ*, but not *aat1Δ*, produced a similar effect **(Figures 3J, 3K, S5F and S5G)**. These results indicate that selective nodes within nitrogen assimilation, rather than global amino acid biosynthesis, shape survival outcomes.

Finally, the SEACIT-dependent metabolic configuration was highly sensitive to nutrient context. When cells were aged in nutrient-replete medium supplemented with amino acids and nucleobases, SEACIT and RTG mutants no longer exhibited enhanced survival and instead displayed shortened survival despite persistently reduced growth **(Figures 3L and S5H–S5K)**. These findings establish nutrient limitation as a critical requirement for stabilization of the SEACIT-dependent low-flux metabolic state.

### Methotrexate phenocopies the SEACIT–TORC1 metabolic state through nucleotide restraint

The SEACIT-dependent metabolic state identified above is characterized by reduced anabolic flux, selective remodeling of nitrogen utilization, and suppression of nucleotide metabolism under nutrient-limited conditions. To test whether direct restraint of nucleotide biosynthesis is sufficient to engage this metabolic configuration, we examined methotrexate (MTX), an antifolate that limits one-carbon metabolism and purine and pyrimidine synthesis ^55^.

MTX caused a dose-dependent reduction in early growth while enhancing long-term survival **(Figures 4A and 4B)**. SEACIT mutants exhibited resistance to MTX-induced growth inhibition, closely mirroring their response to rapamycin **(Figures 4C, S6A and S6B)**. Similar resistance was observed in RTG mutants **(Figure S6C)**, indicating that nucleotide restraint converges on the same low-flux metabolic state defined genetically by SEACIT and mitochondrial stress signaling.

**Figure 4.**
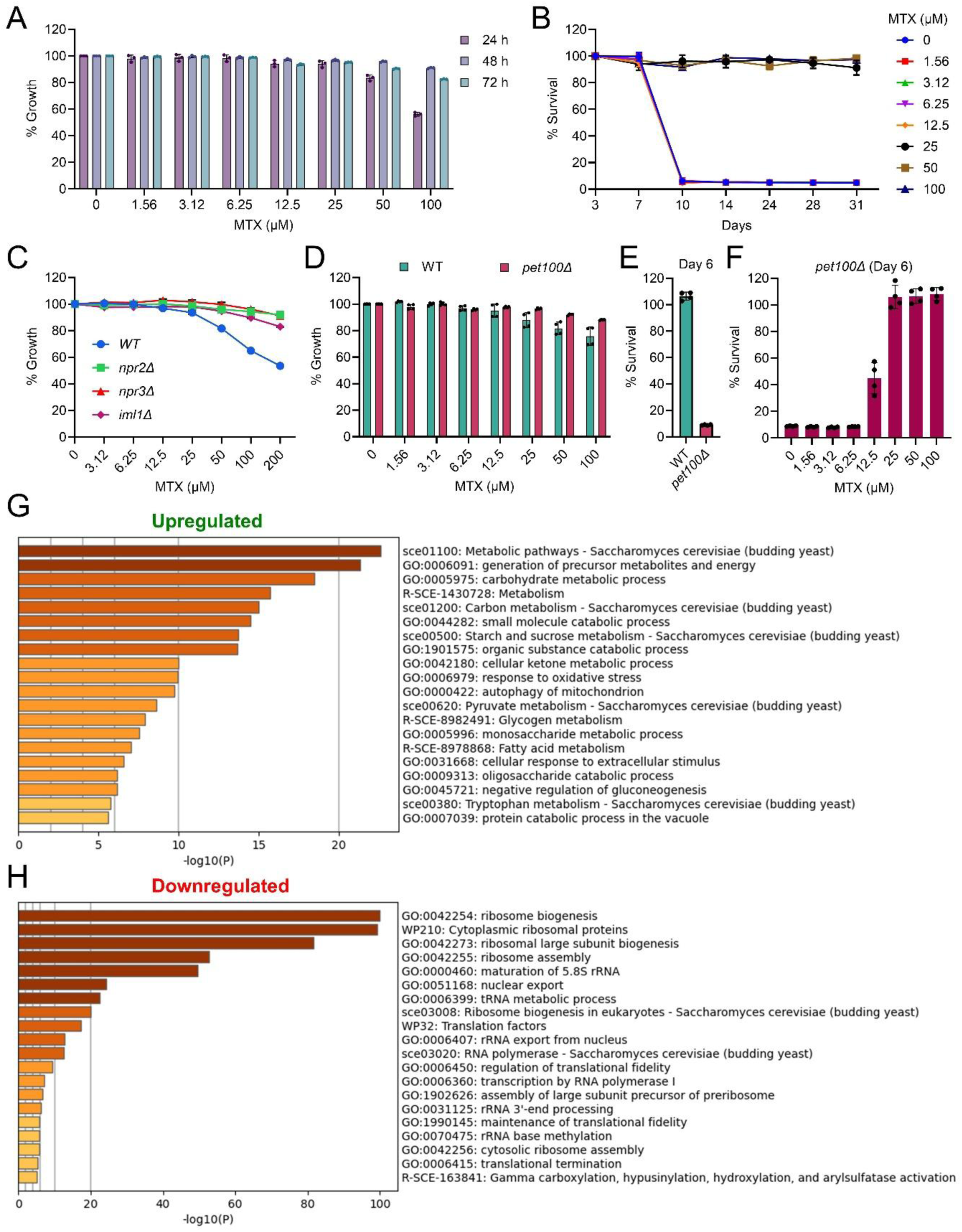
Methotrexate phenocopies the SEACIT–TORC1 low-flux metabolic state through nucleotide restraint. (A) Relative growth of wild-type cells treated with increasing concentrations of methotrexate (MTX), measured at 24 h, 48 h, and 72 h in synthetic defined (SD) medium. Growth is expressed as a percentage relative to untreated controls (mean ± SD, n = 3). (B) Survival analysis of wild-type cells treated with MTX, showing dose-dependent effects on aged-cell survival over time (mean ± SD, n = 3). (C) Relative growth of wild-type and SEACIT mutant strains (*npr2Δ*, *npr3Δ*, *iml1Δ*) treated with increasing concentrations of MTX, measured at 24 h (mean ± SD, n = 2). (D) Relative growth of wild-type and mitochondrial-deficient *pet100Δ* cells treated with increasing concentrations of MTX, measured at 48 h (mean ± SD, n = 4). (E) Comparison of survival for wild-type and *pet100Δ* cells on day 6 (mean ± SD, n = 4). (F) Dose-dependent effects of MTX on survival in *pet100Δ* cells on day 6 (mean ± SD, n = 4). (G–H) Heatmap of Gene Ontology (GO) enrichment for genes upregulated (F) and downregulated (G) following MTX treatment of logarithmic-phase prototrophic wild-type cells.

We next asked whether nucleotide restraint could modify survival outcomes associated with mitochondrial dysfunction. MTX significantly suppressed the shortened survival of *pet100Δ* cells in a dose-dependent manner **(Figures 4D–4F)**, demonstrating that inhibition of nucleotide biosynthesis can mitigate mitochondrial-associated survival decline under nutrient-limited conditions.

To define the transcriptional basis of this response, we performed transcriptomic profiling following acute MTX treatment. MTX induced coordinated repression of ribosome biogenesis and translation-associated processes, accompanied by induction of metabolic and stress-adaptive pathways **(Figures 4F, 4G; Table S1; Extended Data 6 and 7)**. Acute treatments were profiled at early time points to capture primary engagement of the metabolic network, enabling comparison with genetically defined metabolic states. Comparative analysis revealed extensive overlap between MTX and rapamycin responsive gene sets at both upregulated and downregulated levels **(Figures S6D–S6K; Extended Data 8 and 9)**, indicating that nucleotide biosynthetic inhibition phenocopies TORC1 inhibition at the level of global transcriptional output.

Together, these results establish nucleotide restraint as a sufficient intervention to engage the SEACIT-associated low-flux metabolic state, demonstrating that targeted limitation of anabolic input can recapitulate this configuration under nutrient-limited conditions.

### Glycolytic restraint uncovers mitochondrial competence–dependent survival

We next asked whether attenuation of other central anabolic inputs similarly engages the SEACIT-associated metabolic state. Because glycolytic flux is a major determinant of cellular energy balance and mitochondrial demand, we examined the effects of early glycolytic restraint using mechanistically related hexose analogs **(Figure S7A)**.

Treatment with 2,5-anhydromannitol (2,5-AM), a fructose analog phosphorylated by hexokinase but not further metabolized through glycolysis, imposed a moderate restraint on glycolytic flux ^56–58^. In wild-type cells grown under nutrient-limited conditions, 2,5-AM caused a modest reduction in early growth while enhancing long-term survival **(Figures S7B and S7C)**. Consistent with their responses to rapamycin, *npr2Δ* cells were resistant to 2,5-AM–induced growth inhibition **(Figure S7B)**. At low concentrations, however, 2,5-AM reduced survival in *npr2Δ* cells **(Figure S7C)**, phenocopying the paradoxical effect observed with low-dose rapamycin. This survival reduction was absent in *rtg1Δ* cells **(Figure S7D)**, indicating that mitochondrial stress signal transmission is required for glycolytic restraint to destabilize the SEACIT-associated metabolic state.

To define the molecular programs engaged by moderate glycolytic restraint, we performed transcriptomic profiling following acute 2,5-AM treatment. 2,5-AM elicited a transcriptional response highly similar to those induced by rapamycin and methotrexate, characterized by coordinated repression of ribosome biogenesis and translation-associated processes together with induction of metabolic and stress-adaptive programs **(Figures 5A–5H, S8A-S8H and S9A–S9D; Table S1 and Extended Data 10-13)**. These data place moderate glycolytic attenuation within the same TORC1-linked transcriptional response space as the SEACIT-dependent metabolic state.

**Figure 5.**
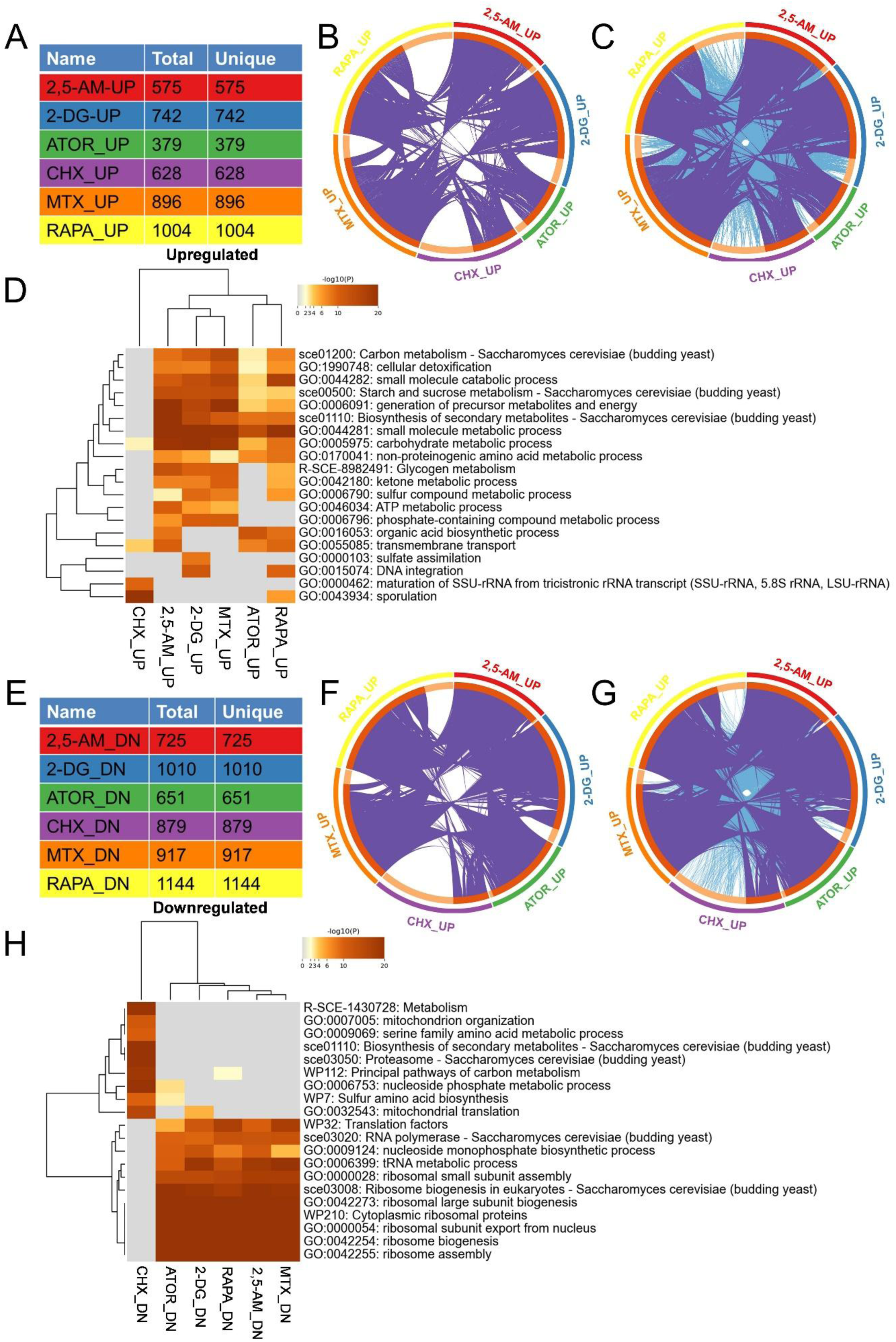
Convergent and divergent transcriptional responses across metabolic perturbations. (A) Summary table showing the total number of genes upregulated following acute treatment with rapamycin (RAPA), methotrexate (MTX), 2,5-anhydromannitol (2,5-AM), 2-deoxyglucose (2-DG), atorvastatin (ATOR), and cycloheximide (CHX). (B) Circular plot showing gene-level overlap among genes upregulated by RAPA, MTX, 2,5-AM, 2-DG, ATOR, and CHX. (C) Circular plot showing functional overlap of Gene Ontology (GO) enrichments associated with upregulated gene sets across treatments. (D) Heatmap of GO enrichment for biological processes associated with upregulated genes across treatments. Color scale represents –log10(P value). (E) Summary table showing the total and unique numbers of genes downregulated following treatment with RAPA, MTX, 2,5-AM, 2-DG, ATOR, and CHX. (F) Circular plot showing gene-level overlap among downregulated genes across treatments. (G) Circular plot showing functional overlap of GO enrichments associated with downregulated gene sets. (H) Heatmap of GO enrichment for biological processes associated with downregulated genes. Color scale represents –log10(P value).

We next compared 2,5-AM with 2-deoxyglucose (2-DG), a glucose analog that more potently blocks early glycolytic flux ^59^. As observed for rapamycin, *npr2Δ* cells were resistant to 2-DG-induced growth inhibition **(Figures S7E and S7F)**. Although 2-DG induced overlapping transcriptional signatures **(Figures 5A–5H, S10A–S10D, S11A–S11D and S12A–S12D; Table S1 and Extended Data 10-13)**, it caused substantially stronger growth inhibition **(Figures S7G and S7H)** and pronounced repression of mitochondrial oxidative phosphorylation genes **(Figures 5H, S12E and S12F)**, indicating a more severe disruption of metabolic homeostasis.

Consistent with these transcriptional differences, 2-DG enhanced survival in wild-type cells but further reduced survival in mitochondrial-deficient *pet100Δ* cells, whereas 2,5-AM and rapamycin suppressed *pet100Δ*-associated survival loss **(Figures 7I–7L)**. These results indicate that glycolytic restraint promotes survival only when mitochondrial respiratory capacity is preserved, revealing mitochondrial competence as a critical determinant of survival outcomes.

Together, these findings demonstrate that moderate attenuation of glycolytic flux engages the SEACIT-associated low-flux metabolic state in a mitochondrial competence–dependent manner, whereas more severe blockade of glycolysis destabilizes this state when mitochondrial function is compromised.

### Sterol restriction converges on the SEACIT–TORC1 low-flux metabolic state

The convergence of nucleotide and glycolytic restraint on a SEACIT-associated low-flux metabolic state raised the question of whether limitation of additional anabolic pathways similarly engages this configuration. We therefore examined sterol biosynthesis, a major anabolic output downstream of acetyl-CoA metabolism, using atorvastatin, an inhibitor of HMG-CoA reductase ^60,61^.

Under nutrient-limited conditions, atorvastatin modestly reduced early growth while enhancing long-term survival in wild-type cells **(Figures S13A and S13B)**. In contrast, atorvastatin further reduced survival in mitochondrial-deficient *pet100Δ* cells **(Figures S13A and S13D)**, indicating that sterol restriction promotes survival only when mitochondrial respiratory capacity is preserved. This mitochondrial dependence parallels that observed under strong glycolytic restraint and supports a shared gating principle across distinct anabolic inputs.

Growth assays revealed that SEACIT mutants (*npr2Δ*, *npr3Δ*, *iml1Δ*) were resistant to atorvastatin-induced growth inhibition, closely paralleling their responses to rapamycin, methotrexate, 2,5-anhydromannitol, and 2-deoxyglucose **(Figure S13E)**. In contrast, RTG mutants displayed growth responses similar to wild-type cells **(Figure S13F)**, indicating that sterol restraint converges on the SEACIT-defined metabolic state at the level of anabolic growth control, independently of mitochondrial stress signal transmission.

Transcriptomic profiling following atorvastatin treatment revealed strong overlap with transcriptional responses elicited by rapamycin, methotrexate, and glycolytic restraint, including coordinated repression of ribosome biogenesis and other anabolic processes together with induction of metabolic and stress-adaptive pathways **(Figures 5A–5H, S13G and S13H; Table S1 and Extended Data 12-15)**. These data indicate that sterol restriction engages the same TORC1-linked low-flux metabolic configuration defined by genetic and pharmacological perturbations upstream of anabolic metabolism.

### Translation inhibition engages a survival state distinct from metabolic restraint

We next examined whether direct inhibition of protein synthesis engages the SEACIT-associated metabolic state. To this end, we analyzed the effects of cycloheximide (CHX), an inhibitor of translational elongation ^62^.

CHX treatment reduced growth and enhanced long-term survival, similar to rapamycin **(Figures S14A-S14D)**. However, in contrast to metabolic restraint, SEACIT and RTG mutants did not exhibit resistance to CHX-induced growth inhibition, indicating that translation inhibition does not converge on the SEACIT-defined growth control logic **(Figures S14E and S14F)**. Transcriptomic analysis further revealed a qualitatively distinct response. While translation-associated pathways were strongly repressed, CHX failed to induce the coordinated metabolic remodeling shared by SEACIT-associated interventions and instead exhibited strong enrichment of sporulation and developmental associated gene ontology terms **(Figures 5A–5H; Table S1; Extended Data 12, 13 16 and 17)**.

To place these differences in the context of nutrient signaling, we examined TORC1 activity under defined starvation and refeeding conditions. In prototrophic CEN.PK cells, glucose addition following water starvation failed to activate TORC1, as assessed by Sch9 phosphorylation, indicating that glucose availability alone is insufficient to stimulate TORC1 under these conditions **(Figure S15A)**. In contrast, glucose addition following CHX treatment resulted in robust TORC1 activation **(Figure S15A)**.

Targeted metabolomic profiling showed that translation inhibition was associated with accumulation of a subset of amino acids, most prominently branched-chain amino acids including leucine, as well as aromatic amino acids such as tyrosine **(Figures S15B, S15D, and S15E)**. In contrast, glutamate levels did not increase **(Figures S15B and S15F)**, indicating that amino acid accumulation was not a uniform consequence of altered nitrogen metabolism. Adenosine levels increased under both water starvation and CHX treatment and were reduced upon glucose addition in both contexts **(Figures S15B and S15G)**. However, despite similar adenosine dynamics, glucose-induced TORC1 activation occurred only in the CHX-treated condition, indicating that adenosine reduction alone is insufficient to account for TORC1 activation.

Consistent with this nutrient dependence, glucose addition in yeast nitrogen base (YNB) medium activated TORC1, albeit less robustly than in CHX-treated cells, highlighting the importance of nitrogen context in TORC1 responsiveness **(Figure S15C)**. Together, these results show that translation inhibition alters cellular growth, survival, transcriptional programs, and TORC1 signaling in a manner that is distinct from the SEACIT–TORC1 low-flux metabolic state engaged by metabolic restraint.

### Context-dependent and conserved responses to metabolic perturbations in human cells

To determine whether metabolic perturbations identified in yeast produce comparable regulatory responses in mammalian systems, we analyzed Connectivity Map (CMAP) transcriptomic datasets for compounds targeting distinct metabolic processes, including TORC1 inhibition (rapamycin (sirolimus) and temsirolimus), nucleotide metabolism (methotrexate), sterol biosynthesis (atorvastatin), and translation inhibition (cycloheximide) ^63,64^. We additionally queried CMAP for transcriptomic profiles of 2-deoxyglucose (2-DG) and 2,5-anhydromannitol (2,5-AM); however, RNA-sequencing datasets for these perturbations were not available under the standardized conditions used for cross-compound comparisons, precluding their inclusion in the transcriptomic clustering analysis. Comparison of perturbation strength across treatment durations and drug concentrations indicated that 24 h at 10 µM produced the most robust transcriptional responses, and this condition was therefore selected for subsequent analyses **(Figures S16-S21)**.

We first examined transcriptional responses to individual perturbations across multiple cell lines. For each compound, comparison of the top 20 up- and downregulated genes revealed minimal overlap between cell types, indicating that gene-level responses to metabolic perturbations are strongly context dependent **(Figures S22–S31).**

We next performed a complementary analysis comparing multiple perturbations within individual cell lines. To enable direct comparison, we restricted the analysis to cell lines with at least three overlapping perturbations in the CMAP dataset, resulting in four cell lines representing distinct tissue origins: A375 (melanoma), A549 (lung adenocarcinoma), MCF7 (breast cancer), and VCAP (prostate cancer) **(Figure 6A)**. Among these, MCF7 exhibited the most comprehensive compound coverage and was therefore used as a reference for cross-perturbation comparisons. Analysis of the top 200 up- and downregulated genes per perturbation revealed substantial overlap in functional pathway enrichment within individual cell types despite limited gene-level concordance **(Figures 6B–6E and S32–S39; Table S2; Extended Data 18–25)**.

**Figure 6.**
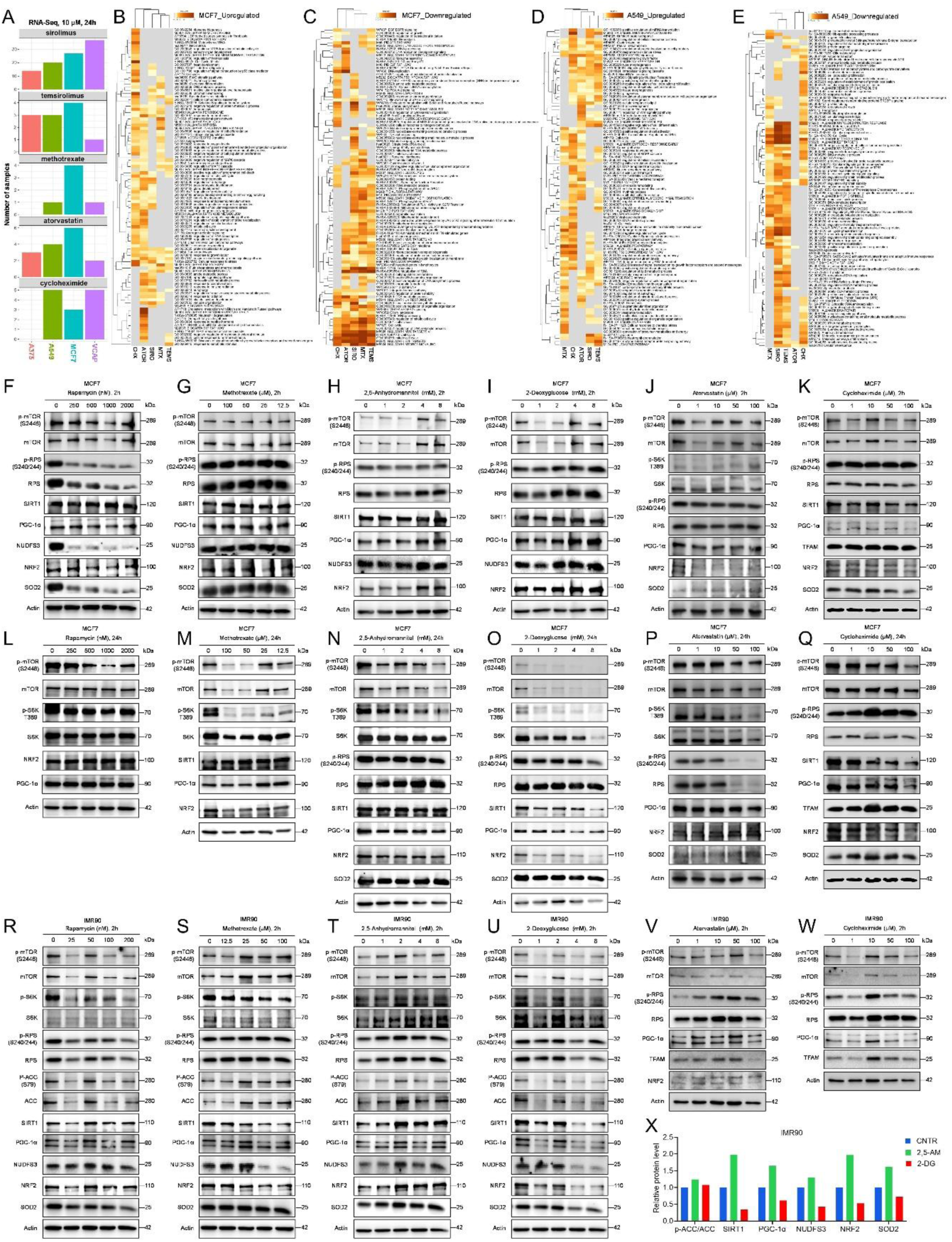
Context-dependent transcriptional and signaling responses to metabolic perturbations in human cells. (A) Summary of transcriptional perturbation strength across compounds and cell lines from Connectivity Map (CMAP) analysis (10 μM, 24 h), showing relative numbers of responsive genes. (B–C) Heatmaps of Gene Ontology (GO) enrichment for biological processes associated with genes upregulated (B) or downregulated (C) by metabolic perturbations in MCF7 cells. (D–E) Heatmaps of GO enrichment for biological processes associated with genes upregulated (D) or downregulated (E) in A549 cells. Color scale represents −log10(P value). (F–K) Immunoblot analysis of mTORC1 signaling, mitochondrial, and stress-associated markers in MCF7 cells following acute (2 h) treatment with rapamycin (F), methotrexate (G), 2,5-anhydromannitol (H), 2-deoxyglucose (I), atorvastatin (J), and cycloheximide (K). (L–Q) Immunoblot analysis of mTORC1 signaling, mitochondrial, and stress-associated markers in MCF7 cells following prolonged (24 h) treatment with the indicated perturbations. (R–W) Immunoblot analysis in IMR90 human lung fibroblasts following acute (2 h) treatment with the indicated perturbations. (X) Quantification of selected mitochondrial and stress-associated protein levels in IMR90 cells under glycolytic perturbations (2,5-anhydromannitol and 2-deoxyglucose), normalized to control.

Across cell lines, hierarchical clustering revealed that sirolimus and temsirolimus consistently grouped together, with methotrexate showing substantial overlap, whereas atorvastatin displayed partial convergence and cycloheximide segregated as a distinct cluster **(Figures 6B–6E and S32–S37)**. Despite targeting different upstream processes, sirolimus, temsirolimus, and methotrexate induced a shared transcriptional state characterized by suppression of MYC–E2F–driven proliferative programs alongside coordinated activation of stress-responsive and metabolic pathways, including mitochondrial-associated processes. In contrast, cycloheximide produced a distinct transcriptional profile marked by translational inhibition without comparable engagement of mitochondrial or stress pathways.

While this overall regulatory architecture was conserved across cell types, differences were observed in the relative enrichment of specific pathways, including enhanced receptor and KRAS-associated signaling in A549 cells, hormone-related signaling in MCF7 cells, and lipid and sterol-associated metabolic programs in VCAP cells **(Figures 6B–6E and S32–S37)**. Together, these findings indicate that distinct perturbations converge on partially conserved cellular states defined by coordinated coupling between mTORC1-associated growth suppression and engagement of mitochondrial and stress pathways, rather than representing uniform outcomes of growth inhibition. Similar relationships were observed across the remaining cell lines **(Figures S38 and S39)**, indicating that while transcriptional responses are highly context-dependent across cell types (**Figures S22–S31**), distinct perturbations converge on similar functional states within individual cellular contexts.

### Shared mTORC1 suppression reveals divergent mitochondrial and stress responses

To assess whether the transcriptional responses identified in CMAP extend beyond the analyzed models and across distinct cellular contexts, we performed experimental validation in MCF7 cells together with IMR90 primary human lung fibroblasts and HEPG2 (hepatocellular carcinoma) cells.

To experimentally evaluate signaling responses, we tested rapamycin, methotrexate, atorvastatin, cycloheximide, 2-deoxyglucose, and 2,5-anhydromannitol, and assessed mTORC1 activity together with mitochondrial- and stress-associated pathways by immunoblotting across acute **(Figures 6F–6K)** and prolonged **(Figures 6L–6Q)** treatments in MCF7 cells and acute treatment in IMR90 cells **(Figures 6R–6W)**. Across these systems, all perturbations except cycloheximide reduced mTORC1 signaling, although the extent of inhibition varied between compounds. In contrast, cycloheximide exhibited a distinct signaling profile and maintained or stabilized mTORC1 activity despite suppressing translation.

These perturbations were accompanied by differential regulation of mitochondrial- and stress-associated markers. In untreated MCF7 cells, the mitochondrial regulator PGC-1α increased over time **(Figures 6F–6Q)**, indicating a baseline enhancement of mitochondrial programs during prolonged culture. Perturbations therefore modulate this trajectory rather than acting on a static baseline.

Glycolytic entry-point perturbations exhibited time- and context-dependent mitochondrial responses. In MCF7 cells, 2-deoxyglucose increased PGC-1α at early time points (2 h), whereas 2,5-anhydromannitol reduced PGC-1α under the same conditions; however, both perturbations were associated with reduced mitochondrial-associated markers upon prolonged treatment (24 h) **(Figures 6H**, 6I**, 6N and 6O)**. In contrast, in IMR90 cells, where only acute responses were assessed, these perturbations diverged, with 2-deoxyglucose associated with reduced mitochondrial components, whereas 2,5-anhydromannitol maintained or increased selected markers **(Figures 6T, 6U and 6X)**. A similar divergence was observed in yeast, indicating that this differential mitochondrial engagement is conserved across species.

Notably, rapamycin decreased mitochondrial-associated markers in MCF7 cells at both time points **(Figures 6F and 6L)**, whereas in IMR90 cells it increased mitochondrial markers under acute treatment **(Figure 6R)**, indicating cell type–dependent differences in mitochondrial responses downstream of TORC1 inhibition. Consistent with these findings, acute treatment in HEPG2 cells showed that rapamycin reduced mTORC1 signaling while increasing mitochondrial- and stress-associated markers, whereas cycloheximide had minimal effects on these pathways and maintained mTORC1 activity **(Figures S40A–S40E)**.

Together, these findings demonstrate that diverse metabolic perturbations converge on mTORC1-linked suppression of proliferative programs while engaging distinct mitochondrial-and stress-associated responses, uncoupling growth inhibition from a single, uniform cellular outcome.

### Nutrient availability determines cellular responses to metabolic perturbations in human cells

To determine whether nutrient availability influences cellular responses to metabolic perturbations, we evaluated cell viability under nutrient-rich (NR; 10% FBS) and nutrient-limited (NL; 1% FBS) conditions across multiple human cell lines, including MCF7 (breast cancer), A549 (lung adenocarcinoma), HEPG2 (hepatocellular carcinoma), SH-SY5Y (neuroblastoma), and IMR90 (primary lung fibroblasts). SH-SY5Y cells were included due to their sensitivity to metabolic stress. Across these models, metabolic perturbations produced distinct viability outcomes depending on both compound and cellular context, with differential sensitivity observed between NR and NL conditions, indicating that nutrient availability strongly shapes cellular responses to metabolic inputs **(Figures 7A–7E)**.

**Figure 7.**
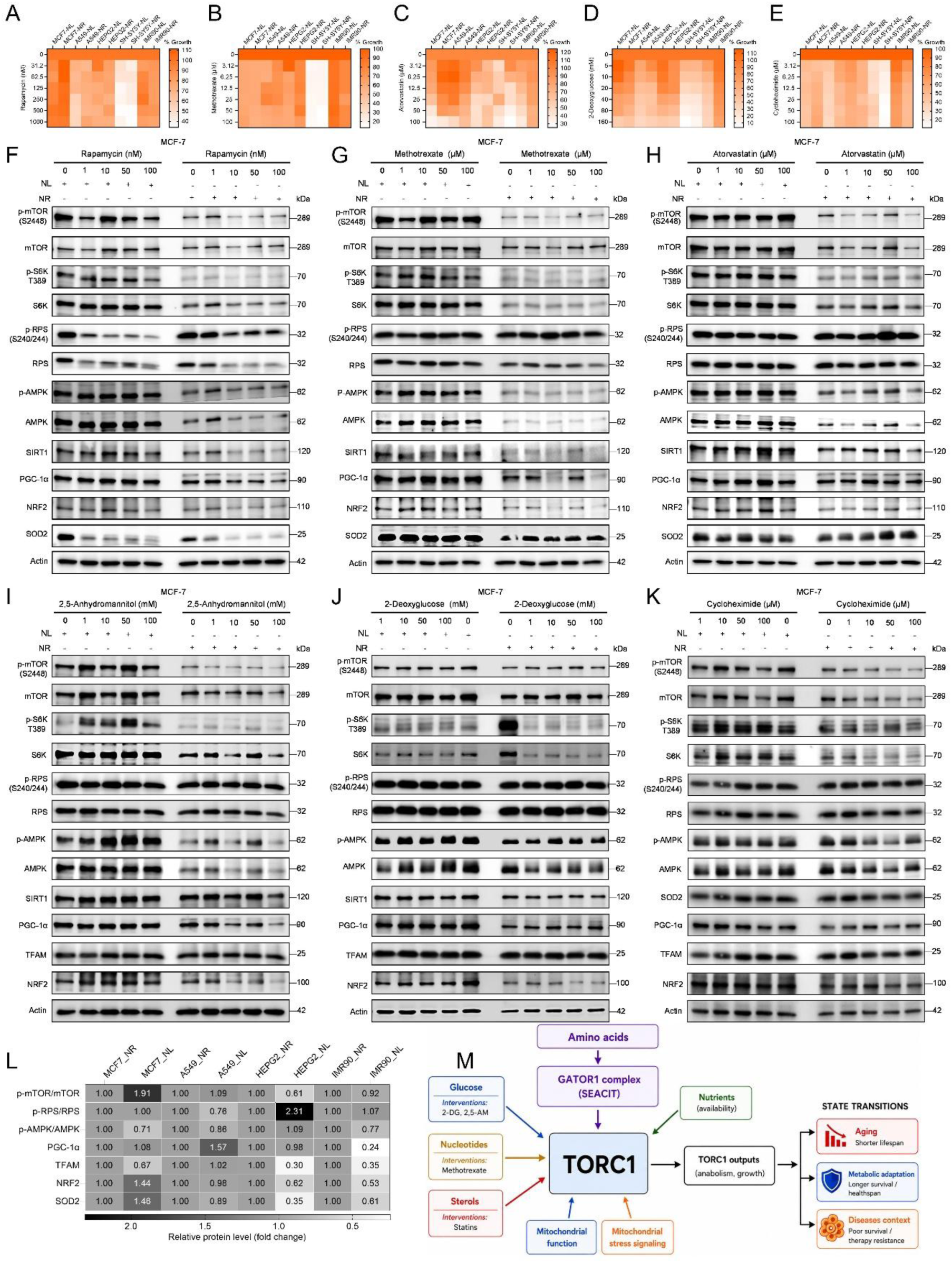
Nutrient availability modulates cellular responses to metabolic perturbations in cancer and primary human cells. (A–E) Cell viability analysis under nutrient-rich (NR; 10% FBS) and nutrient-limited (NL; 1% FBS) conditions in MCF7 (breast cancer), A549 (lung adenocarcinoma), HEPG2 (hepatocellular carcinoma), SH-SY5Y (neuroblastoma), and IMR90 (primary human lung fibroblasts) following treatment with rapamycin (A), methotrexate (B), atorvastatin (C), 2-deoxyglucose (D), and cycloheximide (E). Heatmaps were generated from at least two independent experiments, each performed with at least two biological replicates. Color scale represents relative cell viability. (F–K) Immunoblot analysis of mTORC1 signaling, metabolic, mitochondrial, and stress-associated markers in MCF7 breast cancer cells under NR and NL conditions following treatment with rapamycin (F), methotrexate (G), atorvastatin (H), 2,5-anhydromannitol (I), 2-deoxyglucose (J), and cycloheximide (K). (L) Heatmap summarizing baseline signaling differences between NR and NL conditions across MCF7 (breast cancer), A549 (lung adenocarcinoma), HEPG2 (hepatocellular carcinoma), and IMR90 (primary lung fibroblasts) cells, generated from untreated control samples. Protein levels are shown as fold change relative to NR conditions. (M) Model illustrating that nutrient availability establishes distinct metabolic states that determine cellular responses to metabolic perturbations through TORC1-, mitochondrial-, and stress-associated signaling pathways, resulting in context-dependent cellular outcomes.

To define baseline signaling differences between nutrient conditions, we compared mTORC1, mitochondrial, and stress-associated markers under untreated NR and NL conditions. Nutrient limitation did not induce a uniform response but instead generated cell-type–specific signaling states **(Figures 7F–7L, S41A-S41E, S42A-S42E and S43A-S43E)**. While mTORC1 activity was reduced in HEPG2 and IMR90 cells, it was maintained or increased in MCF7 cells **(Figure 7L)**. Consistent with this, phosphorylation of ribosomal protein S6 (p-RPS) exhibited divergent responses across cell types, indicating differential regulation of translational output. Mitochondrial and stress-associated markers similarly varied, demonstrating that nutrient limitation establishes distinct baseline metabolic configurations rather than a single starvation state.

We next examined how nutrient context modulates responses to metabolic perturbations in MCF7 cells. Although all perturbations except cycloheximide reduced mTORC1 signaling under both NR and NL conditions, downstream responses differed markedly **(Figures 7F-7K)**. Rapamycin and methotrexate produced similar context-dependent effects, with increased SIRT1, PGC-1α, and NRF2 under NL conditions but reduced levels under NR conditions. In contrast, atorvastatin increased these markers under both conditions, indicating a context-independent activation of mitochondrial and stress-associated pathways. Glycolytic perturbations showed more complex behavior: 2,5-anhydromannitol increased AMPK and SIRT1 under both conditions but exhibited nutrient-dependent divergence in mitochondrial and stress responses, whereas 2-deoxyglucose produced more limited and context-restricted effects. Cycloheximide had minimal impact on these pathways.

Similar patterns were observed in additional cell lines (A549, HEPG2, and IMR90), where nutrient availability modulated the extent and direction of mitochondrial and stress-associated responses to metabolic perturbations, although the specific profiles varied between cell types **(Figures S41A-S41E, S42A-S42E and S43A-S43E)**. These data indicate that the influence of nutrient context on cellular response is a generalizable feature rather than a cell line–specific effect.

Together, these findings demonstrate that nutrient availability defines the cellular context in which metabolic perturbations are interpreted. By establishing distinct baseline metabolic states, nutrient conditions determine how mTORC1 suppression is coupled to mitochondrial and stress-associated programs, thereby shaping cellular outcomes.

## DISCUSSION

Cells continuously adapt to fluctuations in nutrient availability by coordinating growth, metabolism, and stress responses through conserved signaling networks centered on TORC1 ^5–15^. Although TORC1 inhibition is widely associated with lifespan extension and stress resistance ^4,14–17,19^, the relationship between reduced TORC1 activity and long-term survival remains incompletely understood. Here, we show that TORC1 attenuation is permissive but not determinative for survival under nutrient limitation. Instead, survival emerges from distinct metabolic states shaped by mitochondrial competence, nutrient availability, and the nature of downstream metabolic remodeling.

A central finding of this study is that the SEACIT complex (Npr2–Npr3–Iml1), despite functioning as a negative regulator of TORC1 ^32,33,44,65,66^, defines a distinct metabolic regime in which reduced proliferative growth is coupled to enhanced long-term survival. SEACIT mutants occupy a unique region of the growth–survival landscape, demonstrating that suppression of proliferation alone is insufficient to predict cellular fate. Rather, survival depends on the specific metabolic configuration established downstream of TORC1-associated signaling networks.

Transcriptomic and metabolic analyses revealed that SEACIT loss promotes dynamic metabolic remodeling across growth states. During proliferative growth, SEACIT mutants retained anabolic and biosynthetic activity, whereas during stationary phase they transitioned into a maintenance-associated state characterized by suppression of central carbon metabolism, energy-consuming biosynthetic pathways, and nucleotide metabolism alongside selective preservation of transport and adaptive functions. These findings suggest that long-term survival is linked not to early growth behavior itself, but to the ability to establish a metabolically restrained state during nutrient limitation.

Our results further identify mitochondrial regulation as a critical determinant of state stability. Although mitochondrial respiratory competence remained preserved in SEACIT mutants, activation of mitochondrial stress signaling through the RTG pathway destabilized the survival-supporting configuration. These findings separate mitochondrial respiratory function from mitochondrial stress signaling and indicate that survival depends not simply on mitochondrial activity, but on how mitochondrial-derived signals are integrated into broader adaptive metabolic programs ^67^. Together, these observations support a model in which mitochondrial competence acts as a gating factor that permits entry into survival-supportive states, whereas excessive mitochondrial stress signaling disrupts their stability.

Nitrogen and nucleotide metabolism emerged as central components of the SEACIT-associated metabolic state. Selective rewiring of nitrogen utilization pathways and reduced nucleotide-associated outputs were consistently observed, and targeted perturbation of these pathways partially recapitulated survival phenotypes ^44,50,52–54^. Importantly, nutrient repletion abolished the survival advantage of SEACIT mutants, indicating that nutrient limitation is required to stabilize this adaptive metabolic configuration. These findings suggest that survival-supportive states are not constitutive properties of reduced TORC1 signaling, but instead emerge specifically under nutrient-constrained conditions.

Pharmacological perturbations further reinforced this framework. Methotrexate phenocopied aspects of the SEACIT-associated state through nucleotide restriction ^17,18,55^, while moderate glycolytic attenuation and sterol perturbation converged on related transcriptional and functional responses ^56–61^. In contrast, severe mitochondrial dysfunction or excessive metabolic disruption impaired survival, indicating that adaptive metabolic restraint differs fundamentally from catastrophic energetic collapse. Notably, translation inhibition promoted survival through a mechanistically distinct route, further demonstrating that growth suppression alone does not define a common cellular state ^38,39,62,68,69^.

Extension of these analyses into mammalian systems revealed a similar principle of context-dependent state remodeling. Although distinct perturbations generated highly variable gene-level responses across cell types, they converged on shared pathway-level adaptations involving mTORC1 signaling, mitochondrial regulation, and stress-associated programs. Moreover, nutrient availability established distinct baseline metabolic states that strongly influenced how perturbations were interpreted. These findings indicate that cellular responses are not fixed outputs of individual signaling pathways, but instead emerge from interactions between nutrient environment, mitochondrial state, and intrinsic metabolic organization

These observations have important implications for aging biology. Aging tissues frequently experience altered nutrient sensing, reduced metabolic flexibility, mitochondrial dysfunction, and impaired stress adaptation ^2,3,20–22^. Our findings suggest that longevity is not simply a consequence of suppressing anabolic signaling, but rather reflects stabilization of adaptive metabolic states that preserve mitochondrial competence while limiting maladaptive stress signaling. This framework may help explain the context-dependent effects of TORC1 inhibition across organisms and physiological conditions, where similar signaling outputs can produce divergent survival outcomes depending on underlying metabolic state.

Our findings may also provide insight into metabolic adaptation in cancer. Tumor cells commonly encounter nutrient-limited and metabolically heterogeneous microenvironments due to poor vascularization and elevated anabolic demand ^20–25^. Under these conditions, survival may depend on the capacity to establish adaptive metabolic states that balance resource conservation with maintenance of mitochondrial function. The observation that mTORC1 suppression alone does not uniformly predict survival outcomes may therefore help explain variability in responses to metabolic and mTOR-targeted therapies across tumor contexts.

More broadly, these principles may extend to persistence-associated cellular states observed during senescence, stress adaptation, and therapeutic resistance ^70–75^. Non-proliferative cells can remain metabolically active and later regain proliferative capacity under favorable conditions. Our findings suggest that long-term persistence may depend on stabilization of adaptive metabolic states rather than proliferation arrest alone.

Together, our findings support a model in which TORC1 functions as part of a broader metabolic state-transition system rather than a linear determinant of survival. Under nutrient limitation, a SEACIT-defined metabolic configuration promotes survival when mitochondrial competence is preserved, whereas nutrient repletion, mitochondrial stress signaling, or excessive metabolic disruption destabilize this state. These results establish a framework in which cellular fate is governed by dynamic metabolic state transitions, providing a unifying perspective for understanding context-dependent outcomes in aging, stress adaptation, and disease **(Figure 7M)**.

## METHODS

### Yeast strains, growth media, and cultivation conditions

All experiments were performed using the prototrophic *Saccharomyces cerevisiae* strain CEN.PK113-7D ^76^. Gene deletion mutants were constructed using a PCR-mediated homologous recombination strategy following established protocols ^77^. Yeast strains were recovered from −80 °C glycerol stocks by streaking onto YPD agar plates (1% yeast extract, 2% peptone, 2% glucose, 2.5% agar) and incubated at 30 °C for 48–72 h. For experimental assays, cells were propagated in synthetic defined (SD) medium containing 6.7 g/L yeast nitrogen base with ammonium sulfate (without amino acids) supplemented with 2% glucose. Synthetic complete (SC) medium was prepared by supplementing SD medium with standard amino acid and nucleobase mixtures ^68,69^.

### Chemical treatments

Small-molecule treatments were performed using stock solutions of rapamycin, methotrexate, atorvastatin and antimycin A prepared in dimethyl sulfoxide (DMSO). The final DMSO concentration was kept at or below 1% (v/v) for experiments. Stock solutions of 2,5-anhydromannitol, 2-deoxyglucose and cycloheximide were prepared in sterile water.

### Chronological aging assay

Cellular survival experiments were performed using the prototrophic *Saccharomyces cerevisiae* strain CEN.PK113-7D cultivated in SD medium, following established procedures with minor adaptations ^49,58,78^. Overnight cultures were grown at 30 °C with shaking (220 rpm) and diluted into fresh SD medium to an initial OD600 of ∼0.2 to initiate chronological aging. Entry into stationary phase was verified by monitoring pre-stationary growth kinetics. Survival during aging was assessed using outgrowth-based viability measurements in three complementary formats. (i) For high-throughput assays, cells were aged in 96-well plates containing 200 µL SD per well at 30 °C. At defined aging intervals, 2 µL of stationary-phase culture was transferred into fresh YPD in 96-well plates and incubated for 24 h without shaking, after which outgrowth was quantified by OD600 using a microplate reader. (ii) For spot dilution assays, cells were aged in flask cultures at 30 °C with shaking. At selected time points, cultures were normalized to OD600 = 1.0, serially diluted in sterile water, and spotted onto YPD agar plates, which were incubated at 30 °C for 48 h before imaging. (iii) For outgrowth dilution assays, OD600-normalized cultures were serially diluted in YPD in 96-well plates and incubated at 30 °C for 24 h, after which outgrowth was quantified by OD600.

### Yeast growth assays

Yeast growth was assessed using both high-throughput plate-based assays and flask-based spectrophotometric measurements. Prototrophic *Saccharomyces cerevisiae* CEN.PK113-7D wild-type and the indicated mutant strains were cultured in synthetic defined (SD) or synthetic complete (SC) medium at 30 °C, as specified for each experiment. For plate-based growth measurements, cultures were diluted to an initial OD600 of approximately 0.2 and dispensed into 96-well plates. Cells were grown either without treatment or in the presence of experimental treatments applied at single or multiple concentrations, depending on the assay design. Plates were incubated at 30 °C, and growth was quantified by measuring OD600 using a microplate reader. For flask-based growth analyses, yeast cultures were grown in SD or SC medium in glass flasks at 30 °C with shaking at 220 rpm. Cultures were initiated at an OD600 of approximately 0.2 and grown under untreated conditions or with experimental treatments, as appropriate. At defined time points, culture aliquots were transferred to cuvettes and OD600 was measured using a spectrophotometer to quantify growth kinetics.

### RNA extraction and quantitative RT–PCR analysis

Yeast cells were mechanically disrupted according to the manufacturer’s recommended lysis protocol. Total RNA was extracted using the RNeasy Mini Kit (Qiagen). RNA concentration and purity were assessed using an ND-1000 UV–visible spectrophotometer (NanoDrop Technologies). Quantitative reverse transcription–PCR (qRT–PCR) was performed as previously described using the QuantiTect Reverse Transcription Kit (Qiagen) and SYBR FAST Universal qPCR Kit (Kapa Biosystems) ^49,68^. Relative transcript abundance was calculated using *ACT1* as the reference gene. For transcriptomic analyses of methotrexate (MTX), 2,5-anhydromannitol (2,5-AM), 2-deoxy-D-glucose (2-DG), atorvastatin (ATOR), cycloheximide (CHX) treatments, *Saccharomyces cerevisiae* CEN.PK cells were grown exponentially in synthetic defined (SD) medium. Cultures were inoculated at an initial OD600 of 0.2 and grown for 5 h at 30 °C. Cells were then treated with MTX (100 µM) for 1 h, 2,5-AM (16 mM) for 30 min, 2DG (16 mM) for 30 min, ATOR (200 µM) for 30 min, or CHX (5 µM) for 1 h, after which cells were harvested and total RNA was extracted for RNA-sequencing analysis as described below.

### RNA sequencing

RNA concentration and integrity were evaluated using the Agilent 2100 Bioanalyzer with the RNA 6000 Nano LabChip kit. High-quality RNA samples were subjected to paired-end RNA sequencing at Novogene. Polyadenylated mRNA was enriched prior to library construction, followed by cDNA synthesis via reverse transcription. Library quality and fragment size distribution were verified using the Bioanalyzer. Sequencing was performed on the NovaSeq PE150 platform. Raw sequencing reads were processed using the nf-core RNA-seq pipeline (version 3.8.1), employing STAR for read alignment and RSEM for transcript quantification ^79^. Reads were aligned to the *Saccharomyces cerevisiae* reference genome with corresponding Ensembl gene annotations (GTF version 1.105), generating gene-level expression counts ^80^. Normalization and differential expression analyses were performed using DESeq2 ^81^. RNA-sequencing datasets for rapamycin-treated cells were obtained from our previously published studies ^49^.

### Gene ontology and pathway enrichment analysis

Genes exhibiting significant differential expression were subjected to functional annotation and enrichment analyses to identify overrepresented biological processes, molecular functions, and signaling pathways. Gene ontology (GO) and pathway enrichment analyses were performed using the clusterProfiler package in R ^82,83^ and the Metascape online analysis platform ^84^. KEGG pathway visualizations were generated using the pathview package in R ^85^.

### Metabolite analysis

Cell cultures were rapidly quenched by adding two volumes of pre-chilled methanol (−80 °C) and immediately centrifuged at 4,500 rpm for 2 min in a centrifuge maintained at −10 °C. The supernatant was removed, and cell pellets were either stored at −20 °C or processed immediately for metabolite extraction. For extraction, a solvent mixture containing 40% (v/v) acetonitrile, 40% (v/v) methanol, and 20% (v/v) water was prepared using HPLC-grade solvents and pre-cooled to −80 °C. The extraction solvent was mixed thoroughly prior to use. Cell pellets were resuspended in 700 µL of the cold extraction solution by repeated pipetting and incubated on ice for 15 min. Samples were then centrifuged at maximum speed at −4 °C for 5 min to pellet insoluble material. The clarified supernatant was transferred to fresh 1.5 mL microcentrifuge tubes and either dried under vacuum or stored at −80 °C until further analysis. Dried metabolite extracts were reconstituted in 100 µL of 98:2 (v/v) water:methanol and subjected to targeted liquid chromatography–mass spectrometry (LC–MS) analysis as described previously ^69^.

### TORC1 activity assay in yeast

TORC1 signaling activity was assessed by monitoring phosphorylation of the downstream TORC1 effector Sch9 using an immunoblotting-based assay with minor modifications ^68,69^. Prototrophic Saccharomyces cerevisiae CEN.PK113-7D cells expressing HA-tagged Sch9 (Sch9–6×HA) were cultured under the different growth conditions and subjected to chemical treatments. Cells were harvested at defined time points and processed for total protein extraction. Protein extracts were separated by SDS–PAGE and transferred onto nitrocellulose membranes. Membranes were blocked in 5% (w/v) non-fat milk prepared in TBS containing 0.1% Tween-20 prior to antibody incubation. Phosphorylation-dependent mobility shifts of Sch9 were detected using an anti-HA (3F10) antibody (1:2000), followed by incubation with a horseradish peroxidase–conjugated goat anti-rat secondary antibody (1:5000). Immunoreactive signals were visualized using enhanced chemiluminescence (ECL Prime), and images were acquired using the imaging system.

### Transcriptional profiling using Connectivity Map (CMap) / LINCS L1000 dataset

1. Data acquisition and preprocessing: Perturbation signatures were obtained from the Connectivity Map (CMap) / LINCS L1000 dataset ^63,64,86–89^. Level 5 consensus signatures (file: level5_beta_trt_cp_n720216×12328.gctx) were retrieved via the CLUE platform, in which replicate measurements were integrated using the moderated Z-score (MODZ) algorithm to improve signal fidelity and reduce experimental noise. The resulting gene-by-signature matrix represents differential expression relative to controls, where both magnitude and sign indicate the strength and direction of transcriptional regulation.

2. Quantification of perturbation intensity: To capture the global transcriptional impact of each perturbation, we defined a perturbation score (Sp) based on the most responsive genes: Gene ranking: Genes within each signature were ranked according to expression change. Selection of responsive genes: The top 20, 100, and 200 upregulated and downregulated genes were selected as primary responsive subsets.

Score calculation: The perturbation score was calculated as the difference between the mean expression of the top and bottom gene sets:

Sp = mean(E_top100) − mean(E_bottom100)

This approach emphasizes highly responsive genes, providing a robust signal-to-noise ratio while minimizing the influence of non-responsive genes and outliers.

3. Comparative analysis and visualization: Initial distribution analyses across multiple doses and time points identified 10 μM at 24 hours as the optimal condition for inducing consistent and high-intensity transcriptional responses. Subsequent analyses focused on five benchmark compounds: sirolimus, temsirolimus, methotrexate, atorvastatin, and cycloheximide.

For each compound within a given cell line, gene ranks were aggregated across replicates using the median rank. Consistently differentially expressed genes (DEGs) were defined from the top and bottom of this aggregated distribution. Heatmaps were generated in R (v4.4.2) using the pheatmap package. Genes not meeting the defined regulatory threshold in a given cell line were assigned a sentinel value and displayed in white to preserve visual clarity.

4. Functional annotation: Functional enrichment analysis of DEGs was performed using Metascape to identify biological pathways associated with each perturbation signature ^84^.

5. Cross-cell-line consistency of sirolimus response: To assess conservation of transcriptional responses, sirolimus (BRD-K84937637) signatures were extracted across cell lines under matched experimental conditions, enabling direct comparison of its transcriptional footprint.

### Cell viability measurement

Cell viability was determined using the Cell Counting Kit-8 (CCK-8; Dojindo, Japan) following the manufacturer’s protocol. Cells were seeded in 96-well plates and cultured under nutrient-limited (1% FBS) or nutrient-rich (10% FBS) conditions with the indicated treatments for 48 h. After treatment, 10 µL of CCK-8 reagent was added to each well and incubated at 37 °C. Absorbance was then measured at 450 nm using a microplate reader.

### Western blot

Cells were lysed in RIPA buffer (Thermo Fisher Scientific, USA) supplemented with 1× Halt™ protease and phosphatase inhibitor cocktail (Thermo Fisher Scientific, USA). Protein concentrations were determined using the Pierce™ BCA Protein Assay Kit (Thermo Fisher Scientific, USA). Equal amounts of protein were mixed with 6× Laemmli sample buffer (Nacalai Tesque, Japan) containing 5% β-mercaptoethanol (Sigma-Aldrich, USA) and denatured prior to loading. Samples were separated by SDS–PAGE at 100 V for 2 hours and subsequently transferred onto nitrocellulose membranes (Bio-Rad, USA) at 85 V for 1.5 hours. Membranes were then blocked for 1 hour in 1% (w/v) fish skin gelatin (FSG; Sigma-Aldrich, USA) prepared in Tris-buffered saline with Tween-20 (TBST). Membranes were incubated overnight at 4°C with primary antibodies against total mTOR (1:1000, CST-2983S) and p-mTOR (Ser-2448) (1:1000, CST-2971), total p70 S6 Kinase (1:1000, CST-9202S) and p-p70 S6 Kinase (Thr389) (1:1000, CST-9234S), total S6 Ribosomal Protein RPS6 (1:1000, CST-2217S), P-S6 Ribosomal Protein RPS6 (Ser-240/244), (1:1000, CST-5364S), total ULK1 (1:1000, CST-8054S) and p-ULK1 (Ser555) (1:1000, CST-5869S), total AMPKα (1:1000, CST-5831S) and p-AMPKα (Thr172) (1:1000,CST-2535S), total ACC (1:1000, CST-3662S) and p-ACC (Ser79) (1:1000, CST-3661S), SIRT1 (1:1000, 13161-1-AP), PGC1α (1:1000, NBP1-04676), TFAM (1:1000, CST-8076S), SOD2 (1:1000, ab13533), NRF2 (1:1000, CST-33649S), NDUFS3 (1:1000, ab14711), beta-Actin (1:1000, CST; 8457S) followed by incubation with HRP-conjugated secondary antibodies (HRP-linked anti-rabbit IgG (1:3000, CST; 7074S) for 1 hour at room temperature. Detection was performed using Clarity™ Western ECL Blotting Substrate (Bio-Rad, USA) and imaged using a Bio-Rad ChemiDoc Imaging System. Phospho-protein membranes were stripped using Restore™ PLUS Stripping Buffer (Thermo Fisher Scientific, USA).

### Statistical analysis

The analysis of all experimental data, including calculations of mean values, standard deviations, significance, and graphical representation, was conducted using GraphPad Prism v.11 software. In all graphical representations, significance levels are denoted as *P < 0.05, **P < 0.01, ***P < 0.001, and ****P < 0.0001. Results with P values falling below these thresholds were considered statistically significant, while ‘ns’ signifies non-significance.

## Data availability

Further information and requests for resources and reagents should be directed to and will be fulfilled by the Lead Contact, Dr. Mohammad Alfatah.

## ACKNOWLEDGMENTS

This work is supported by the Young Investigator Research Grant (YIRG), National Medical Research Council, Singapore (MOH-001348-00) and US NAM Healthy Longevity Catalyst Awards Grant (MOH-001439). Arshia Naaz is supported by A*STAR-CDF grant (C243512027).

## AUTHOR CONTRIBUTIONS STATEMENT

Trishia Cheng Yi Ning: Investigation, Formal analysis

Arshia Naaz: Investigation, Formal analysis, Funding Acquisition

Mingtong Gao: Investigation, Formal analysis

Wu Hao: Investigation, Formal analysis

Yizhong Zhnag: Investigation, Formal analysis

Liang Cui: Investigation, Formal analysis

Jovian Jing Lin: Investigation, Formal analysis

Sonia Yogasundaram: Investigation, Formal analysis

Nashrul Afiq Faidzinn: Investigation, Formal analysis

Ong Yong Qing Victoria: Investigation, Formal analysis

Rajkumar Dorajoo: Review and editing

Brian K Kennedy: Review and editing

Mohammad Alfatah: Conceptualization, Supervision, Writing-original draft, Writing-review and editing, Funding acquisition.

All authors read, critically reviewed and approved the final manuscript.

M.A is the guarantor of this work.

