## Supplemental Figures for "TORC1 integrates metabolic state transitions during aging"

Affiliations:
1. Healthy Longevity Translational Research Program, Yong Loo Lin School of Medicine, National University of Singapore, Singapore, Singapore.
2. Centre for Healthy Longevity, National University Health System, Singapore, Singapore.
3. Genome Institute of Singapore (GIS), Agency for Science, Technology and Research (A*STAR), 60 Biopolis Street, Genome #02-01, Singapore 138672, Republic of Singapore.
4. Bioinformatics Institute (BII), Agency for Science, Technology and Research (A*STAR), 30 Biopolis Street, Matrix #07-01, Singapore 138671, Republic of Singapore.
5. Antimicrobial Resistance Interdisciplinary Research Group, Singapore-MIT Alliance for Research and Technology, 1 CREATE Way, Singapore 138602, Singapore.
6. Department of Paediatrics, Yong Loo Lin School of Medicine, National University of Singapore, Singapore 119228, Singapore.
7. Departments of Biochemistry and Physiology, Yong Loo Lin School of Medicine, National University of Singapore, Singapore, Singapore.

#Equal authorship contributions

*To whom the correspondence should be addressed.
 (Mohammad Alfatah)

**Keywords**
TORC1 signaling; metabolic state transitions; cellular aging; nutrient limitation; systems metabolism; anabolic metabolism; metabolic adaptation; mitochondrial function; SEACIT/GATOR1; cellular survival

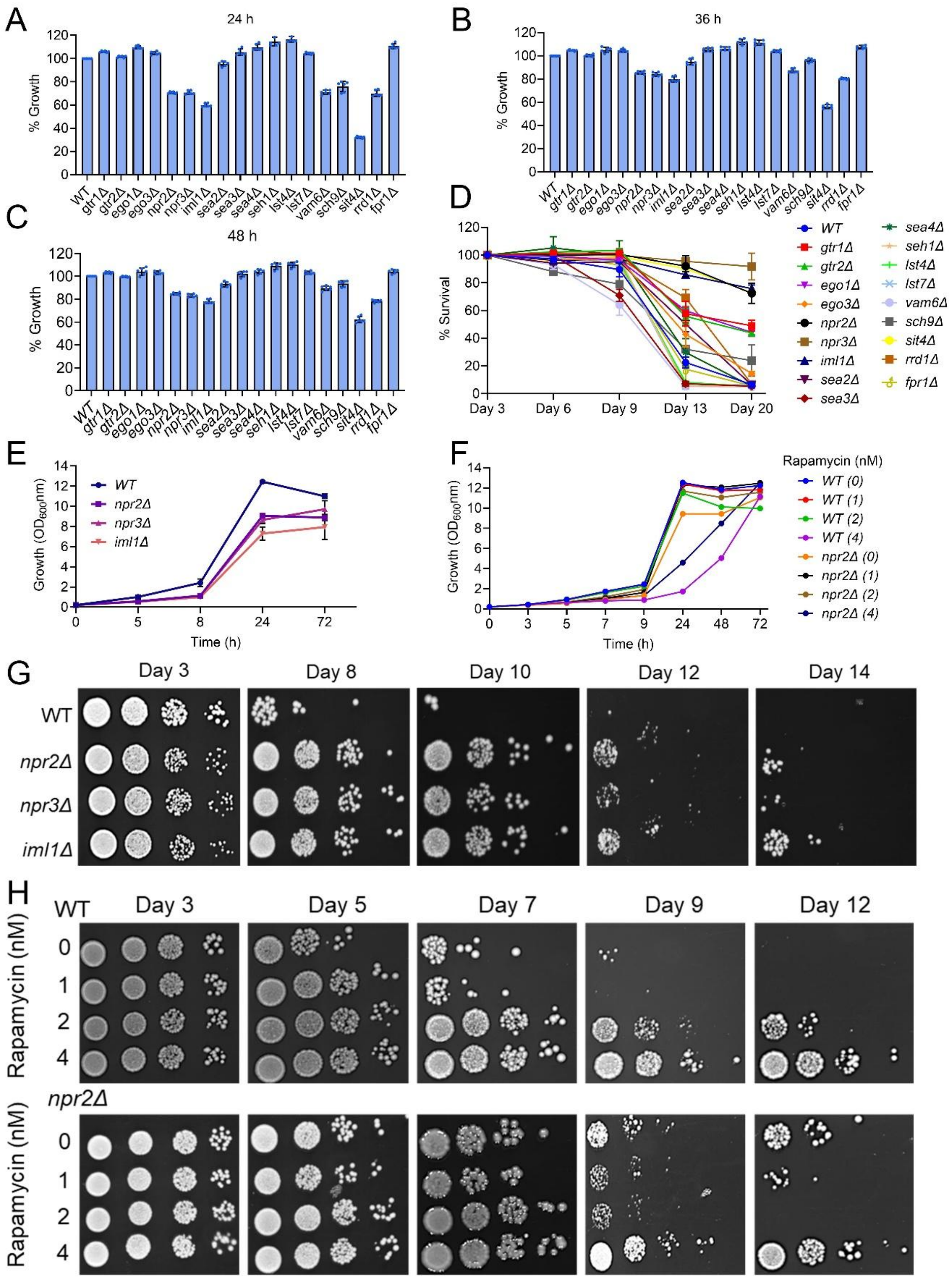


**Figure S1. Temporal growth and survival dynamics of TORC1 pathway mutants, Related to Figure 1**

(A–C) Relative growth of wild-type and indicated TORC1 pathway mutants measured after 24 h (A), 36 h (B), and 48 h (C) of culture in synthetic defined (SD) medium. Values are normalized to wild-type and provide extended temporal resolution corresponding to the 12 h growth measurements shown in Figure 1B (mean ± SD, n = 6).

(D) Full stationary-phase survival time course for wild-type and indicated mutants, showing the percentage of viable cells from day 3 to day 20. For clarity, survival at day 13 and day 20 is highlighted in the main text (Figure 1C–D).

(E) Growth kinetics of wild-type and SEACIT-deficient strains (*npr2Δ*, *npr3Δ*, *iml1Δ*) measured over time in glass flask cultures.

(F) Growth kinetics of wild-type and *npr2Δ* cells in the presence of increasing concentrations of rapamycin, measured over time in glass flask cultures.

(G) Stationary-phase survival of wild-type and SEACIT-deficient strains assessed by spot dilution assays at the indicated days. Data are replicates of Figure 1G.

(H) Stationary-phase survival of wild-type and *npr2Δ* cells treated with rapamycin, assessed by spot dilution assays at the indicated days. Data are replicates of Figure 1H.

A

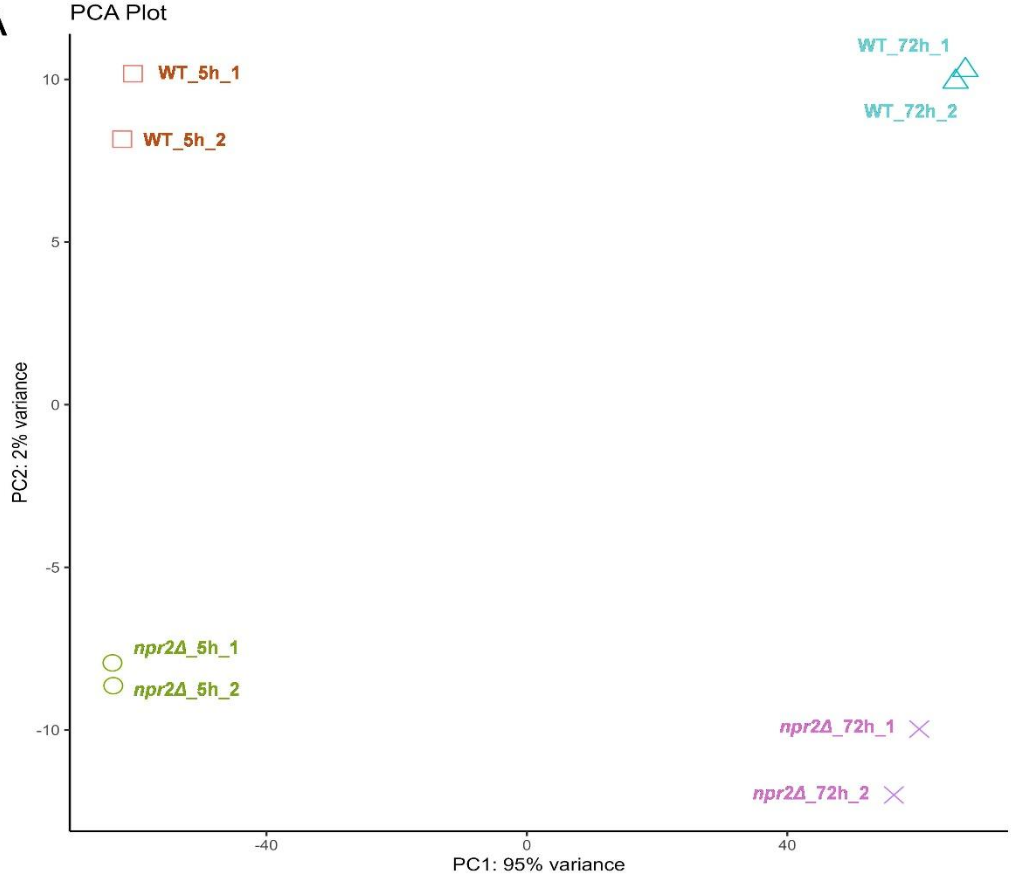
PCA Plot
WT_5h_1
WT_5h_2
WT_72h_1
WT_72h_2
npr2Δ_5h_1
npr2Δ_5h_2
npr2Δ_72h_1
npr2Δ_72h_2
PC2: 2% variance
PC1: 95% variance
10
5
0
-5
-10
-40
0
40

B

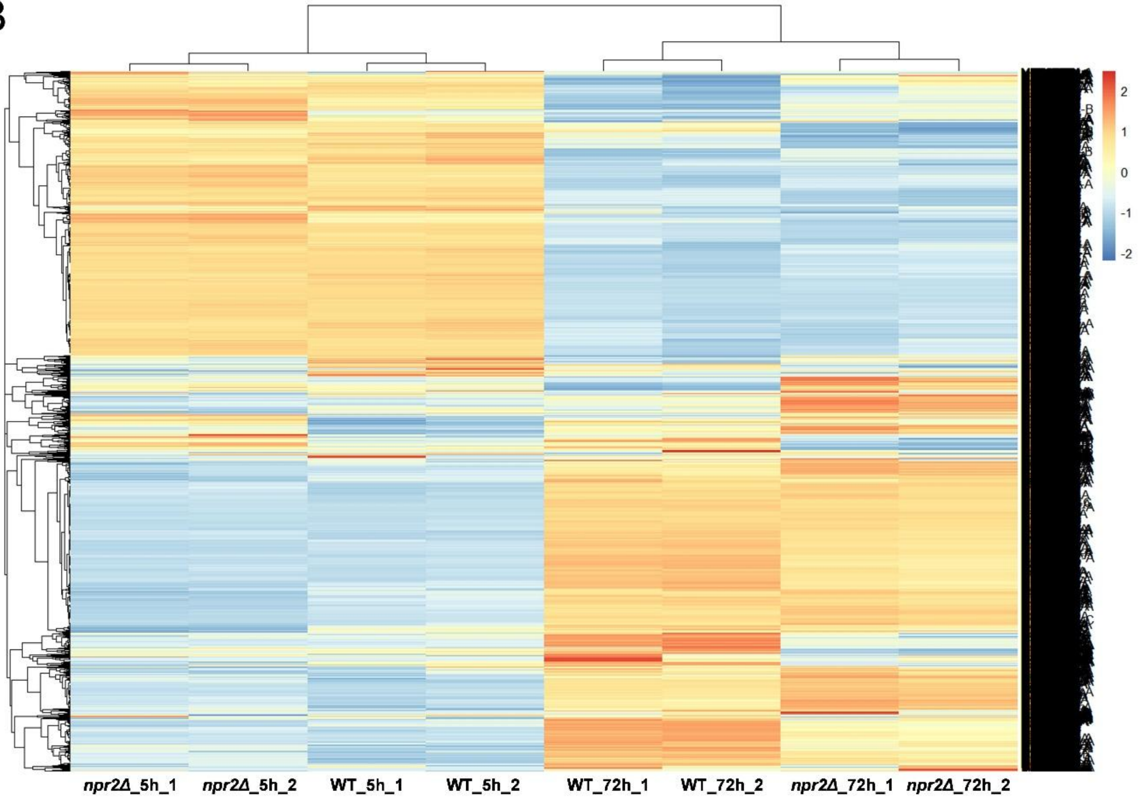
2
1
0
-1
-2
npr2Δ_5h_1
npr2Δ_5h_2
WT_5h_1
WT_5h_2
WT_72h_1
WT_72h_2
npr2Δ_72h_1
npr2Δ_72h_2

**Figure S2. Principal component and clustering analysis of transcriptomic states, Related to Figure 2**

(A) Principal component analysis (PCA) of transcriptomic profiles from wild-type and *npr2Δ* cells harvested during exponential (5 h) and stationary (72 h) phases. PC1 and PC2 explain the indicated proportions of variance and show separation by growth phase and genotype.

(B) Unsupervised hierarchical clustering heatmap of normalized gene expression across all samples, supporting the phase- and *NPR2*-dependent transcriptional differences shown in Figure 2.

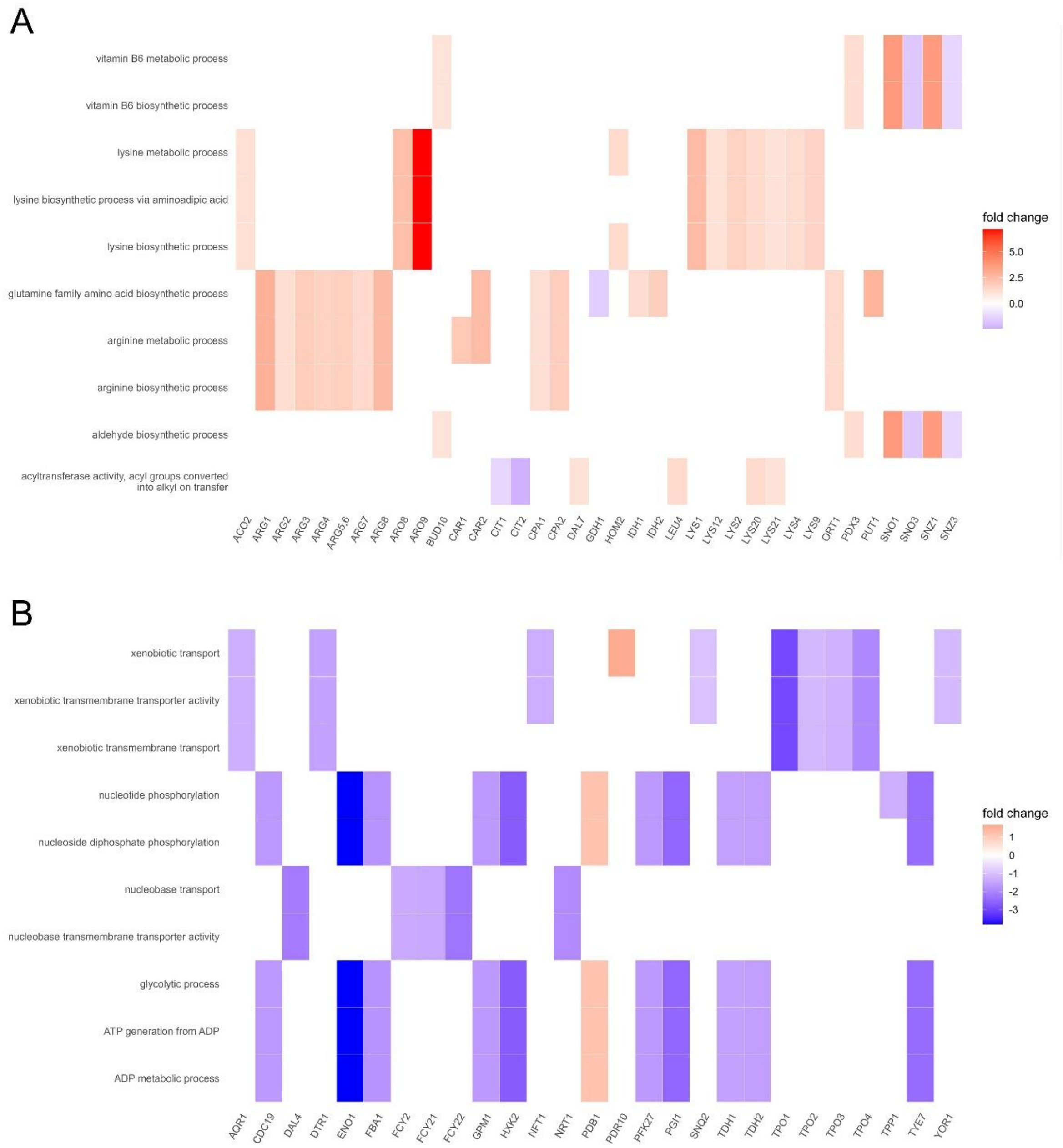


**Figure S3. Phase-specific transcriptional programs associated with NPR2 loss, Related to Figure 2**

(A) Heatmap showing differentially expressed genes (DEGs) commonly modulated in *npr2Δ* cells during the exponential (5 h) phase. Colors represent fold change relative to wild-type.

(B) Heatmap showing DEGs commonly modulated in *npr2Δ* cells during the stationary (72 h) phase, highlighting phase-specific transcriptional programs associated with *NPR2* loss. Colors represent fold change relative to wild-type.

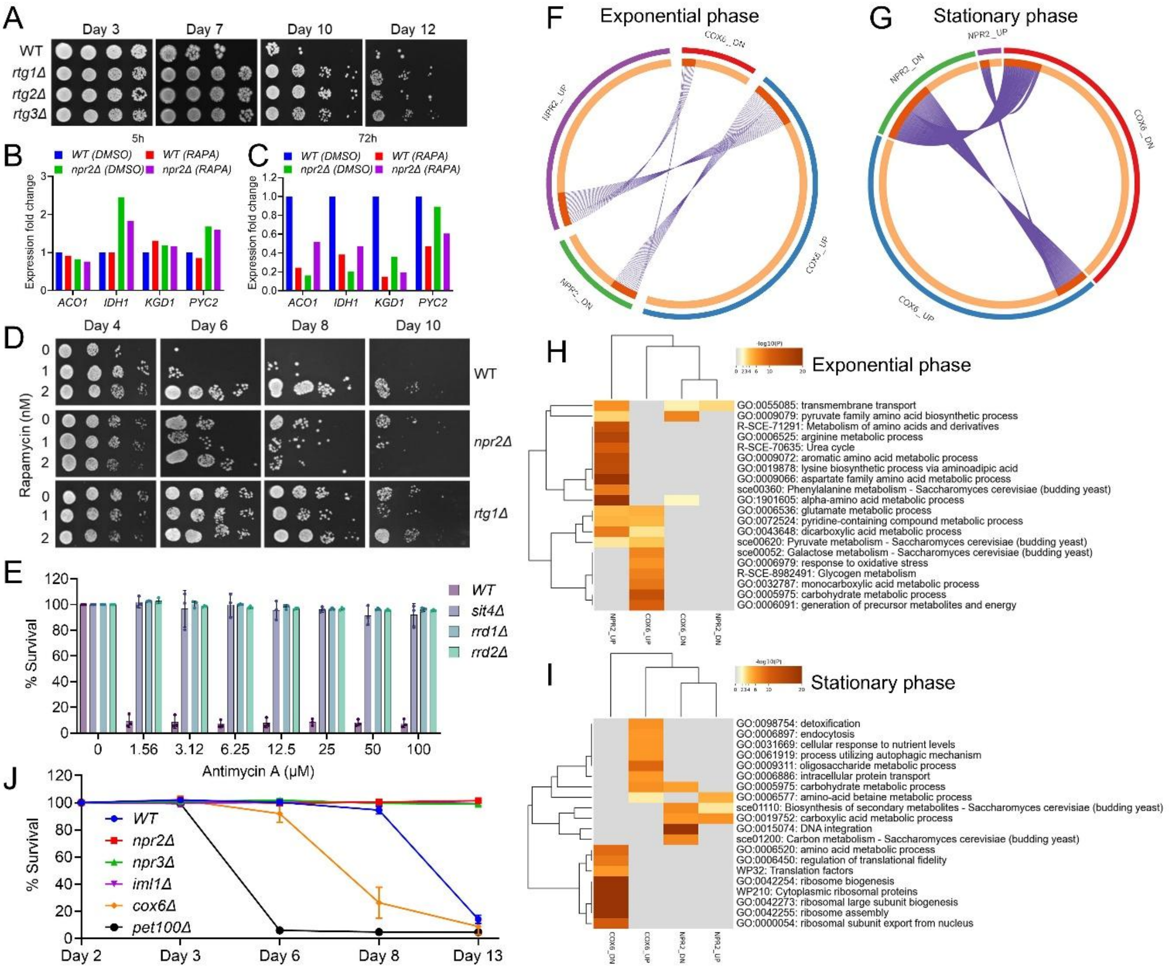


**Figure S4. Mitochondrial and retrograde signaling regulate survival downstream of TORC1, Related to Figure 3**

(A) Survival assessed by spot dilution assays for wild-type and retrograde signaling mutants (*rtg1Δ*, *rtg2Δ*, *rtg3Δ*) at the indicated days during stationary phase.

(B–C) Expression of mitochondrial metabolic genes (*ACO1*, *IDH1*, *KGD1*, *PYC2*) in wild-type and *npr2Δ* cells treated with DMSO or rapamycin, measured during exponential growth (5 h; B) and stationary phase (72 h; C).

(D) Survival of wild-type, *npr2Δ*, and *rtg1Δ* cells treated with increasing concentrations of rapamycin, monitored by spot dilution assays at the indicated days. Data are replicates of Figure 3C.

(E) Quantification of survival following treatment with increasing concentrations of antimycin A for wild-type and phosphatase pathway mutants (*sit4Δ*, *rrd1Δ*, *rrd2Δ*). Survival is expressed as a percentage relative to untreated controls. Data represent mean ± SD (n = 3).

(F–G) Circular plots illustrating gene-level overlap between transcriptional responses associated with *NPR2* and *COX6* perturbation during exponential growth (F) and stationary phase (G).

(H–I) Heatmaps of Gene Ontology (GO) enrichment for biological processes associated with transcriptional changes linked to *NPR2* and *COX6* during exponential growth (H) and stationary phase (I). Color scale represents –log10(P value).

(J) Time-course analysis of survival for wild-type and *npr2Δ*, *npr3Δ*, *iml1Δ*, *cox6Δ*, *pet100Δ* mutants.

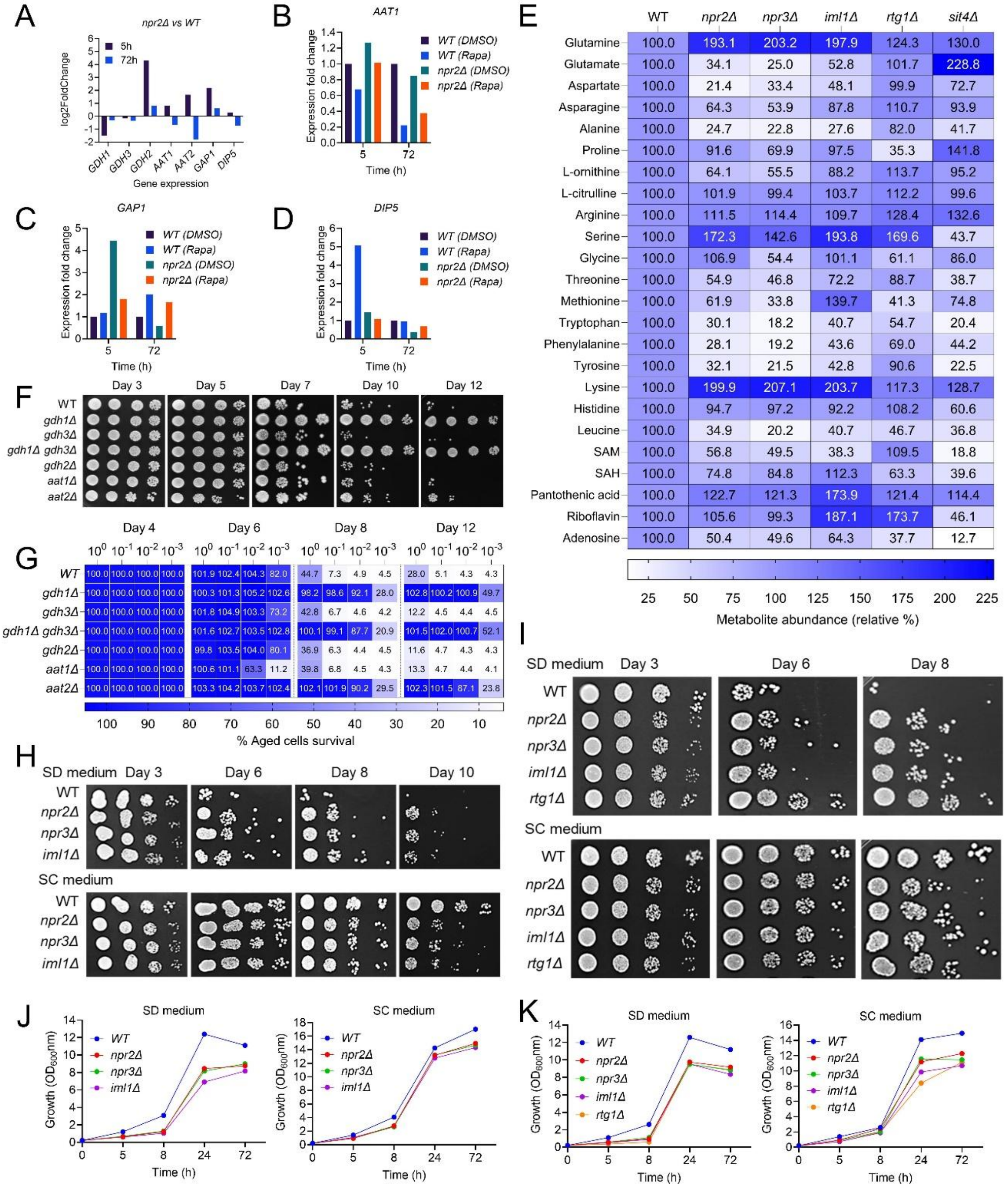


**Figure S5. Nitrogen metabolism remodeling and survival downstream of SEACIT–TORC1 signaling, Related to Figure 3**

(A) RNA-sequencing–derived log2 fold change of selected nitrogen metabolism–associated genes in *npr2Δ* relative to wild-type during exponential growth (5 h) and stationary phase (72 h). RNA-seq data correspond to Supplementary Table 1.

(B–D) Expression fold change of *AAT1* (B), *GAP1* (C), and *DIP5* (D) in wild-type and *npr2Δ* cells treated with DMSO or rapamycin, measured during exponential growth (5 h) and stationary phase (72 h).

(E) Heatmap showing relative intracellular metabolite abundance in *npr2Δ*, *npr3Δ*, and *iml1Δ* cells compared with wild-type (set to 100%), with additional inclusion of *rtg1Δ* and *sit4Δ*. Metabolites associated with amino acid, nitrogen, and one-carbon metabolism are shown.

(F) Survival assessed by spot dilution assays for wild-type and indicated amino acid metabolism mutants (*gdh1Δ*, *gdh3Δ*, *gdh1Δ gdh3Δ*, *gdh2Δ*, *aat1Δ*, and *aat2Δ*). Data are replicates of Figure 3J.

(G) Quantification of survival for the strains shown in (F) using an outgrowth dilution assay in YPD medium, expressed as a percentage relative to wild-type.

(H–I) Spot dilution–based survival analysis of wild-type, SEACIT mutants (*npr2Δ*, *npr3Δ*, *iml1Δ*), and *rtg1Δ* cells aged in synthetic defined (SD) or synthetic complete (SC) medium.

(J–K) Growth kinetics of wild-type and indicated mutants measured over time in SD or SC medium.

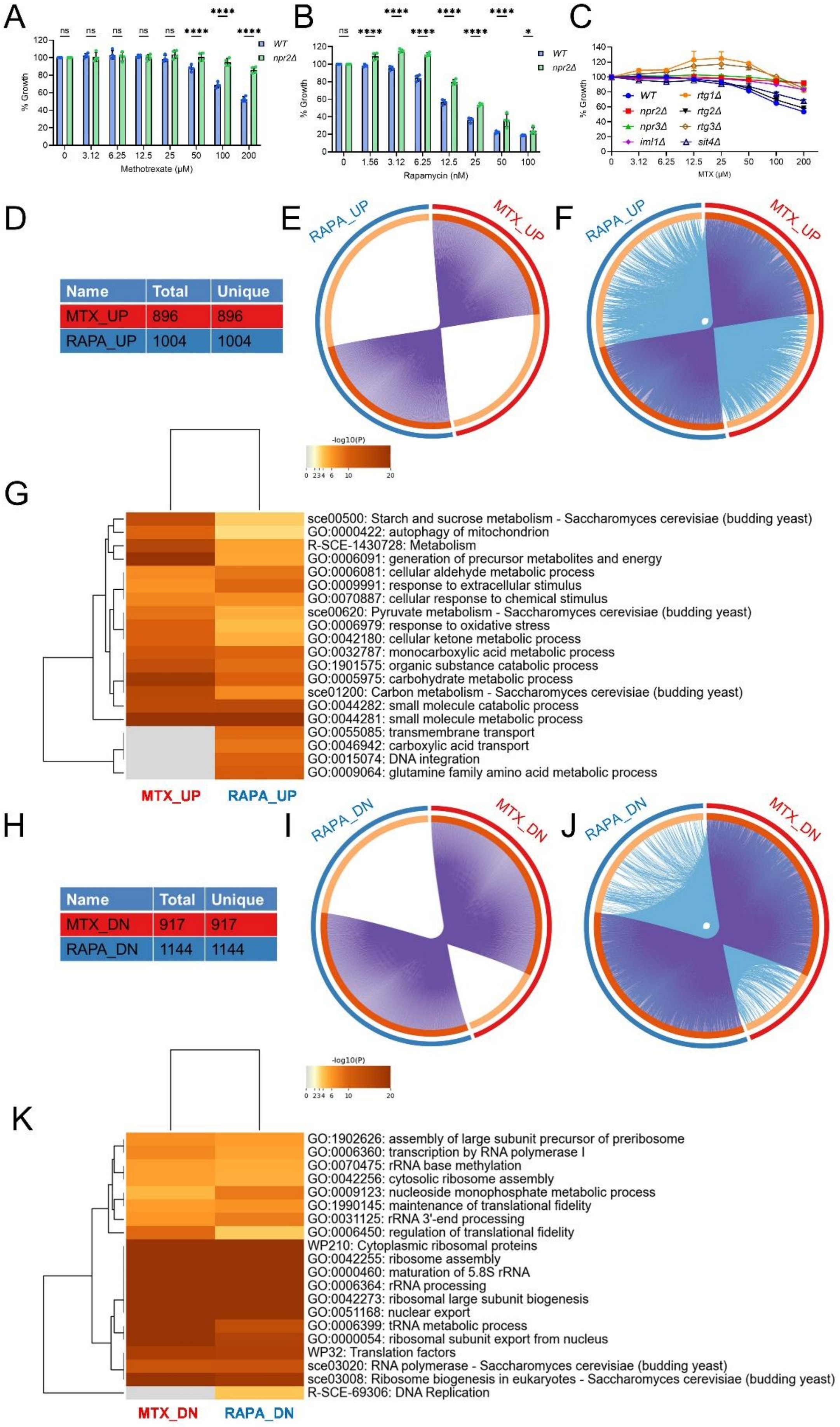
A
B
C
WT
npr2Δ
% Growth
Methotrexate (μM)
Rapamycin (nM)
MTX (μM)
rtg1Δ
rtg2Δ
npr3Δ
rtg3Δ
iml1Δ
sit4Δ
D
E
F
RAPA_UP
MTX_UP
Name | Total | Unique
MTX_UP | 896 | 896
RAPA_UP | 1004 | 1004
-log10(P)
G
sce00500: Starch and sucrose metabolism - Saccharomyces cerevisiae (budding yeast)
GO:0000422: autophagy of mitochondrion
R-SCE-1430728: Metabolism
GO:0006091: generation of precursor metabolites and energy
GO:0006081: cellular aldehyde metabolic process
GO:0009991: response to extracellular stimulus
GO:0070887: cellular response to chemical stimulus
sce00620: Pyruvate metabolism - Saccharomyces cerevisiae (budding yeast)
GO:0006979: response to oxidative stress
GO:0042180: cellular ketone metabolic process
GO:0032787: monocarboxylic acid metabolic process
GO:1901575: organic substance catabolic process
GO:0005975: carbohydrate metabolic process
sce01200: Carbon metabolism - Saccharomyces cerevisiae (budding yeast)
GO:0044282: small molecule catabolic process
GO:0044281: small molecule metabolic process
GO:0055085: transmembrane transport
GO:0046942: carboxylic acid transport
GO:0015074: DNA integration
GO:0009064: glutamine family amino acid metabolic process
MTX_UP
RAPA_UP
H
I
J
RAPA_DN
MTX_DN
Name | Total | Unique
MTX_DN | 917 | 917
RAPA_DN | 1144 | 1144
-log10(P)
K
GO:1902626: assembly of large subunit precursor of preribosome
GO:0006360: transcription by RNA polymerase I
GO:0070475: rRNA base methylation
GO:0042256: cytosolic ribosome assembly
GO:0009123: nucleoside monophosphate metabolic process
GO:1990145: maintenance of translational fidelity
GO:0031125: rRNA 3'-end processing
GO:0006450: regulation of translational fidelity
WP210: Cytoplasmic ribosomal proteins
GO:0042255: ribosome assembly
GO:0000460: maturation of 5.8S rRNA
GO:0006364: rRNA processing
GO:0042273: ribosomal large subunit biogenesis
GO:0051168: nuclear export
GO:0006399: tRNA metabolic process
GO:0000054: ribosomal subunit export from nucleus
WP32: Translation factors
sce03020: RNA polymerase - Saccharomyces cerevisiae (budding yeast)
sce03008: Ribosome biogenesis in eukaryotes - Saccharomyces cerevisiae (budding yeast)
R-SCE-69306: DNA Replication
MTX_DN
RAPA_DN

**Figure S6. Methotrexate recapitulates TORC1 inhibition–associated growth and transcriptional programs, Related to Figure 4**

(A–B) Relative growth of wild-type and *npr2Δ* cells treated with increasing concentrations of methotrexate (MTX) (A) or rapamycin (B) for 24 h. Growth is expressed as percentage relative to untreated controls. Data are shown as mean ± SD (n = 4). Statistical significance was assessed by two-way ANOVA ($P < 0.05$, ****$P < 0.0001$; ns, not significant).

(C) Relative growth of additional TORC1-pathway mutants following methotrexate (MTX) treatment, extending the analysis shown in Figure 4C. Strains include *rtg1Δ*, *rtg2Δ*, *rtg3Δ*, and *sit4Δ*. Data are shown as mean ± SD (n = 2).

(D) Summary table showing the total number of genes upregulated following MTX or rapamycin treatment.

(E) Circular plot showing gene-level overlap between genes upregulated by methotrexate (MTX_UP) and rapamycin (RAPA_UP).

(F) Circular plot showing functional overlap of Gene Ontology (GO) enrichments associated with MTX_UP and RAPA_UP gene sets.

(G) Heatmap of GO enrichment for biological processes associated with genes upregulated by MTX or rapamycin.

(H) Summary table showing the total number of genes downregulated following MTX or rapamycin treatment.

(I) Circular plot showing gene-level overlap between genes downregulated by methotrexate (MTX_DN) and rapamycin (RAPA_DN).

(J) Circular plot showing functional overlap of GO enrichments associated with MTX_DN and RAPA_DN gene sets.

(K) Heatmap of GO enrichment for biological processes associated with genes downregulated by MTX or rapamycin.

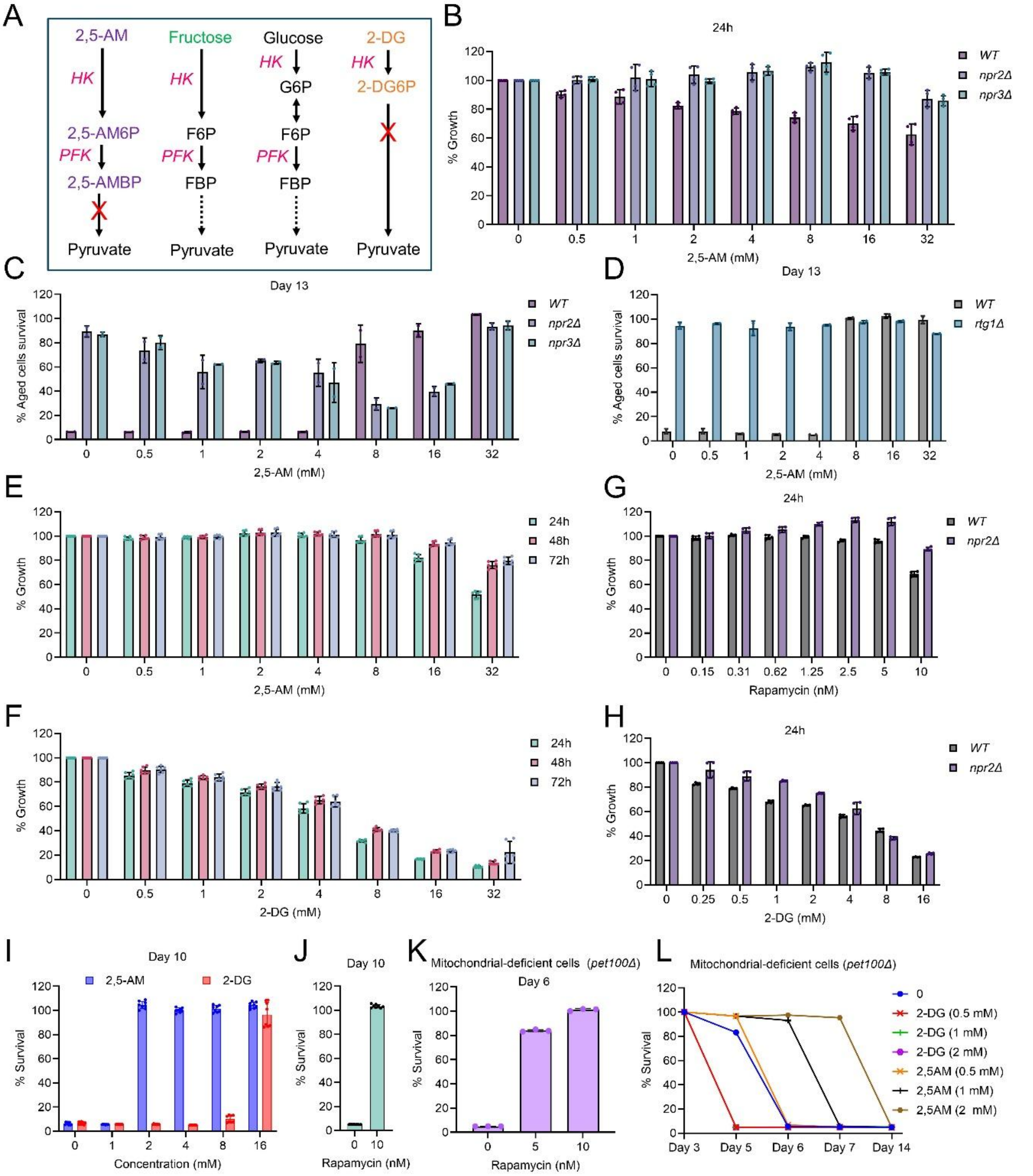


**Figure S7. Glycolytic perturbations differentially regulate growth and survival in a mitochondrial-dependent manner, Related to Figure 5**

(A) Schematic representation of glycolytic entry points for 2,5-anhydromannitol (2,5-AM), fructose, glucose, and 2-deoxyglucose (2-DG), illustrating differential enzymatic processing and points of metabolic blockade.

(B) Relative growth of wild-type, *npr2Δ*, and *npr3Δ* cells treated with increasing concentrations of 2,5-AM. Growth is expressed as percentage relative to untreated controls. Data are shown as mean ± SD (n = 4). Statistical significance was assessed by two-way ANOVA ($P < 0.05$, ****$P < 0.0001$; ns, not significant).

(C) Cell survival during stationary phase on day 13 for wild-type, *npr2Δ*, and *npr3Δ* cells treated with increasing concentrations of 2,5-AM, assessed using a chronological aging assay, mean ± SD (n = 2).

(D) Cell survival during stationary phase on day 13 for wild-type and *rtg1Δ* cells treated with increasing concentrations of 2,5-AM, mean ± SD (n = 2).

(E) Relative growth of wild-type cells treated with increasing concentrations of 2,5-AM measured at 24 h, 48 h, and 72 h, mean ± SD (n = 6).

(F) Relative growth of wild-type cells treated with increasing concentrations of 2-DG measured at 24 h, 48 h, and 72 h, mean ± SD (n = 6).

(G) Relative growth of wild-type and *npr2Δ* cells treated with increasing concentrations of rapamycin measured at 24 h, mean ± SD (n = 4).

(H) Relative growth of wild-type and *npr2Δ* cells treated with increasing concentrations of 2-DG measured at 24 h, mean ± SD (n = 4).

(I) Cell survival during stationary phase on day 10 for wild-type cells treated with 2,5-AM or 2-DG, mean ± SD (n = 8).

(J) Cell survival during stationary phase on day 10 for wild-type cells treated with rapamycin, mean ± SD (n = 8).

(K) Cell survival during stationary phase on day 6 for mitochondrial-deficient (*pet100Δ*) cells treated with rapamycin, mean ± SD (n = 3).

(L) Time-course analysis of stationary-phase survival for mitochondrial-deficient (*pet100Δ*) cells treated with 2-DG or 2,5-AM at the indicated concentrations, mean ± SD (n = 3).

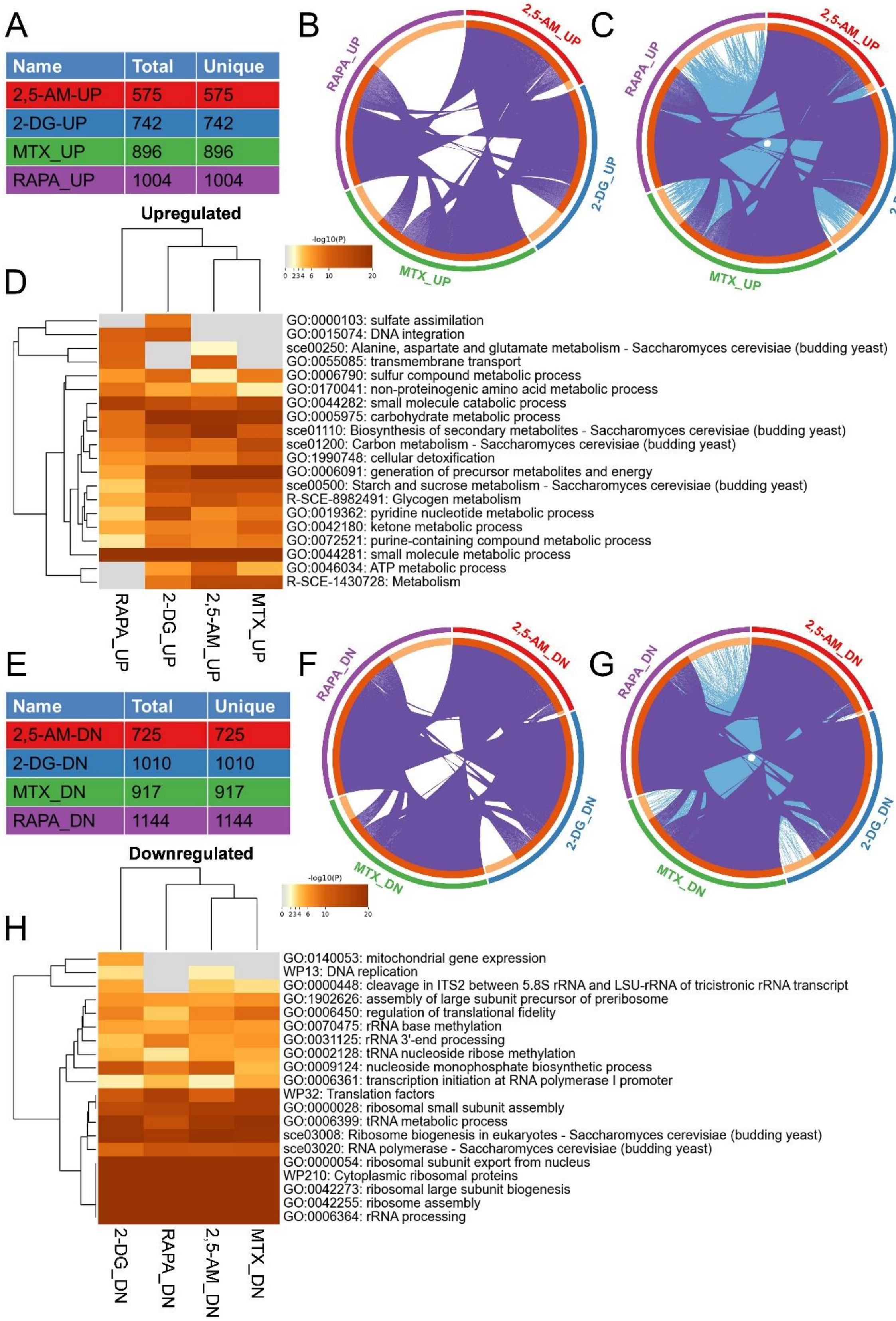
A
Name | Total | Unique
2,5-AM-UP | 575 | 575
2-DG-UP | 742 | 742
MTX_UP | 896 | 896
RAPA_UP | 1004 | 1004
B
C
2,5-AM_UP
2-DG_UP
MTX_UP
RAPA_UP
Upregulated
-log10(P)
0 234 6 10 20
D
GO:0000103: sulfate assimilation
GO:0015074: DNA integration
sce00250: Alanine, aspartate and glutamate metabolism - Saccharomyces cerevisiae (budding yeast)
GO:0055085: transmembrane transport
GO:0006790: sulfur compound metabolic process
GO:0170041: non-proteinogenic amino acid metabolic process
GO:0044282: small molecule catabolic process
GO:0005975: carbohydrate metabolic process
sce01110: Biosynthesis of secondary metabolites - Saccharomyces cerevisiae (budding yeast)
sce01200: Carbon metabolism - Saccharomyces cerevisiae (budding yeast)
GO:1990748: cellular detoxification
GO:0006091: generation of precursor metabolites and energy
sce00500: Starch and sucrose metabolism - Saccharomyces cerevisiae (budding yeast)
R-SCE-8982491: Glycogen metabolism
GO:0019362: pyridine nucleotide metabolic process
GO:0042180: ketone metabolic process
GO:0072521: purine-containing compound metabolic process
GO:0044281: small molecule metabolic process
GO:0046034: ATP metabolic process
R-SCE-1430728: Metabolism
RAPA_UP
2-DG_UP
2,5-AM_UP
MTX_UP
E
Name | Total | Unique
2,5-AM-DN | 725 | 725
2-DG-DN | 1010 | 1010
MTX_DN | 917 | 917
RAPA_DN | 1144 | 1144
F
G
2,5-AM_DN
2-DG_DN
MTX_DN
RAPA_DN
Downregulated
-log10(P)
0 234 6 10 20
H
GO:0140053: mitochondrial gene expression
WP13: DNA replication
GO:0000448: cleavage in ITS2 between 5.8S rRNA and LSU-rRNA of tricistronic rRNA transcript
GO:1902626: assembly of large subunit precursor of preribosome
GO:0006450: regulation of translational fidelity
GO:0070475: rRNA base methylation
GO:0031125: rRNA 3'-end processing
GO:0002128: tRNA nucleoside ribose methylation
GO:0009124: nucleoside monophosphate biosynthetic process
GO:0006361: transcription initiation at RNA polymerase I promoter
WP32: Translation factors
GO:0000028: ribosomal small subunit assembly
GO:0006399: tRNA metabolic process
sce03008: Ribosome biogenesis in eukaryotes - Saccharomyces cerevisiae (budding yeast)
sce03020: RNA polymerase - Saccharomyces cerevisiae (budding yeast)
GO:0000054: ribosomal subunit export from nucleus
WP210: Cytoplasmic ribosomal proteins
GO:0042273: ribosomal large subunit biogenesis
GO:0042255: ribosome assembly
GO:0006364: rRNA processing
2-DG_DN
RAPA_DN
2,5-AM_DN
MTX_DN

**Figure S8. Transcriptional and functional convergence of 2,5-anhydromannitol and 2-deoxyglucose with TORC1-linked metabolic restraint, Related to Figure 5**

(A) Summary table showing the total number of genes upregulated following treatment with 2,5-anhydromannitol (2,5-AM), 2-deoxyglucose (2-DG), methotrexate (MTX), or rapamycin (RAPA).

(B) Circular plot showing gene-level overlap among genes upregulated by 2,5-AM, 2-DG, MTX, and rapamycin.

(C) Circular plot showing functional overlap of Gene Ontology (GO) enrichments associated with genes upregulated by 2,5-AM, 2-DG, MTX, and rapamycin.

(D) Heatmap of GO enrichment for biological processes associated with genes upregulated by 2,5-AM, 2-DG, MTX, and rapamycin. Color scale represents −log10(P value).

(E) Summary table showing the total number of genes downregulated following treatment with 2,5-AM, 2-DG, MTX, or rapamycin.

(F) Circular plot showing gene-level overlap among genes downregulated by 2,5-AM, 2-DG, MTX, and rapamycin.

(G) Circular plot showing functional overlap of GO enrichments associated with genes downregulated by 2,5-AM, 2-DG, MTX, and rapamycin.

(H) Heatmap of GO enrichment for biological processes associated with genes downregulated by 2,5-AM, 2-DG, MTX, and rapamycin. Color scale represents −log10(P value).

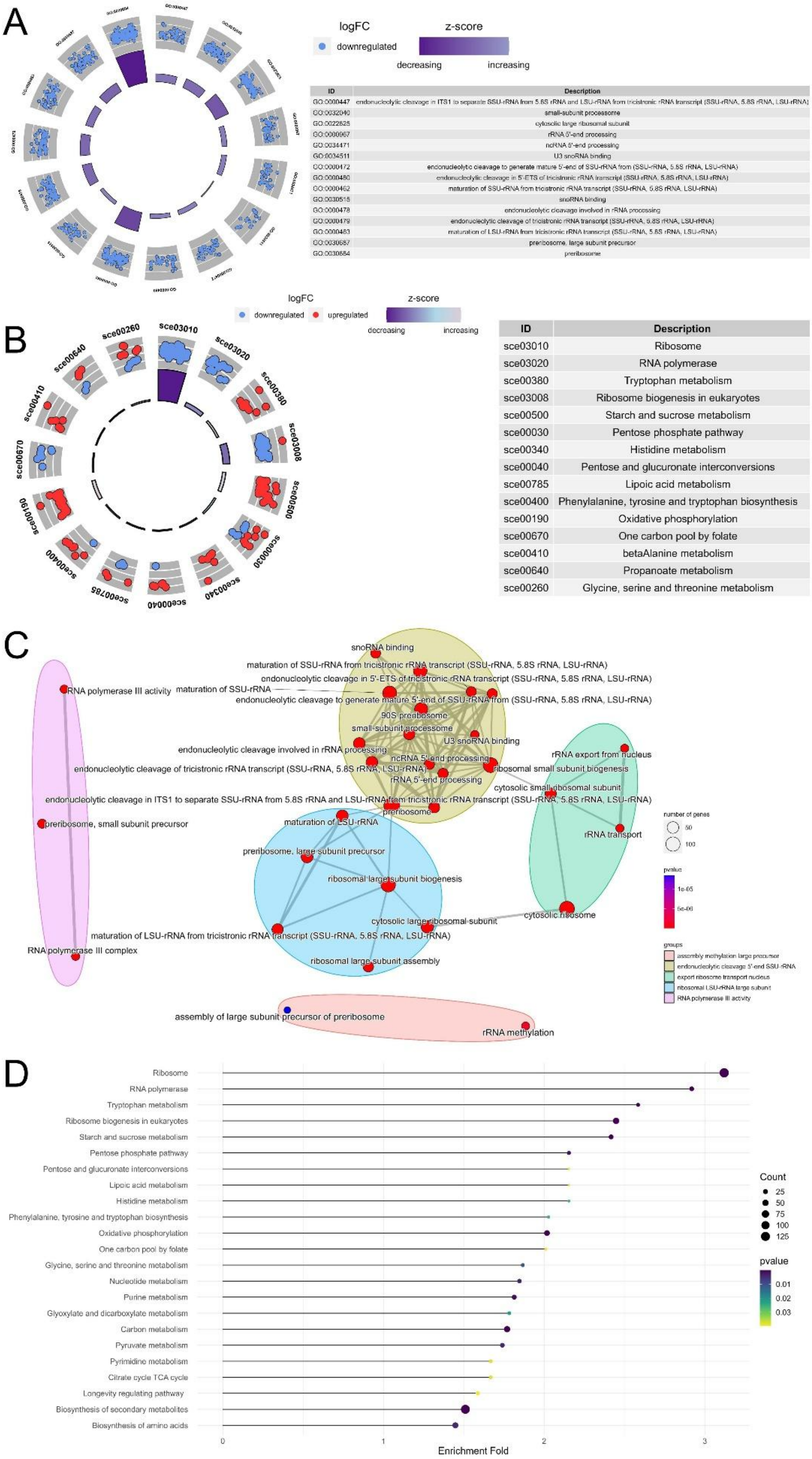
A
logFC
downregulated
z-score
decreasing
increasing
ID | Description
GO:0000447 | endonucleolytic cleavage in ITS1 to separate SSU-rRNA from 5.8S rRNA and LSU-rRNA from tricistronic rRNA transcript (SSU-rRNA, 5.8S rRNA, LSU-rRNA)
GO:0032040 | small-subunit processome
GO:0022625 | cytosolic large ribosomal subunit
GO:0000967 | rRNA 5'-end processing
GO:0034471 | ncRNA 5'-end processing
GO:0034511 | U3 snoRNA binding
GO:0000472 | endonucleolytic cleavage to generate mature 5'-end of SSU-rRNA from (SSU-rRNA, 5.8S rRNA, LSU-rRNA)
GO:0000480 | endonucleolytic cleavage in 5'-ETS of tricistronic rRNA transcript (SSU-rRNA, 5.8S rRNA, LSU-rRNA)
GO:0000462 | maturation of SSU-rRNA from tricistronic rRNA transcript (SSU-rRNA, 5.8S rRNA, LSU-rRNA)
GO:0030515 | snoRNA binding
GO:0000478 | endonucleolytic cleavage involved in rRNA processing
GO:0000479 | endonucleolytic cleavage of tricistronic rRNA transcript (SSU-rRNA, 5.8S rRNA, LSU-rRNA)
GO:0000463 | maturation of LSU-rRNA from tricistronic rRNA transcript (SSU-rRNA, 5.8S rRNA, LSU-rRNA)
GO:0030687 | preribosome, large subunit precursor
GO:0030684 | preribosome
B
logFC
downregulated
upregulated
z-score
decreasing
increasing
sce03010
sce03020
sce00380
sce03008
sce00500
sce00030
sce00340
sce00040
sce00785
sce00400
sce00190
sce00670
sce00410
sce00640
sce00260
ID | Description
sce03010 | Ribosome
sce03020 | RNA polymerase
sce00380 | Tryptophan metabolism
sce03008 | Ribosome biogenesis in eukaryotes
sce00500 | Starch and sucrose metabolism
sce00030 | Pentose phosphate pathway
sce00340 | Histidine metabolism
sce00040 | Pentose and glucuronate interconversions
sce00785 | Lipoic acid metabolism
sce00400 | Phenylalanine, tyrosine and tryptophan biosynthesis
sce00190 | Oxidative phosphorylation
sce00670 | One carbon pool by folate
sce00410 | betaAlanine metabolism
sce00640 | Propanoate metabolism
sce00260 | Glycine, serine and threonine metabolism
C
snoRNA binding
maturation of SSU-rRNA from tricistronic rRNA transcript (SSU-rRNA, 5.8S rRNA, LSU-rRNA)
endonucleolytic cleavage in 5'-ETS of tricistronic rRNA transcript (SSU-rRNA, 5.8S rRNA, LSU-rRNA)
RNA polymerase III activity
maturation of SSU-rRNA
endonucleolytic cleavage to generate mature 5'-end of SSU-rRNA from (SSU-rRNA, 5.8S rRNA, LSU-rRNA)
90S preribosome
small-subunit processome
U3 snoRNA binding
endonucleolytic cleavage involved in rRNA processing
ncRNA 5'-end processing
rRNA export from nucleus
endonucleolytic cleavage of tricistronic rRNA transcript (SSU-rRNA, 5.8S rRNA, LSU-rRNA)
ribosomal small subunit biogenesis
rRNA 5'-end processing
cytosolic small ribosomal subunit
endonucleolytic cleavage in ITS1 to separate SSU-rRNA from 5.8S rRNA and LSU-rRNA from tricistronic rRNA transcript (SSU-rRNA, 5.8S rRNA, LSU-rRNA)
preribosome
preribosome, small subunit precursor
maturation of LSU-rRNA
rRNA transport
preribosome, large subunit precursor
ribosomal large subunit biogenesis
cytosolic large ribosomal subunit
cytosolic ribosome
maturation of LSU-rRNA from tricistronic rRNA transcript (SSU-rRNA, 5.8S rRNA, LSU-rRNA)
RNA polymerase III complex
ribosomal large subunit assembly
assembly of large subunit precursor of preribosome
rRNA methylation
number of genes
50
100
pvalue
1e-05
5e-06
groups
assembly methylation large precursor
endonucleolytic cleavage 5'-end SSU-rRNA
export ribosome transport nucleus
ribosomal LSU-rRNA large subunit
RNA polymerase III activity
D
Ribosome
RNA polymerase
Tryptophan metabolism
Ribosome biogenesis in eukaryotes
Starch and sucrose metabolism
Pentose phosphate pathway
Pentose and glucuronate interconversions
Lipoic acid metabolism
Histidine metabolism
Phenylalanine, tyrosine and tryptophan biosynthesis
Oxidative phosphorylation
One carbon pool by folate
Glycine, serine and threonine metabolism
Nucleotide metabolism
Purine metabolism
Glyoxylate and dicarboxylate metabolism
Carbon metabolism
Pyruvate metabolism
Pyrimidine metabolism
Citrate cycle TCA cycle
Longevity regulating pathway
Biosynthesis of secondary metabolites
Biosynthesis of amino acids
0
1
2
3
Enrichment Fold
Count
25
50
75
100
125
pvalue
0.01
0.02
0.03

**Figure S9. Functional pathway enrichment associated with 2,5-anhydromannitol–induced transcriptional remodeling, Related to Figure 5**

(A) Circular plot showing Gene Ontology (GO) enrichment of downregulated genes following 2,5-anhydromannitol (2,5-AM) treatment.

(B) Circular plot showing GO enrichment of upregulated and downregulated pathways following 2,5-AM treatment.

(C) Network representation of enriched GO biological processes affected by 2,5-AM treatment.

(D) Dot plot of enriched KEGG pathways associated with genes differentially expressed following 2,5-AM treatment. Dot size indicates gene count and color represents adjusted P value.

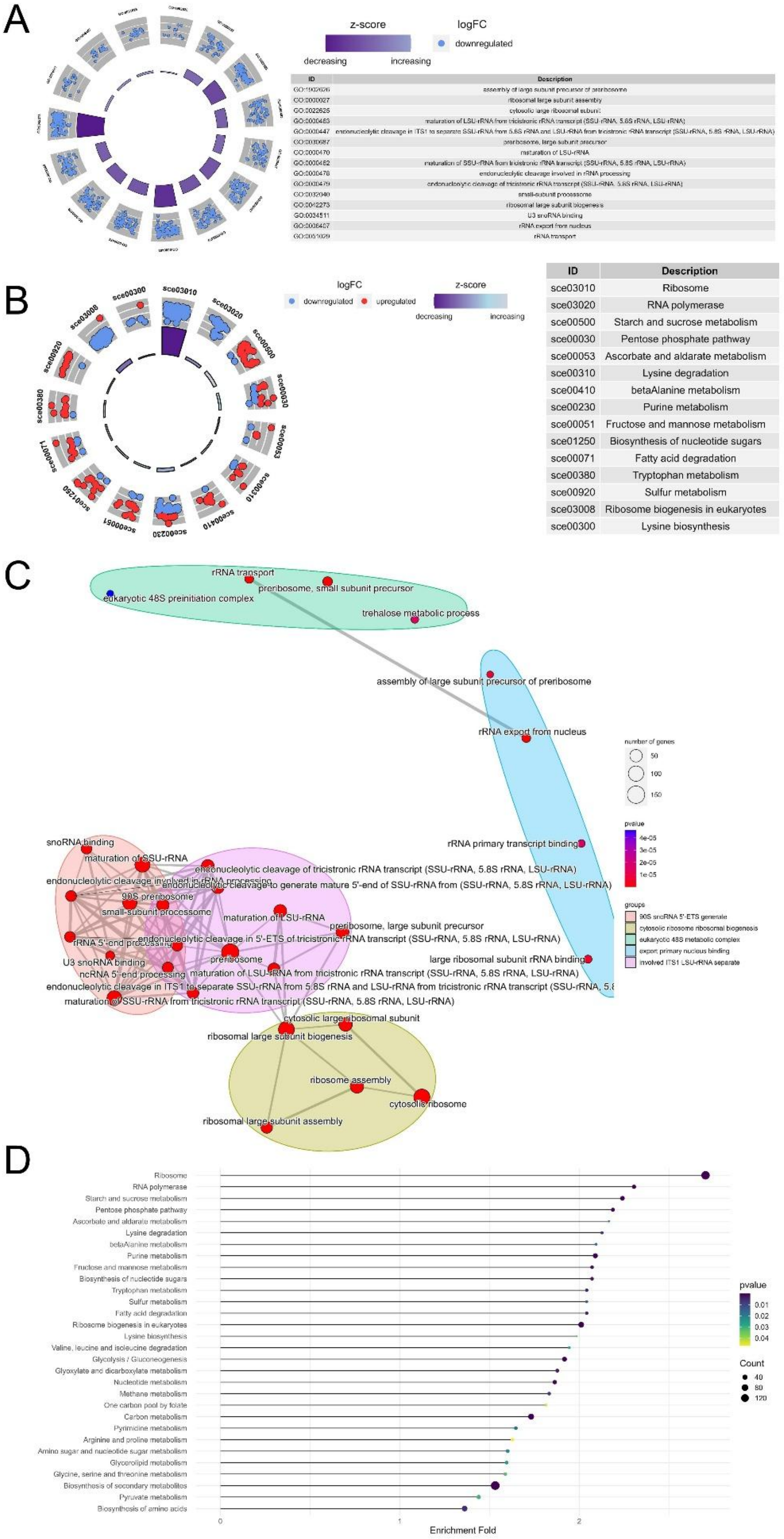
A
z-score
logFC
downregulated
decreasing
increasing
ID | Description
GO:1902626 | assembly of large subunit precursor of preribosome
GO:0000027 | ribosomal large subunit assembly
GO:0022625 | cytosolic large ribosomal subunit
GO:0000463 | maturation of LSU-rRNA from tricistronic rRNA transcript (SSU-rRNA, 5.8S rRNA, LSU-rRNA)
GO:0000447 | endonucleolytic cleavage in ITS1 to separate SSU-rRNA from 5.8S rRNA and LSU-rRNA from tricistronic rRNA transcript (SSU-rRNA, 5.8S rRNA, LSU-rRNA)
GO:0030687 | preribosome, large subunit precursor
GO:0000470 | maturation of LSU-rRNA
GO:0000462 | maturation of SSU-rRNA from tricistronic rRNA transcript (SSU-rRNA, 5.8S rRNA, LSU-rRNA)
GO:0000478 | endonucleolytic cleavage involved in rRNA processing
GO:0000479 | endonucleolytic cleavage of tricistronic rRNA transcript (SSU-rRNA, 5.8S rRNA, LSU-rRNA)
GO:0032040 | small-subunit processome
GO:0042273 | ribosomal large subunit biogenesis
GO:0034511 | U3 snoRNA binding
GO:0006407 | rRNA export from nucleus
GO:0051029 | rRNA transport
B
logFC
downregulated
upregulated
z-score
decreasing
increasing
sce03010
sce03020
sce00500
sce00030
sce00053
sce00410
sce00230
sce00051
sce01250
sce00071
sce00380
sce00920
sce03008
sce00300
ID | Description
sce03010 | Ribosome
sce03020 | RNA polymerase
sce00500 | Starch and sucrose metabolism
sce00030 | Pentose phosphate pathway
sce00053 | Ascorbate and aldarate metabolism
sce00310 | Lysine degradation
sce00410 | betaAlanine metabolism
sce00230 | Purine metabolism
sce00051 | Fructose and mannose metabolism
sce01250 | Biosynthesis of nucleotide sugars
sce00071 | Fatty acid degradation
sce00380 | Tryptophan metabolism
sce00920 | Sulfur metabolism
sce03008 | Ribosome biogenesis in eukaryotes
sce00300 | Lysine biosynthesis
C
rRNA transport
preribosome, small subunit precursor
eukaryotic 48S preinitiation complex
trehalose metabolic process
assembly of large subunit precursor of preribosome
rRNA export from nucleus
rRNA primary transcript binding
snoRNA binding
maturation of SSU-rRNA
endonucleolytic cleavage of tricistronic rRNA transcript (SSU-rRNA, 5.8S rRNA, LSU-rRNA)
endonucleolytic cleavage to generate mature 5'-end of SSU-rRNA from (SSU-rRNA, 5.8S rRNA, LSU-rRNA)
90S preribosome
small-subunit processome
maturation of LSU-rRNA
preribosome, large subunit precursor
endonucleolytic cleavage in 5'-ETS of tricistronic rRNA transcript (SSU-rRNA, 5.8S rRNA, LSU-rRNA)
preribosome
large ribosomal subunit rRNA binding
U3 snoRNA binding
ncRNA 5'-end processing
maturation of LSU-rRNA from tricistronic rRNA transcript (SSU-rRNA, 5.8S rRNA, LSU-rRNA)
maturation of SSU-rRNA from tricistronic rRNA transcript (SSU-rRNA, 5.8S rRNA, LSU-rRNA)
cytosolic large ribosomal subunit
ribosomal large subunit biogenesis
ribosome assembly
cytosolic ribosome
ribosomal large subunit assembly
number of genes
50
100
150
pvalue
4e-05
3e-05
2e-05
1e-05
groups
90S snoRNA 5'-ETS generate
cytosolic ribosome ribosomal biogenesis
eukaryotic 48S metabolic complex
export primary nucleus binding
involved ITS1 LSU-rRNA separate
D
Ribosome
RNA polymerase
Starch and sucrose metabolism
Pentose phosphate pathway
Ascorbate and aldarate metabolism
Lysine degradation
betaAlanine metabolism
Purine metabolism
Fructose and mannose metabolism
Biosynthesis of nucleotide sugars
Tryptophan metabolism
Sulfur metabolism
Fatty acid degradation
Ribosome biogenesis in eukaryotes
Lysine biosynthesis
Valine, leucine and isoleucine degradation
Glycolysis / Gluconeogenesis
Glyoxylate and dicarboxylate metabolism
Nucleotide metabolism
Methane metabolism
One carbon pool by folate
Carbon metabolism
Pyrimidine metabolism
Arginine and proline metabolism
Amino sugar and nucleotide sugar metabolism
Glycerolipid metabolism
Glycine, serine and threonine metabolism
Biosynthesis of secondary metabolites
Pyruvate metabolism
Biosynthesis of amino acids
0
1
2
Enrichment Fold
pvalue
0.01
0.02
0.03
0.04
Count
40
80
120

**Figure S10. Functional pathway enrichment associated with 2-deoxyglucose–induced transcriptional remodeling, Related to Figure 5**

(A) Circular plot showing Gene Ontology (GO) enrichment of downregulated genes following 2-deoxyglucose (2-DG) treatment.

(B) Circular plot showing GO enrichment of upregulated and downregulated pathways following 2-DG treatment.

(C) Network representation of enriched GO biological processes affected by 2-DG treatment.

(D) Dot plot of enriched KEGG pathways associated with genes differentially expressed following 2-DG treatment. Dot size indicates gene count and color represents adjusted P value.

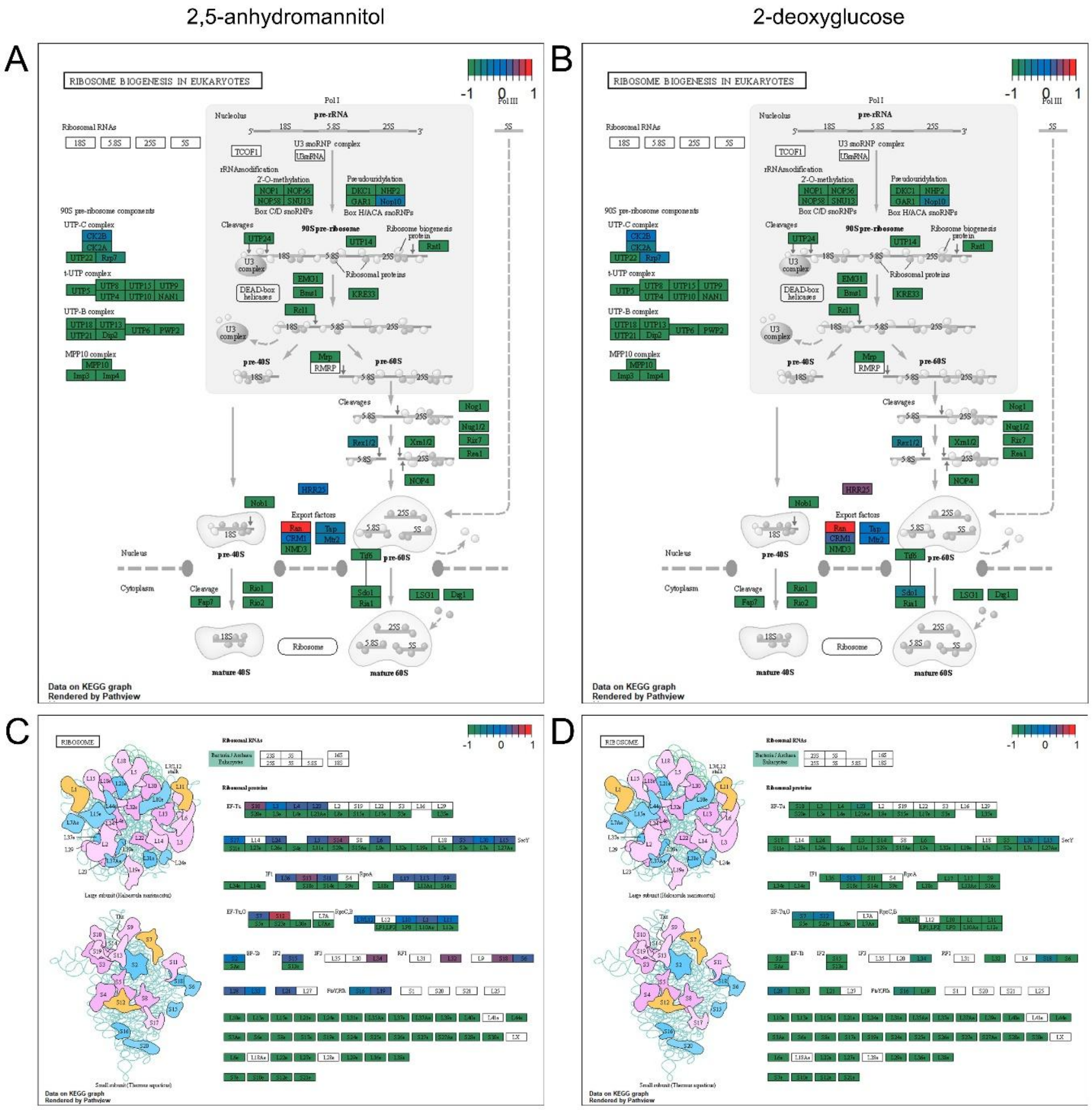


**Figure S11. Differential regulation of ribosome biogenesis and ribosomal pathways by glycolytic perturbations, Related to Figure 5**

(A–B) KEGG pathway maps of ribosome biogenesis in eukaryotes showing transcriptional changes induced by 2,5-anhydromannitol (2,5-AM) (A) or 2-deoxyglucose (2-DG) (B). Gene expression changes are overlaid onto pathway components, with color indicating relative fold change.

(C–D) KEGG pathway maps of the ribosome showing differential regulation of ribosomal protein genes following treatment with 2,5-AM (C) or 2-DG (D). Gene-level expression changes are indicated by the color scale.

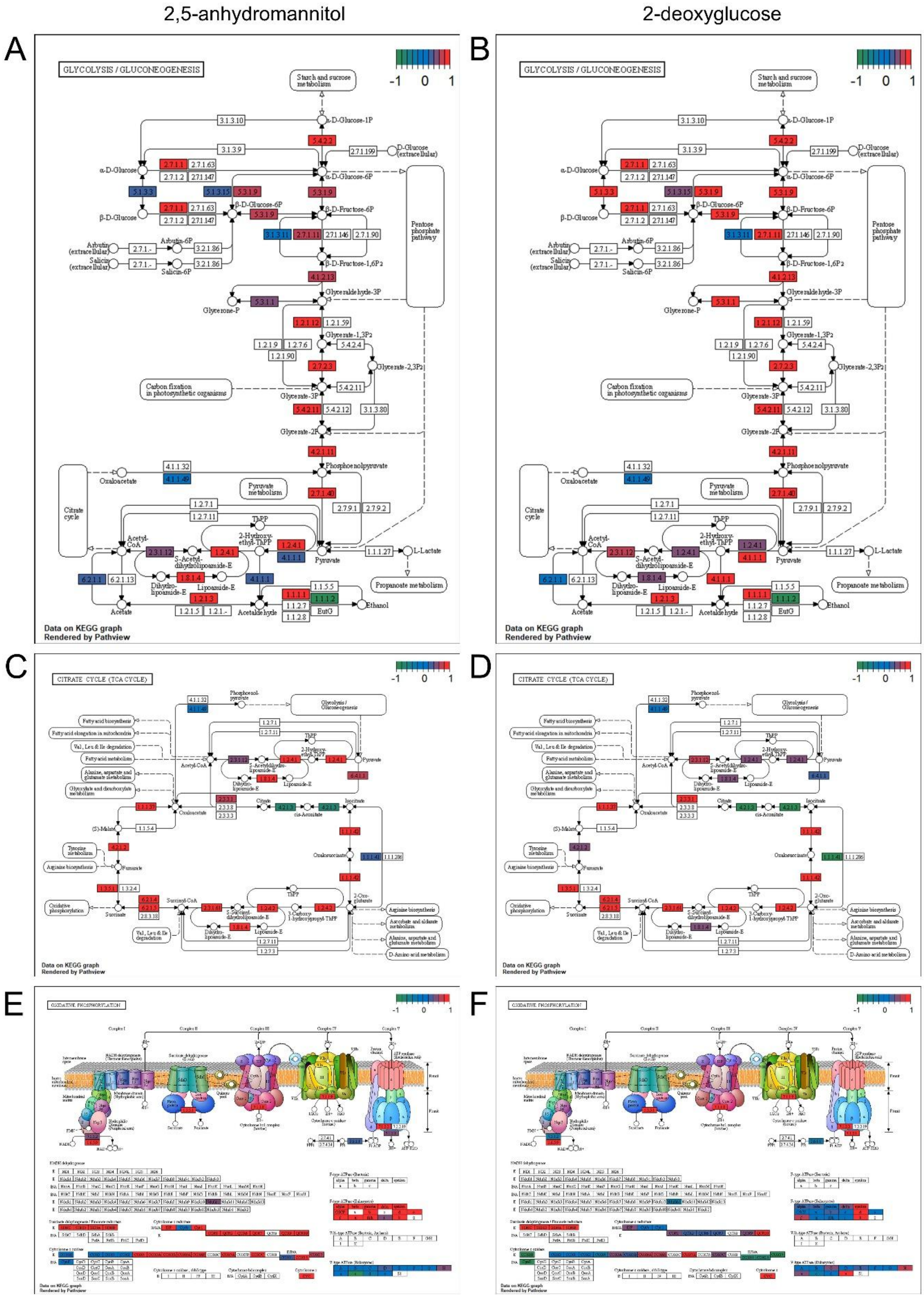
2,5-anhydromannitol
2-deoxyglucose
A
B
C
D
E
F
GLYCOLYSIS / GLUCONEOGENESIS
CITRATE CYCLE (TCA CYCLE)
OXIDATIVE PHOSPHORYLATION
-1 0 1
Data on KEGG graph
Rendered by Pathview

**Figure S12. Glycolytic perturbations differentially regulate central carbon metabolism and mitochondrial pathways, Related to Figure 5**

(A–B) KEGG pathway maps of glycolysis/gluconeogenesis showing transcriptional changes following treatment with 2,5-anhydromannitol (2,5-AM) (A) or 2-deoxyglucose (2-DG) (B). Differentially expressed genes are overlaid onto pathway components, with colors indicating relative log2 fold change.

(C–D) KEGG pathway maps of the tricarboxylic acid (TCA) cycle showing transcriptional changes induced by 2,5-AM (C) or 2-DG (D).

(E–F) KEGG pathway maps of oxidative phosphorylation illustrating transcriptional regulation of mitochondrial respiratory complexes following 2,5-AM (E) or 2-DG (F) treatment.

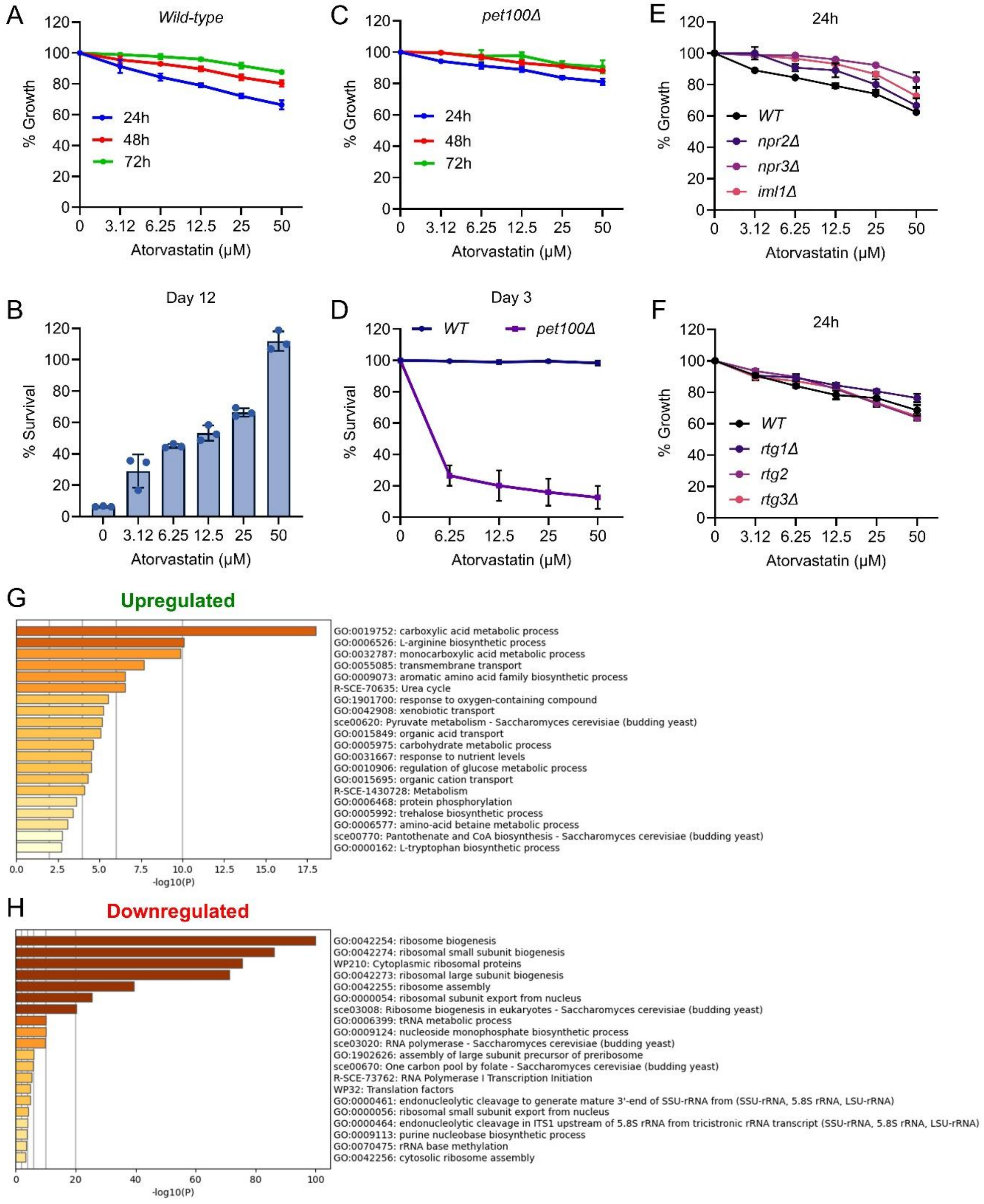


**Figure S13. Sterol perturbation regulates growth and survival in a mitochondrial- and TORC1-dependent manner, Related to Figure 5**

(A) Relative growth of wild-type cells treated with increasing concentrations of atorvastatin, measured at 24 h, 48 h, and 72 h. Growth is expressed as percentage relative to untreated controls, mean ± SD (n = 3).

(B) Quantification of cell survival of wild-type cells on day 12 of stationary phase following treatment with increasing concentrations of atorvastatin. Survival is expressed as percentage relative to untreated controls, mean ± SD (n = 3).

(C) Relative growth of mitochondrial-deficient *pet100Δ* cells treated with increasing concentrations of atorvastatin, measured at 24 h, 48 h, and 72 h, mean ± SD (n = 3).

(D) Survival of wild-type and *pet100Δ* cells on day 3 of stationary phase following atorvastatin treatment, mean ± SD (n = 3).

(E) Relative growth of wild-type and SEACIT mutants (*npr2Δ*, *npr3Δ*, and *iml1Δ*) treated with atorvastatin for 24 h, expressed as percentage relative to untreated controls, mean ± SD (n = 3).

(F) Relative growth of wild-type and retrograde signaling mutants (*rtg1Δ*, *rtg2Δ*, and *rtg3Δ*) treated with atorvastatin for 24 h, mean ± SD (n = 3).

(G, H) Gene Ontology (GO) enrichment analysis of genes upregulated (G) and downregulated (H) following atorvastatin treatment. Bars represent –log10(P value).

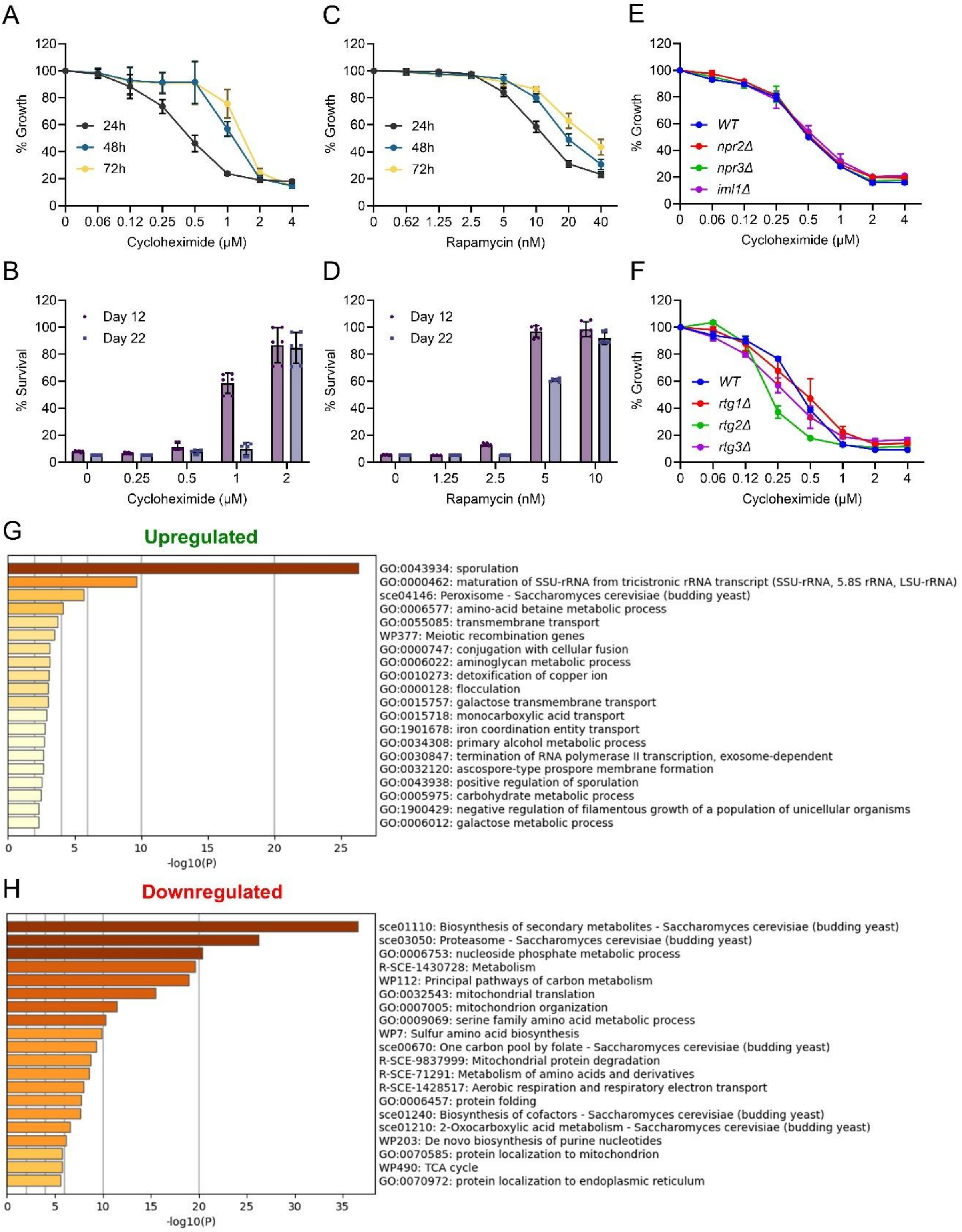


**Figure S14. Translation inhibition promotes survival through a distinct mechanism from TORC1-mediated metabolic regulation, Related to Figure 5**

(A) Relative growth of wild-type cells treated with increasing concentrations of cycloheximide (CHX), measured at 24 h, 48 h, and 72 h. Growth is expressed as percentage relative to untreated controls, mean ± SD (n =6).

(B) Quantification of survival of wild-type cells following CHX treatment, assessed on day 12 and day 22 of stationary phase. Survival is expressed as percentage relative to untreated controls, mean ± SD (n =6).

(C) Relative growth of wild-type cells treated with increasing concentrations of rapamycin, measured at 24 h, 48 h, and 72 h. Data are shown as mean ± SD (n =6).

(D) Quantification of survival of wild-type cells following rapamycin treatment, assessed on day 12 and day 22 of stationary phase, mean ± SD (n =6).

(E) Relative growth of wild-type and SEACIT mutants (*npr2Δ*, *npr3Δ*, and *iml1Δ*) treated with increasing concentrations of CHX for 24 h, expressed as percentage relative to untreated controls, mean ± SD (n =3).

(F) Relative growth of wild-type and retrograde signaling mutants (*rtg1Δ*, *rtg2Δ*, and *rtg3Δ*) treated with increasing concentrations of CHX for 24 h, expressed as percentage relative to untreated controls, mean ± SD (n =3).

(G, H) Gene Ontology (GO) enrichment analysis of genes upregulated (G) and downregulated (H) following CHX treatment. Bars represent –log10(P value).

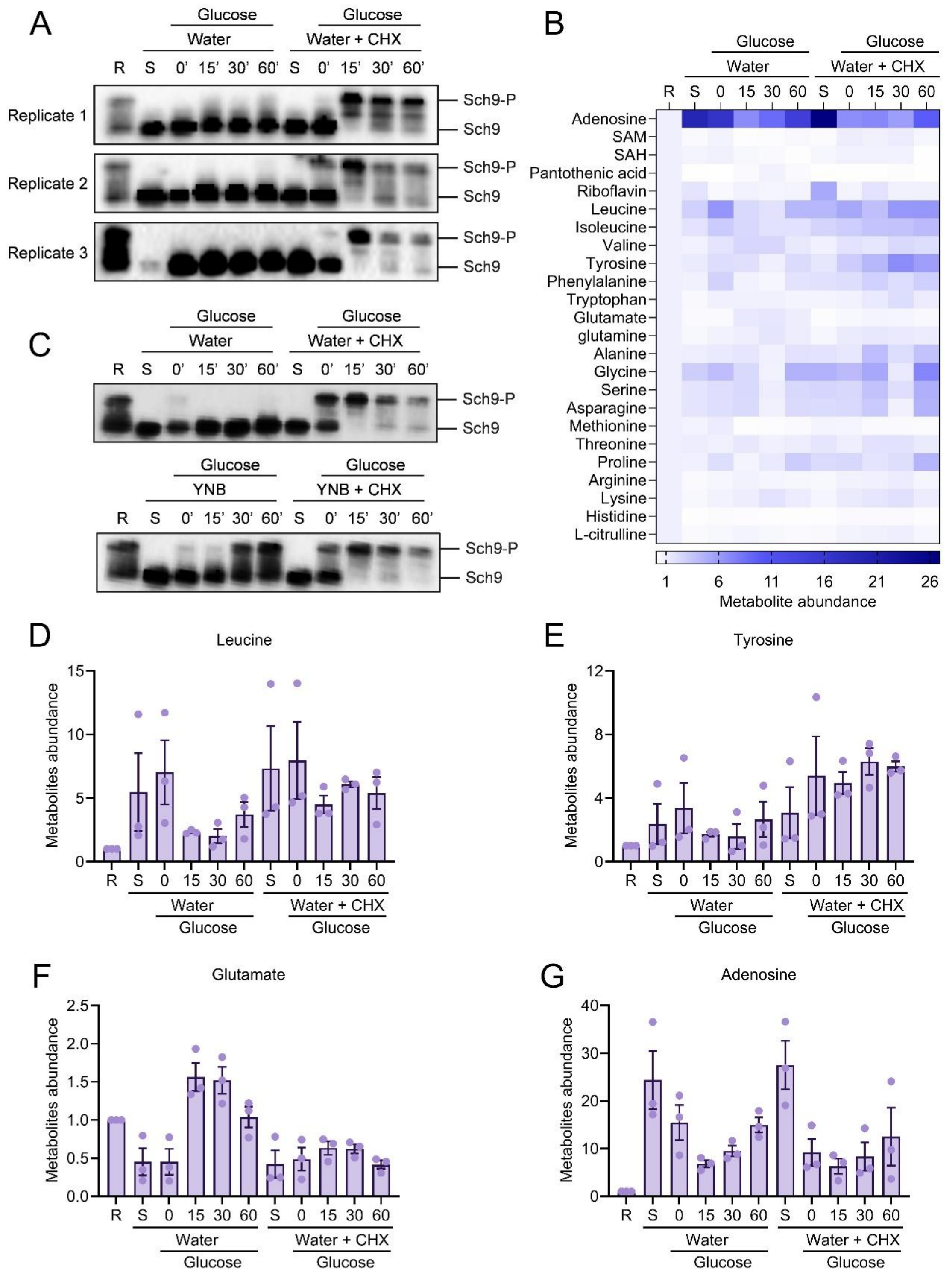


**Figure S15. Translation inhibition alters TORC1 reactivation and metabolic responses upon nutrient repletion, Related to Figure 5**

(A) Immunoblot analysis of TORC1 activity assessed by phosphorylation-dependent mobility shift of Sch9 (Sch9-P) following glucose re-addition. Prototrophic CEN.PK cells expressing Sch9-6×HA were analyzed under two conditions: glucose re-addition after water starvation (Water) or glucose re-addition following water starvation in the presence of cycloheximide (Water + CHX). Samples were collected at the indicated time points (0, 15, 30, and 60 min).

R, exponentially growing cells in synthetic defined (SD) medium; S, water-starved cells prior to glucose re-addition. Three independent biological replicates are shown.

(B) Heatmap showing relative intracellular metabolite abundance following glucose re-addition after water starvation (Water) or water starvation with cycloheximide treatment (Water + CHX). Metabolites include amino acids, nucleotide-related metabolites, and cofactors. Values are shown as fold change relative to exponentially growing wild-type cells (R, set to 1).

(C) Immunoblot analysis of Sch9 phosphorylation following glucose re-addition in cells starved in water or yeast nitrogen base (YNB), with or without cycloheximide (CHX), collected at the indicated time points. R and S denote exponential and starved states, respectively.

D–G) Quantification of selected representative metabolites derived from the dataset shown in (B) following glucose re-addition after water starvation or water starvation with CHX treatment: leucine (D), tyrosine (E), glutamate (F), and adenosine (G). Metabolite abundance is shown as fold change relative to exponentially growing cells (R, set to 1). Bars represent mean ± SEM with individual biological replicates indicated.

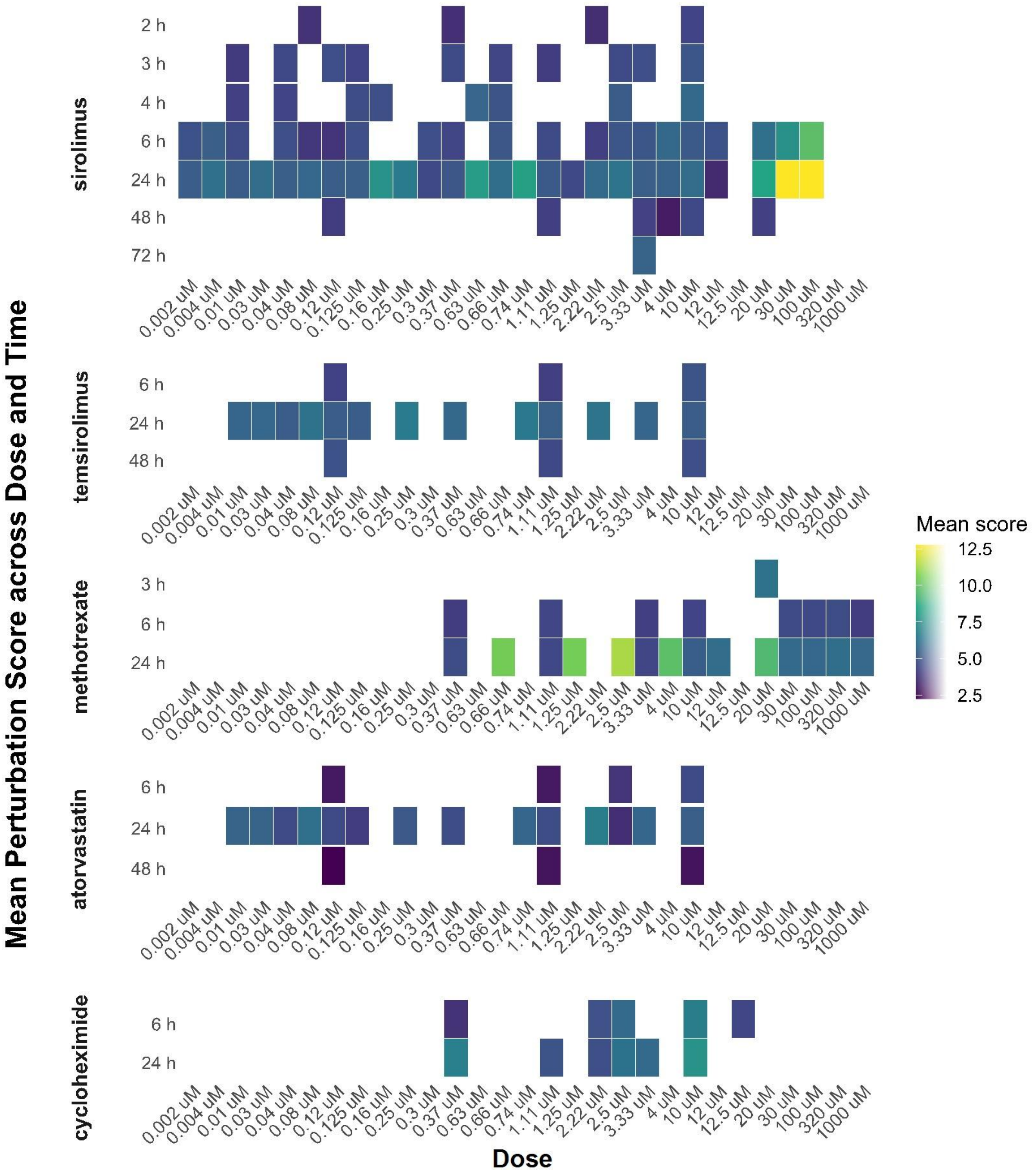


**Figure S16. Dose- and time-dependent transcriptional perturbation signatures identified using CMAP analysis, Related to Figure 6**

Heatmaps show mean perturbation scores derived from Connectivity Map (CMAP) RNA-seq signatures following treatment with sirolimus, temsirolimus, methotrexate, atorvastatin, and cycloheximide across the indicated doses (μM) and time points (2–72 h). Colors represent the magnitude of transcriptional perturbation (low to high). Blank spaces indicate conditions for which RNA-seq data were not available.

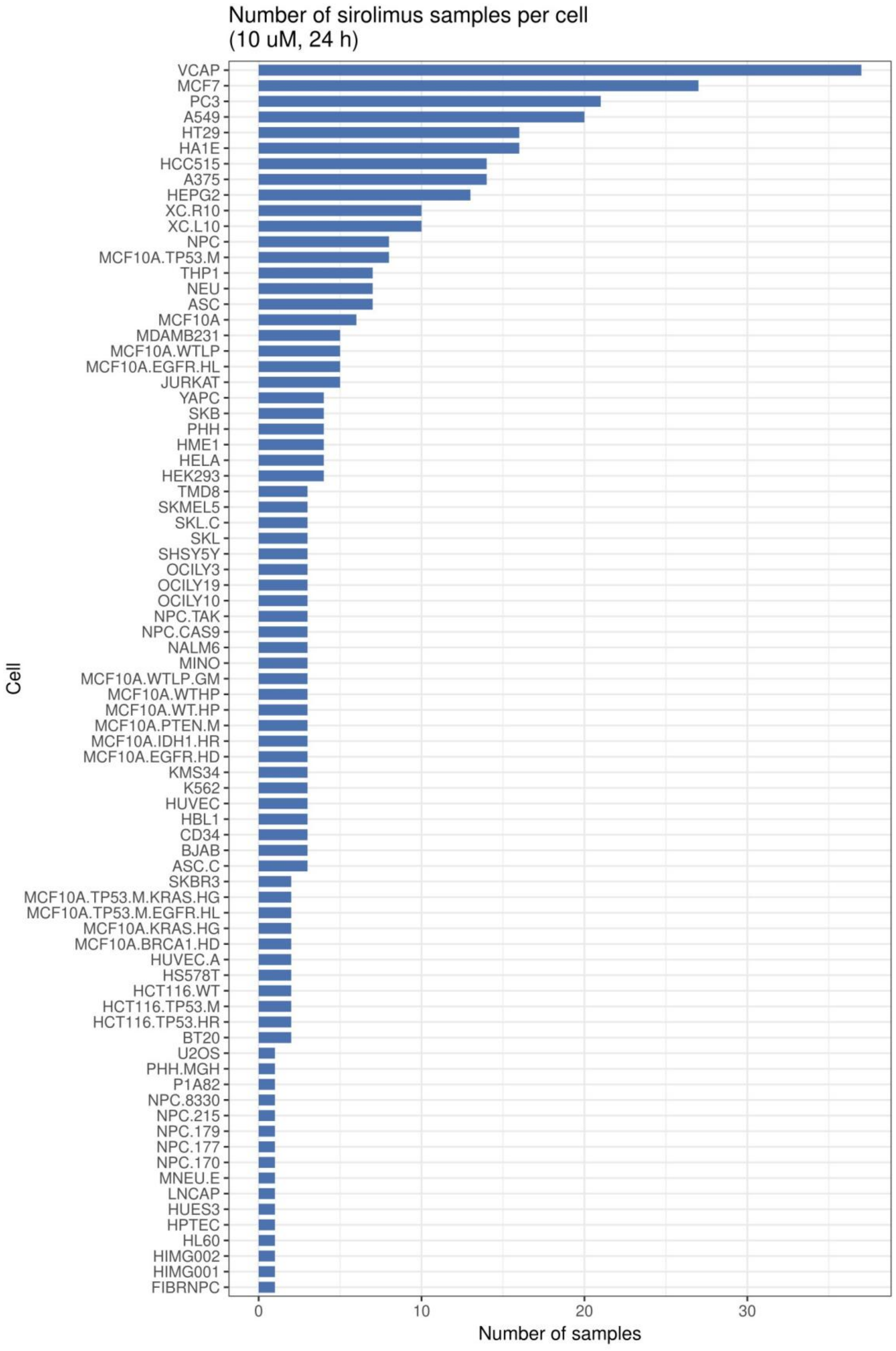


**Figure S17. Sirolimus RNA-seq samples across cell lines in CMAP, Related to Figure 6**

Bar plot showing the number of RNA-seq samples treated with sirolimus (10 µM, 24 h) across different cell lines in the Connectivity Map (CMAP) dataset.

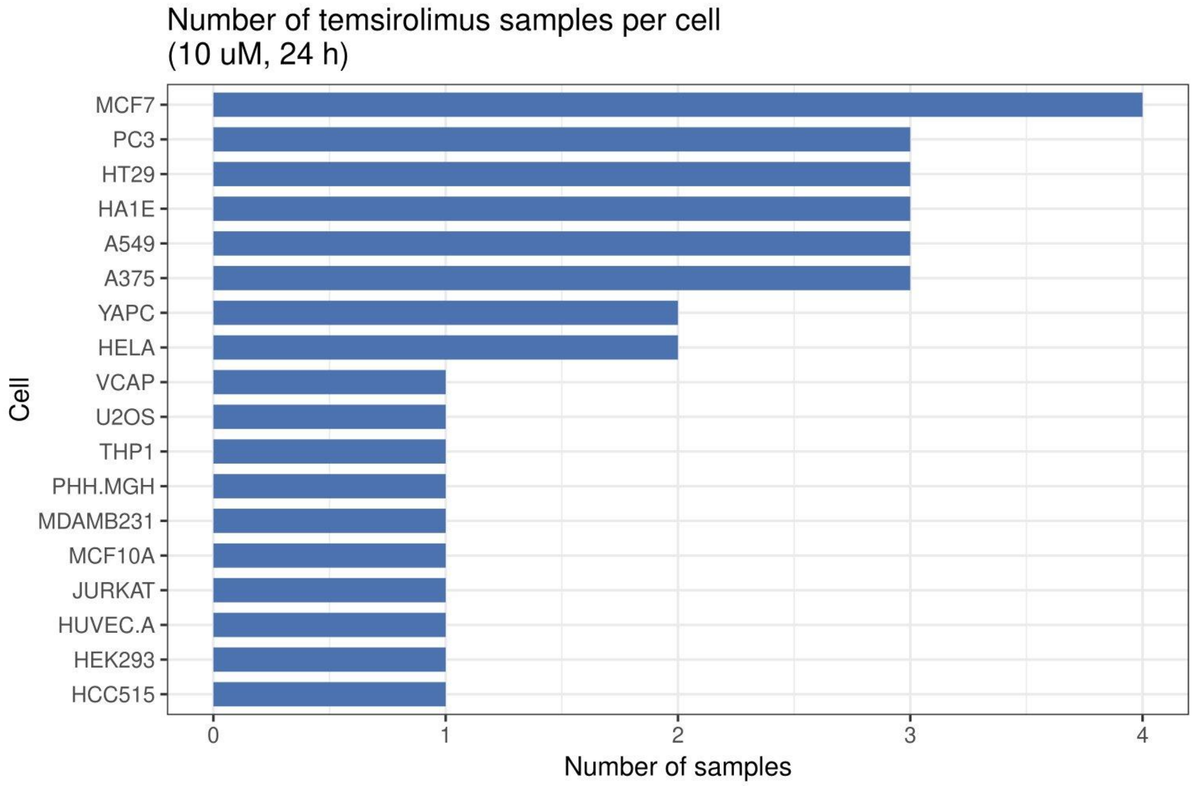


**Figure S18. Temsirolimus RNA-seq samples across cell lines in CMAP, Related to Figure 6**

Bar plot showing the number of RNA-seq samples treated with temsirolimus (10 μM, 24 h) across different cell lines in the Connectivity Map (CMAP) dataset.

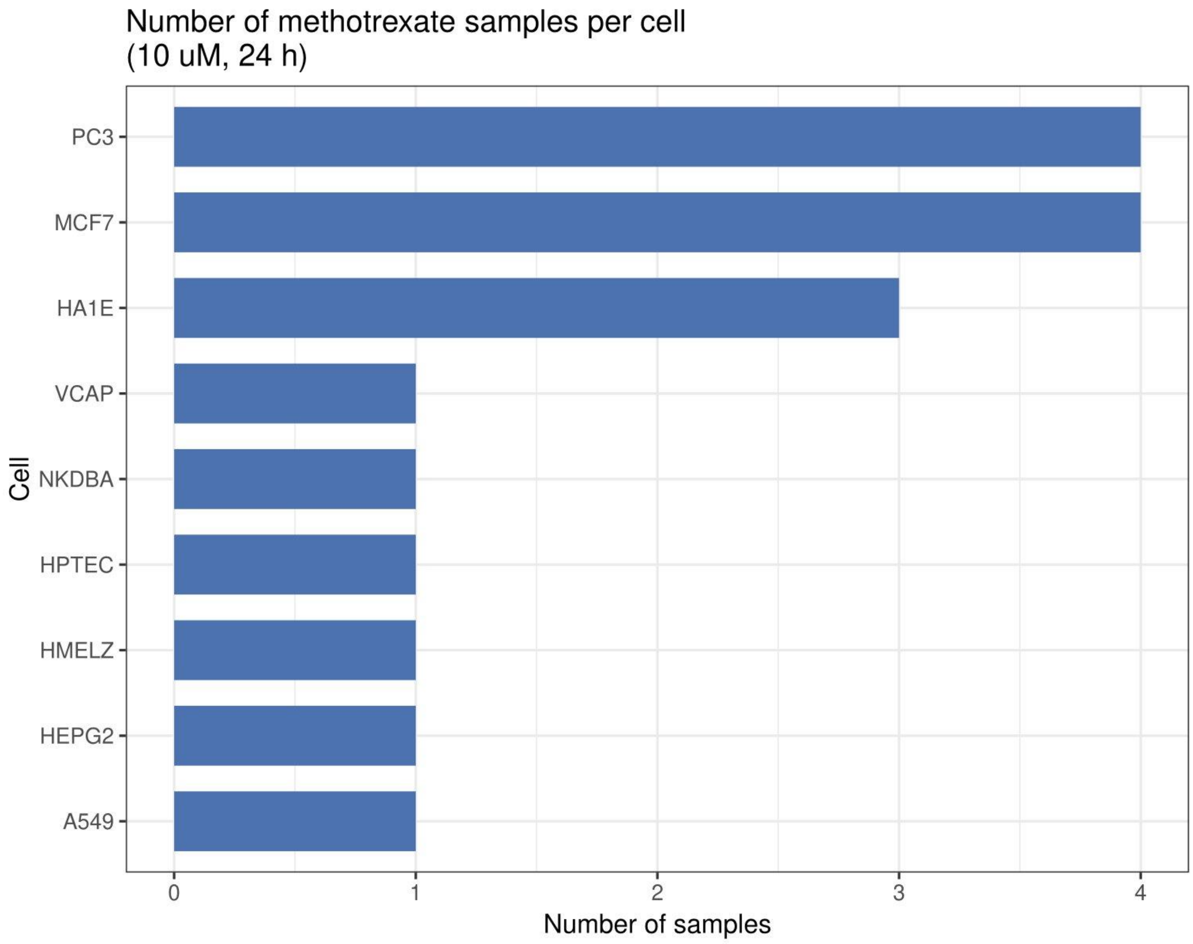


**Figure S19. Methotrexate RNA-seq samples across cell lines in CMAP, Related to Figure 6**

Bar plot showing the number of RNA-seq samples treated with methotrexate (10 μM, 24 h) across different cell lines in the Connectivity Map (CMAP) dataset.

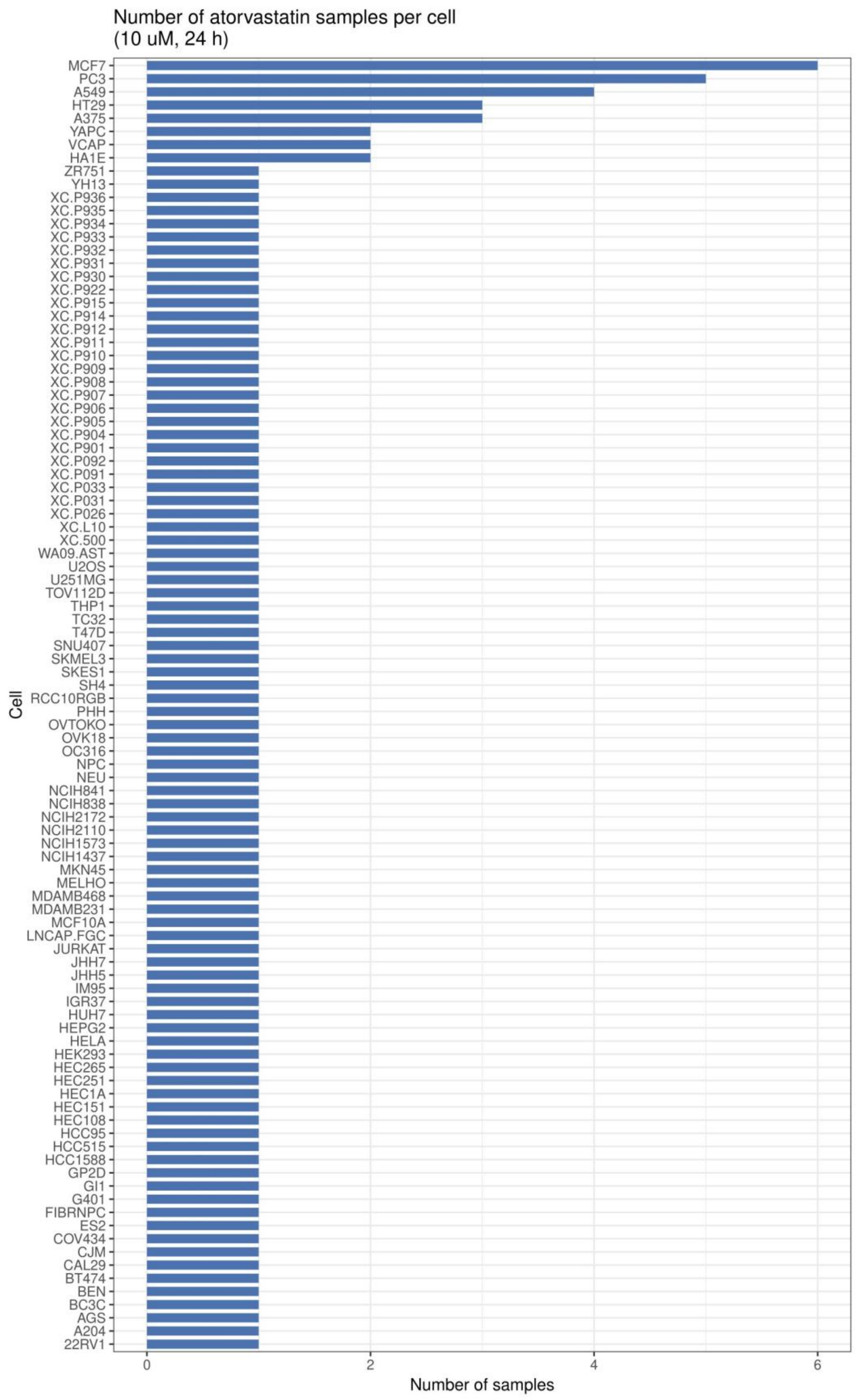


**Figure S20. Atorvastatin RNA-seq samples across cell lines in CMAP, Related to Figure 6**
Bar plot showing the number of RNA-seq samples treated with atorvastatin (10 μM, 24 h) across different cell lines in the Connectivity Map (CMAP) dataset.

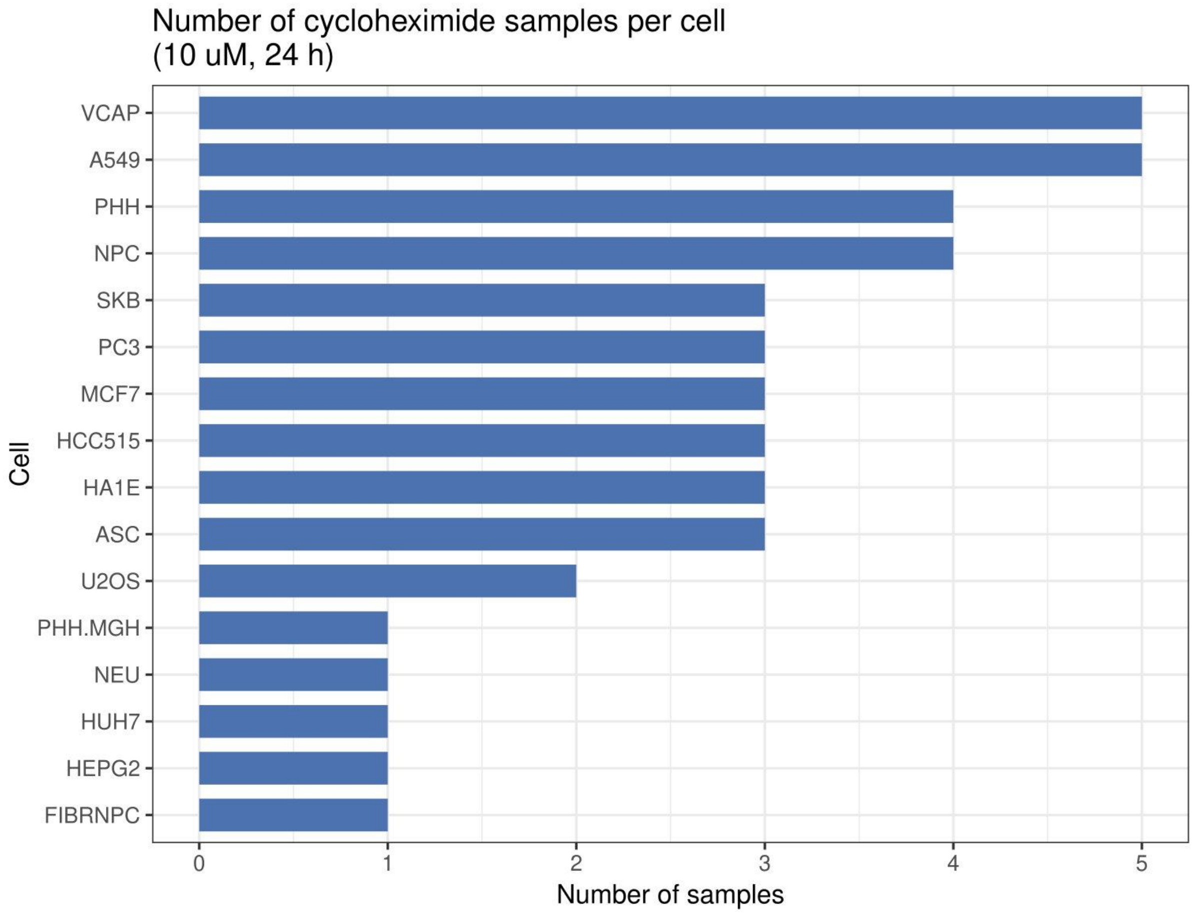


**Figure S21. Cycloheximide RNA-seq samples across cell lines in CMAP, Related to Figure 6**

Bar plot showing the number of RNA-seq samples treated with cycloheximide (10 μM, 24 h) across different cell lines in the Connectivity Map (CMAP) dataset.

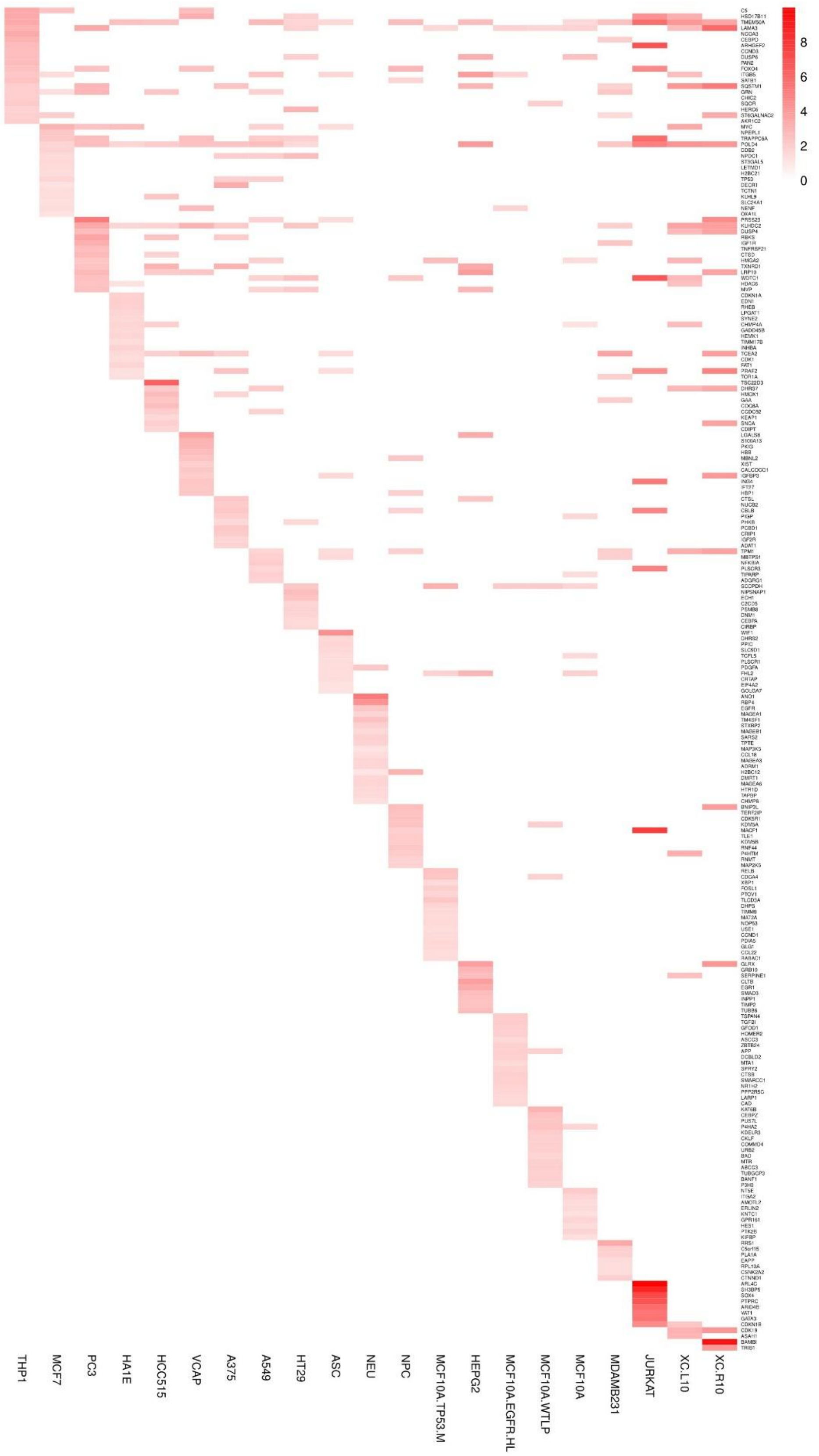


**Figure S22. Sirolimus -induced upregulated gene signatures across cell lines, Related to Figure 6**

Heatmap comparing the top 20 genes upregulated by sirolimus across indicated cell lines based on CMAP transcriptional profiles.

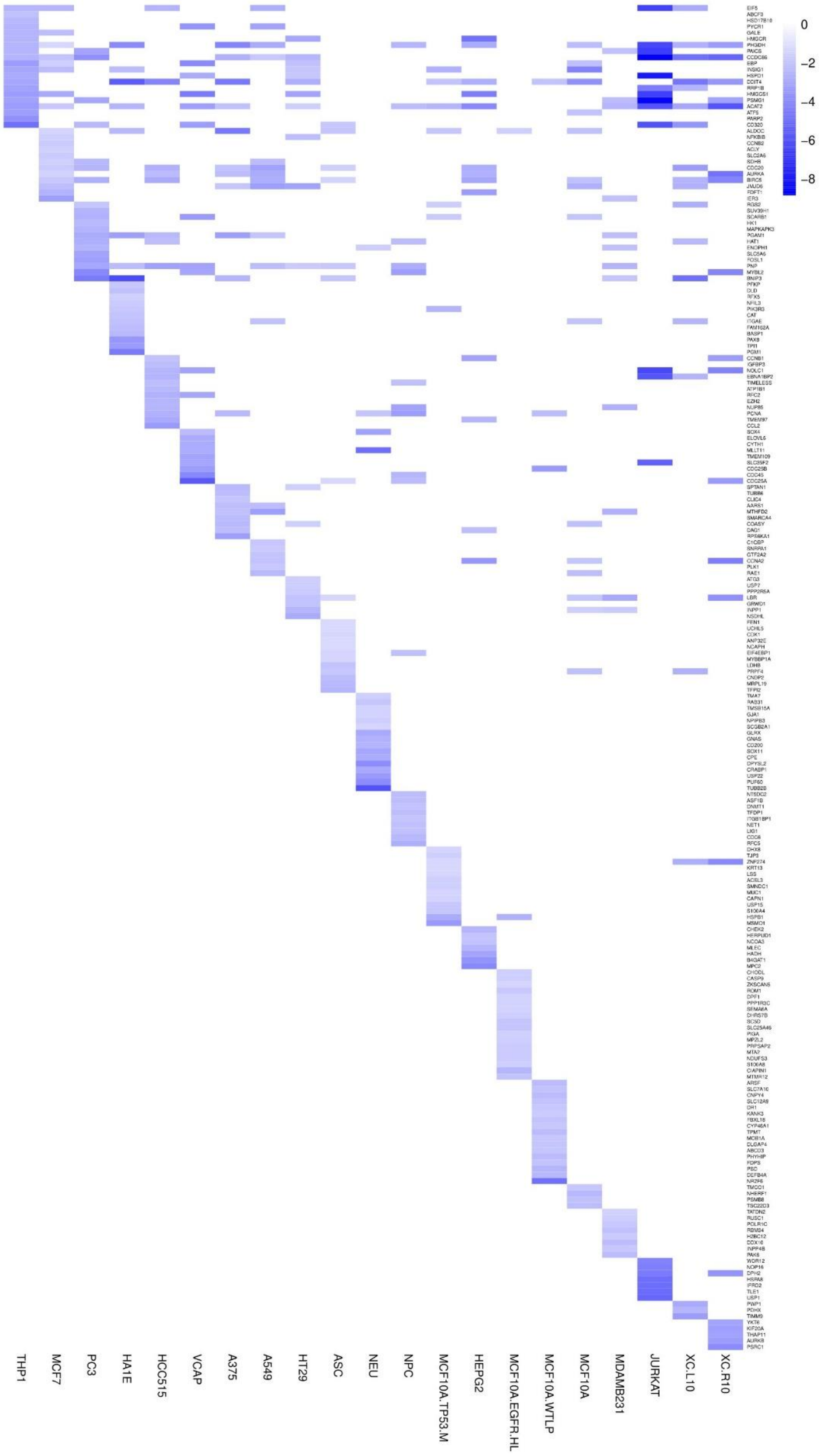


**Figure S23. Sirolimus -induced downregulated gene signatures across cell lines, Related to Figure 6**

Heatmap comparing the top 20 genes downregulated by sirolimus across indicated cell lines based on CMAP transcriptional profiles.

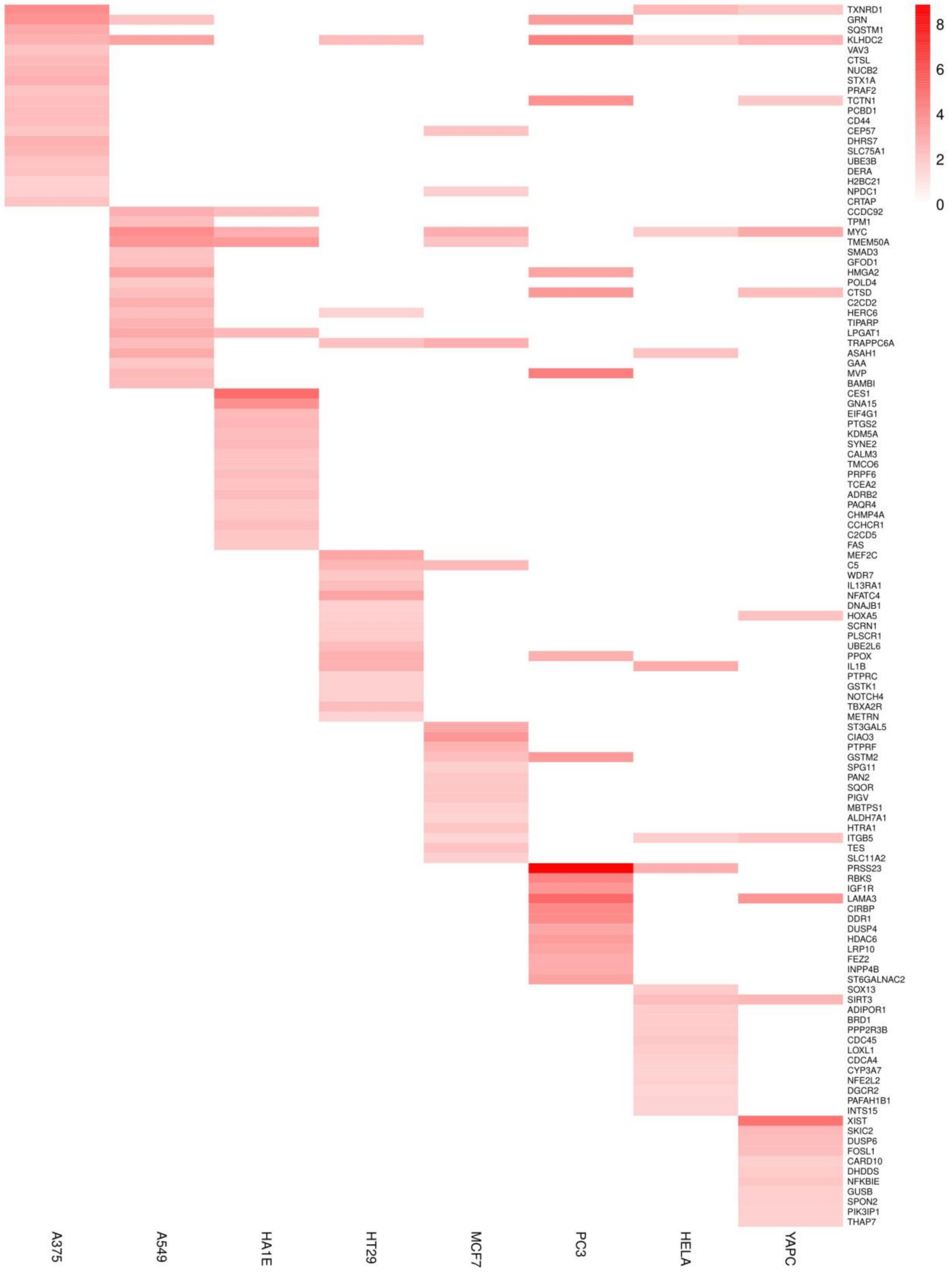


**Figure S24. Temsirolimus -induced upregulated gene signatures across cell lines, Related to Figure 6**

Heatmap comparing the top 20 genes upregulated by temsirolimus across indicated cell lines based on CMAP transcriptional profiles.

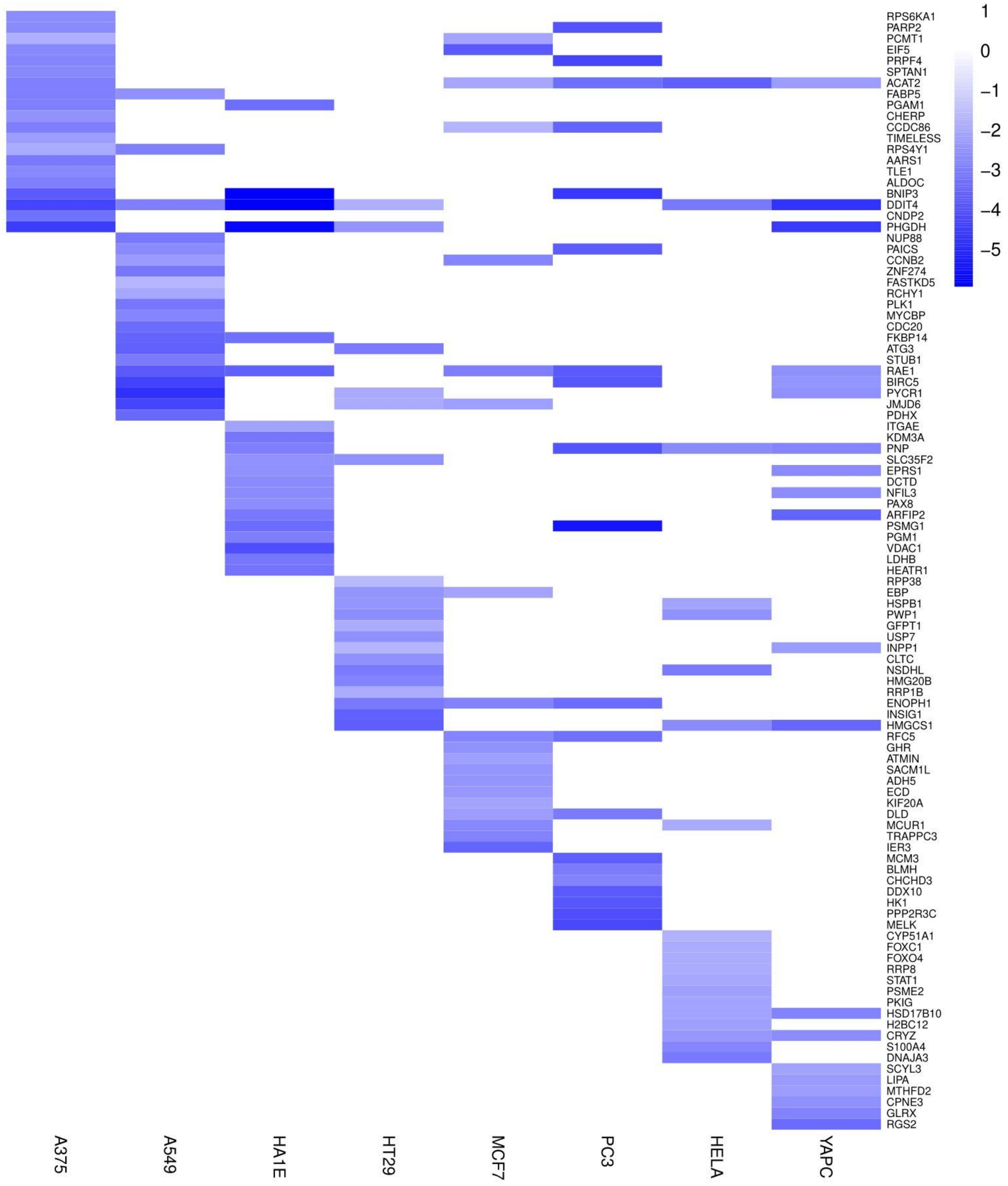


**Figure S25. Temsirolimus -induced downregulated gene signatures across cell lines, Related to Figure 6**
Heatmap comparing the top 20 genes downregulated by temsirolimus across indicated cell lines based on CMAP transcriptional profiles.

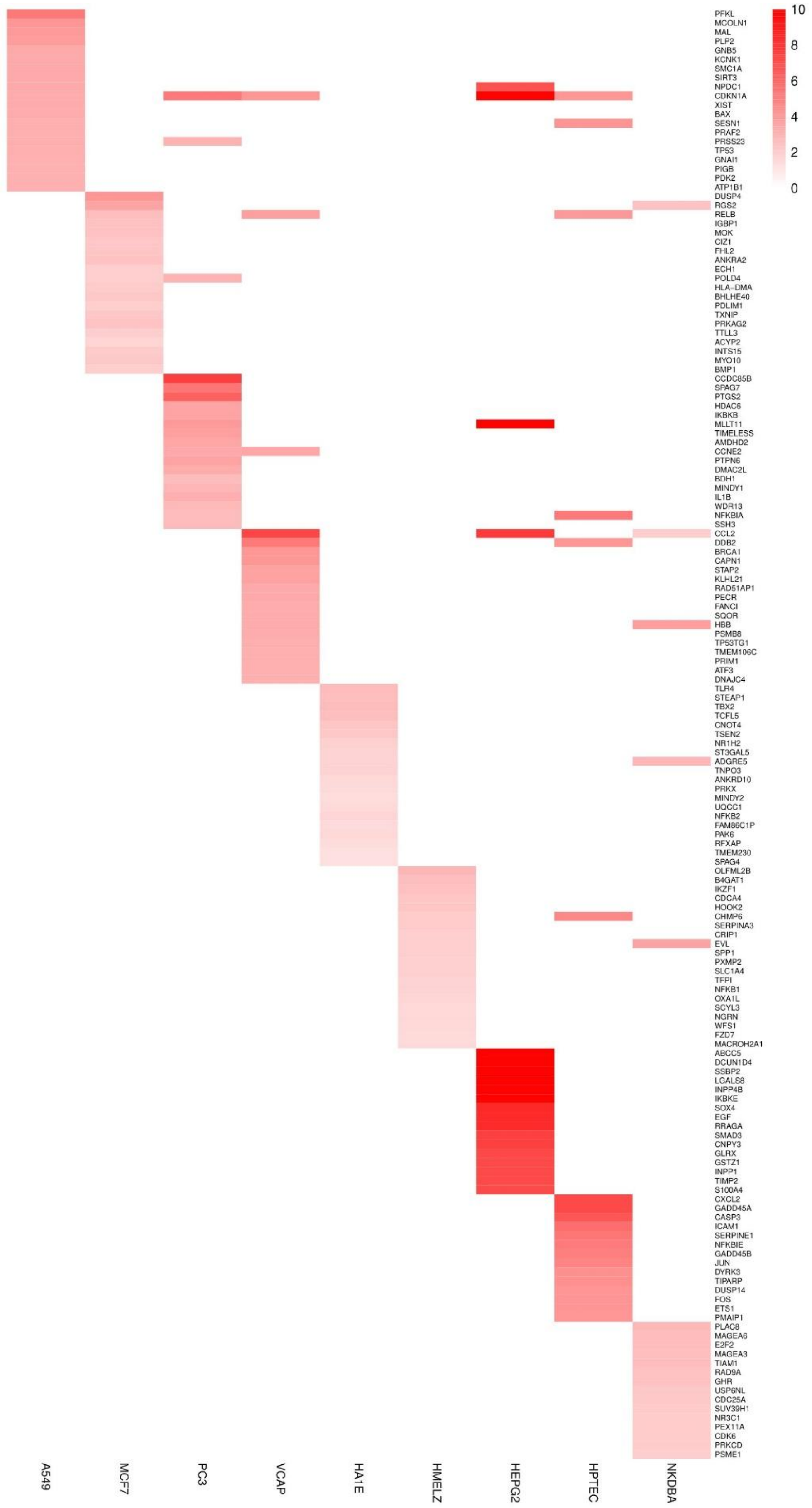


**Figure S26. Methotrexate -induced upregulated gene signatures across cell lines, Related to Figure 6**

Heatmap comparing the top 20 genes upregulated by methotrexate across indicated cell lines based on CMAP transcriptional profiles.

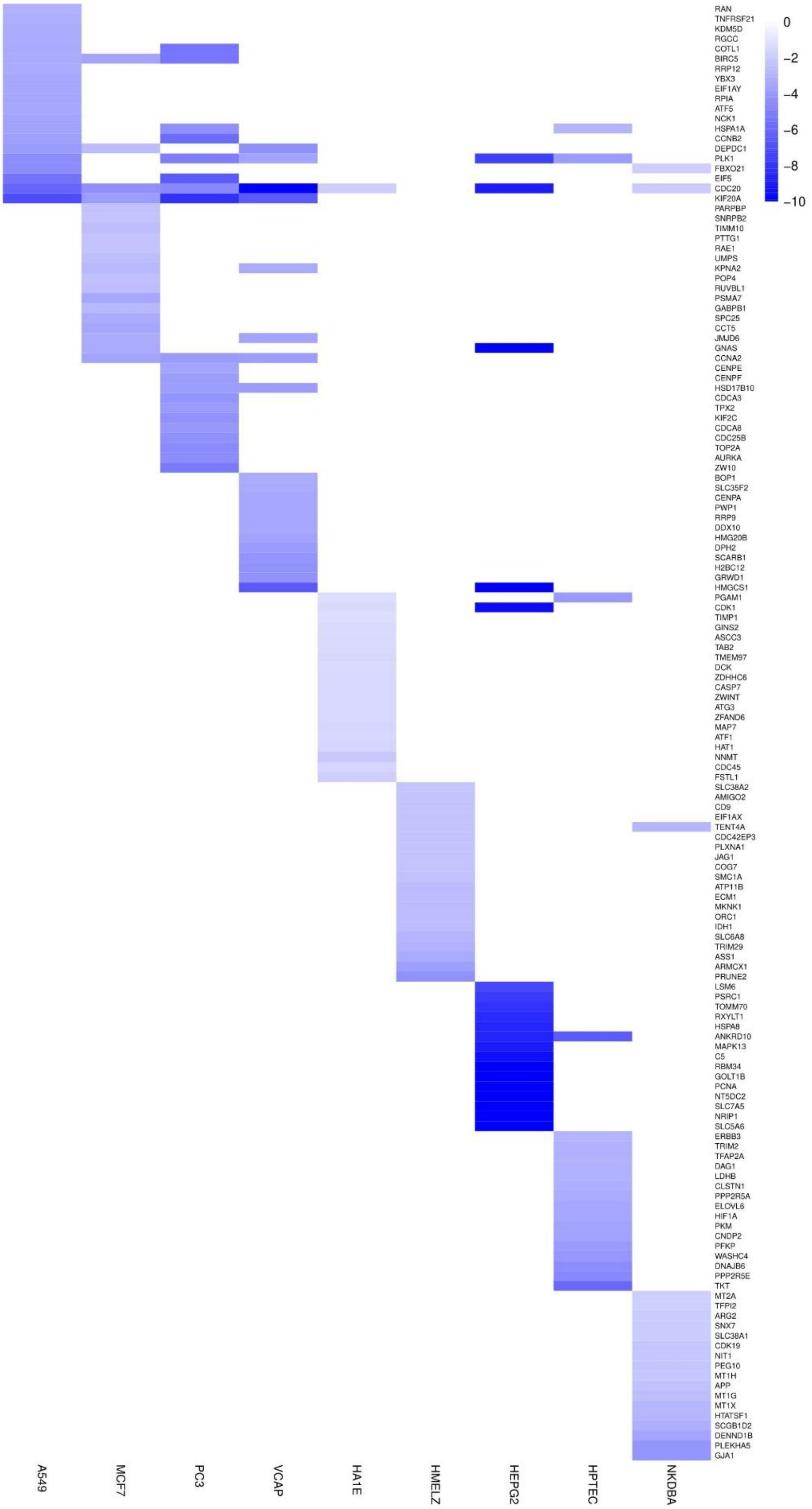


**Figure S27. Methotrexate -induced downregulated gene signatures across cell lines, Related to Figure 6**

Heatmap comparing the top 20 genes downregulated by methotrexate across indicated cell lines based on CMAP transcriptional profiles.

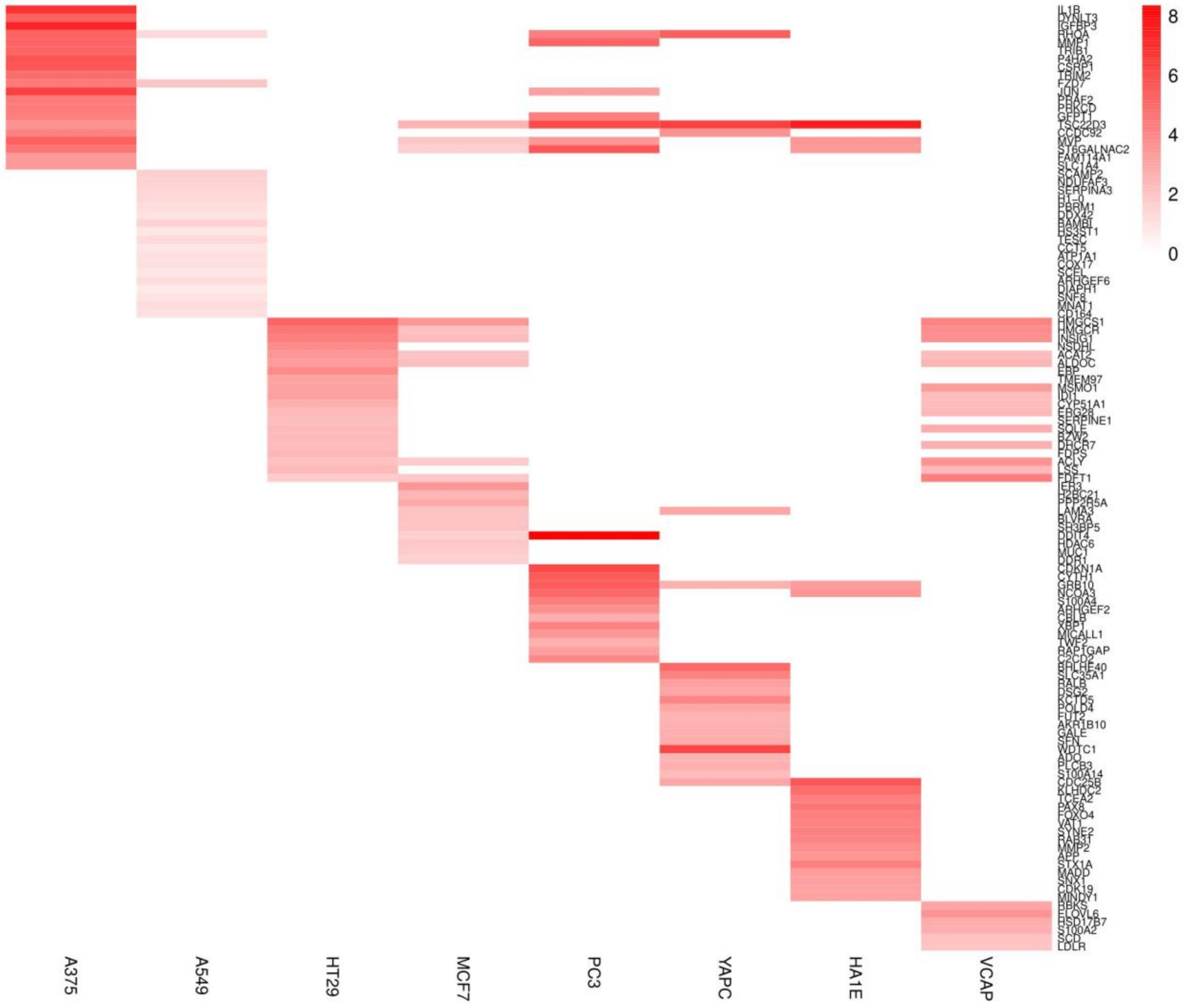


**Figure S28. Atorvastatin -induced upregulated gene signatures across cell lines, Related to Figure 6**

Heatmap comparing the top 20 genes upregulated by atorvastatin across indicated cell lines based on CMAP transcriptional profiles.

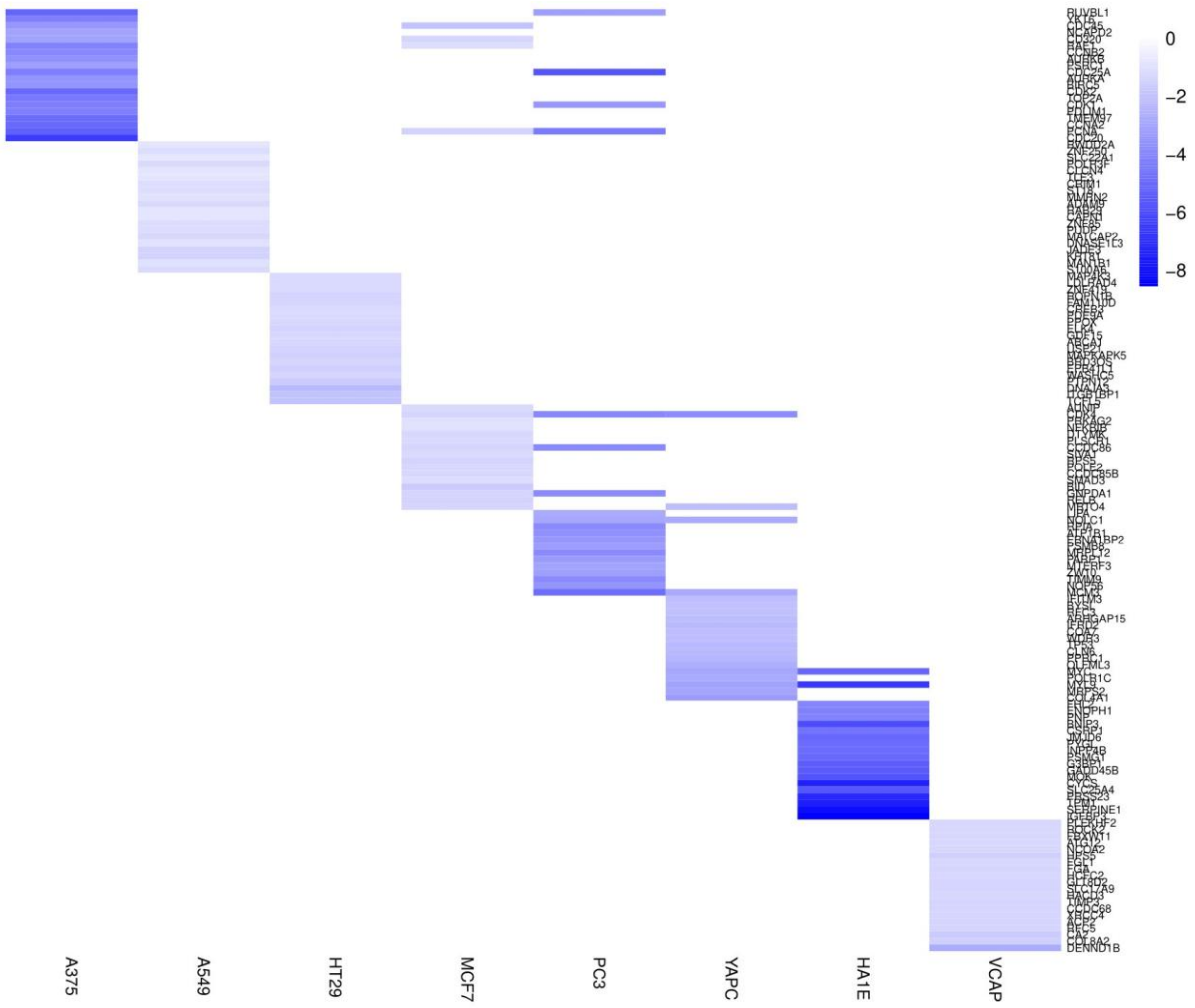


**Figure S29. Atorvastatin -induced downregulated gene signatures across cell lines, Related to Figure 6**

Heatmap comparing the top 20 genes downregulated by atorvastatin across indicated cell lines based on CMAP transcriptional profiles.

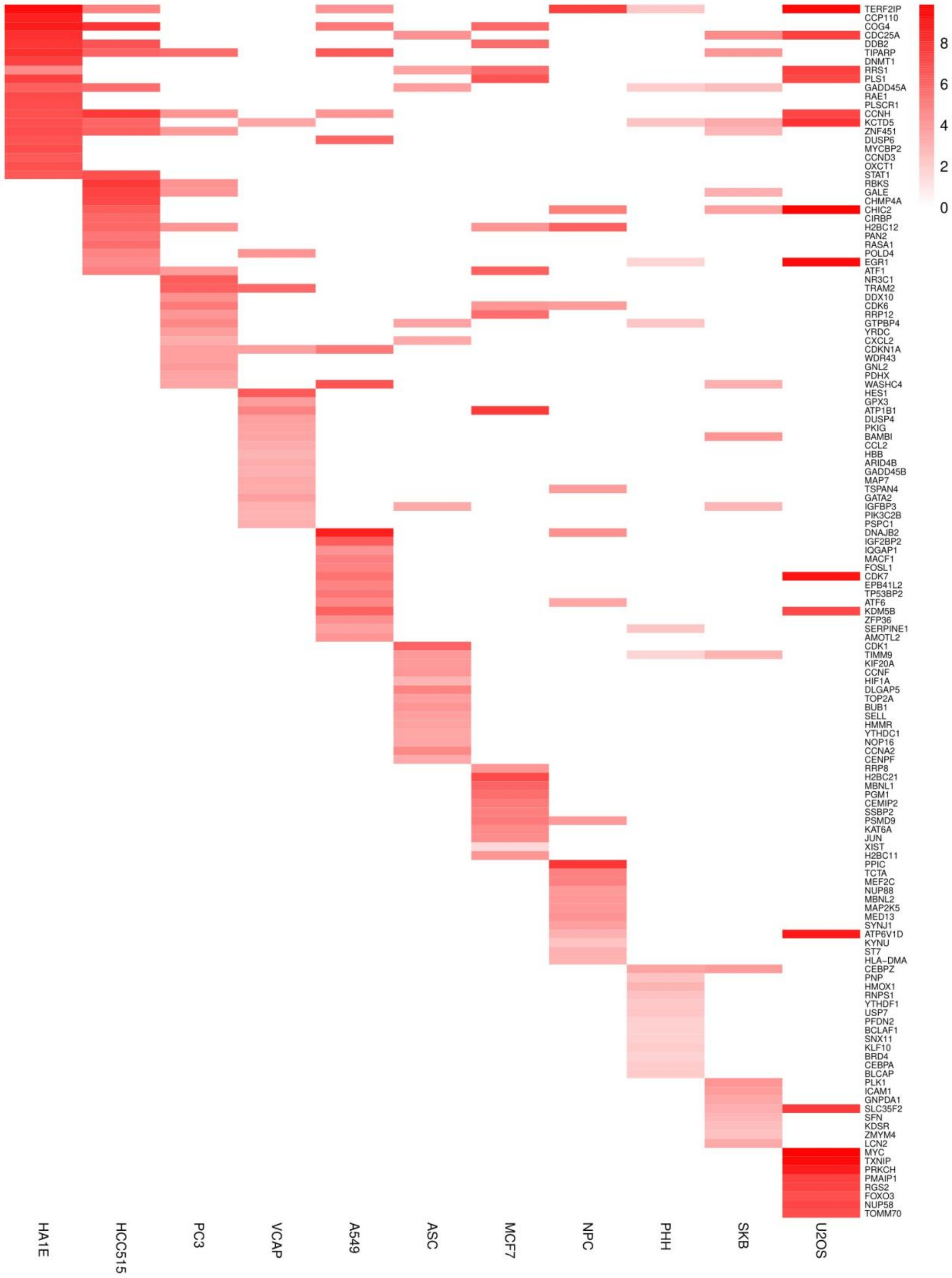


**Figure S30. Cycloheximide -induced upregulated gene signatures across cell lines, Related to Figure 6**

Heatmap comparing the top 20 genes upregulated by cycloheximide across indicated cell lines based on CMAP transcriptional profiles.

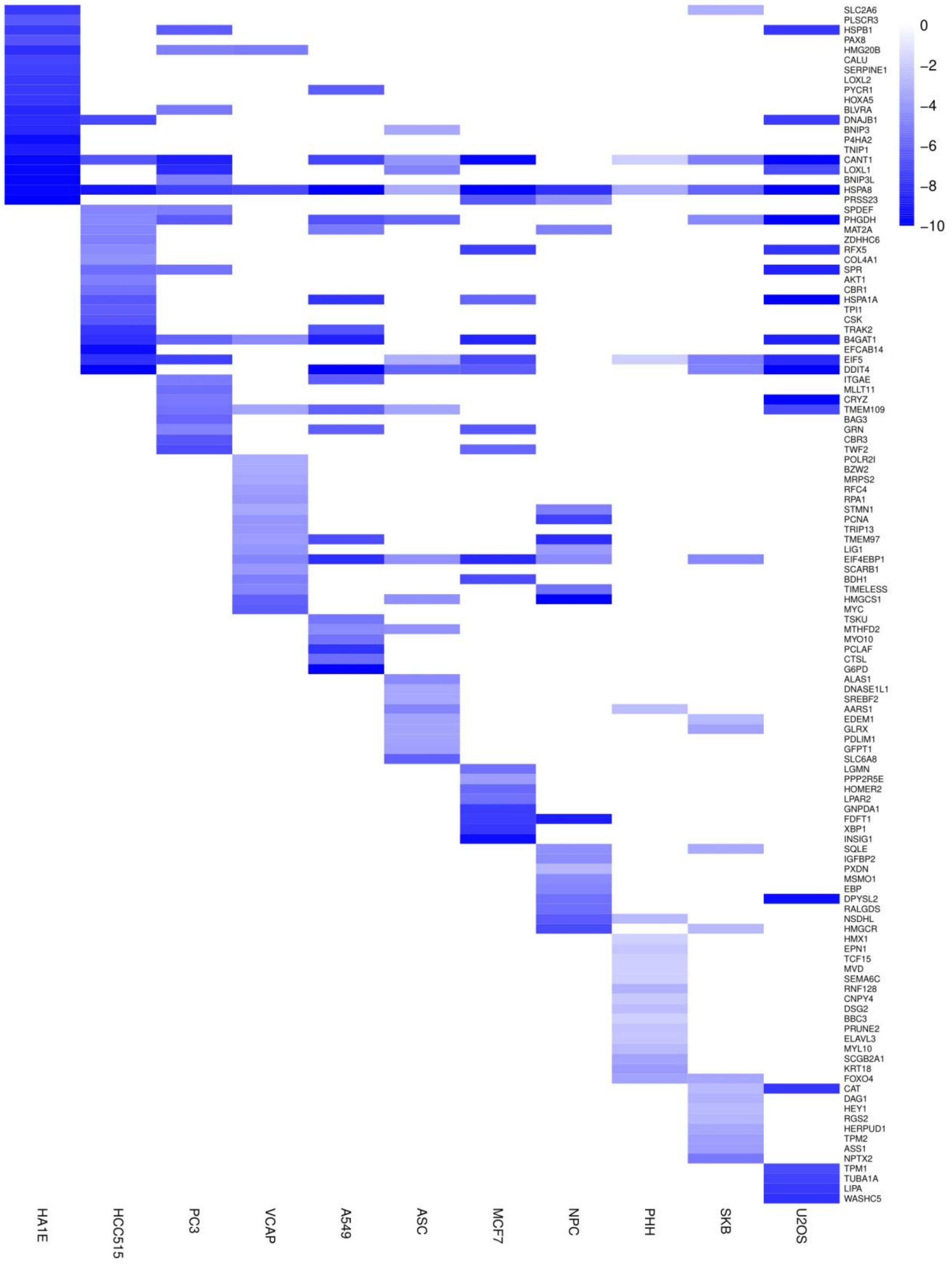


**Figure S31. Cycloheximide -induced downregulated gene signatures across cell lines, Related to Figure 6**

Heatmap comparing the top 20 genes downregulated by Cycloheximide across indicated cell lines based on CMAP transcriptional profiles.

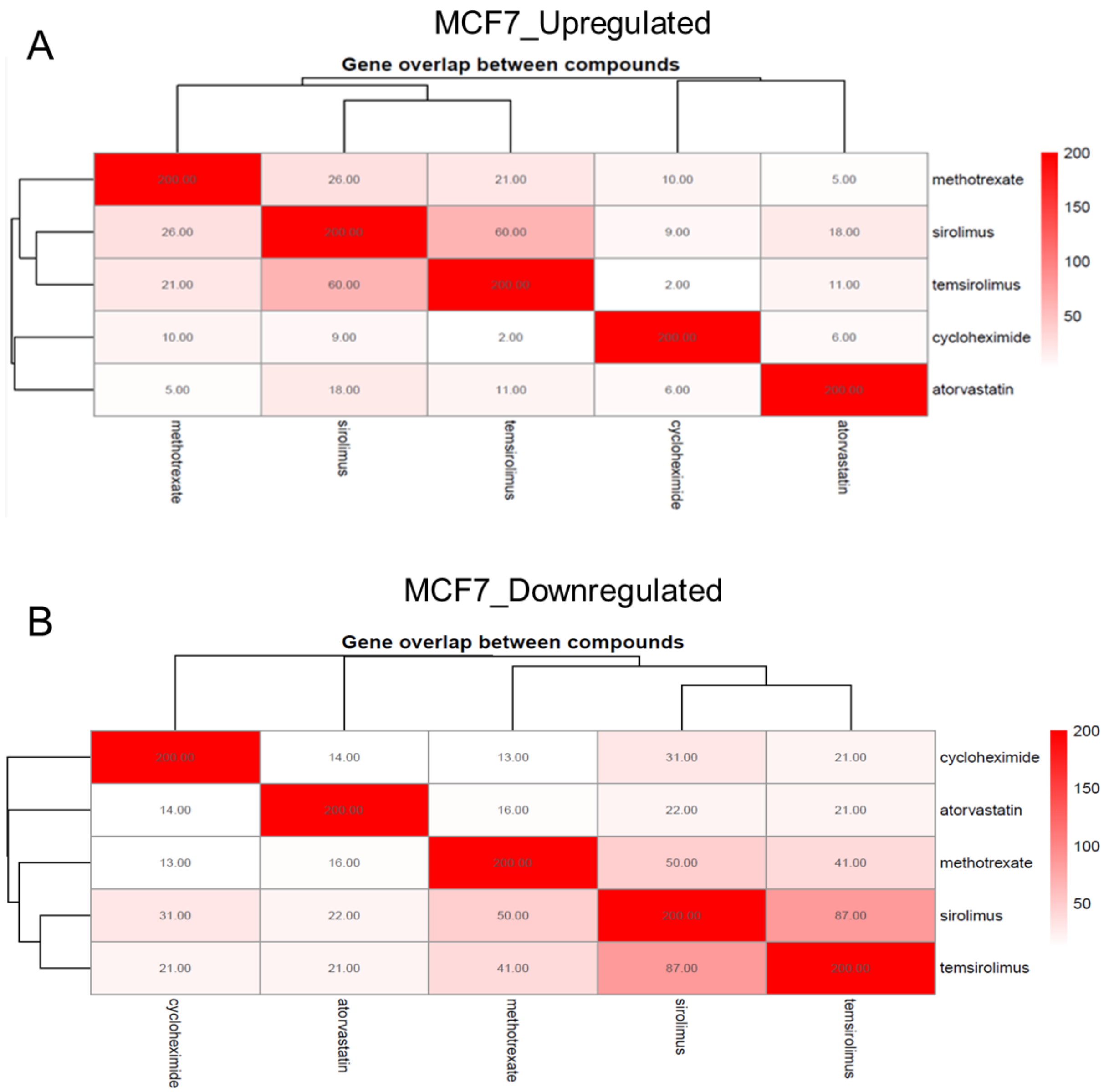


**Figure S32. Gene overlap among compound-induced transcriptional signatures in MCF7 cells, Related to Figure 6**

Heatmaps showing the overlap of the top 200 genes regulated by sirolimus, temsirolimus, methotrexate, atorvastatin, and cycloheximide based on CMAP analysis. Numbers indicate the count of shared genes between compound pairs. (A) Upregulated genes in MCF7. (B) Downregulated genes in MCF7.

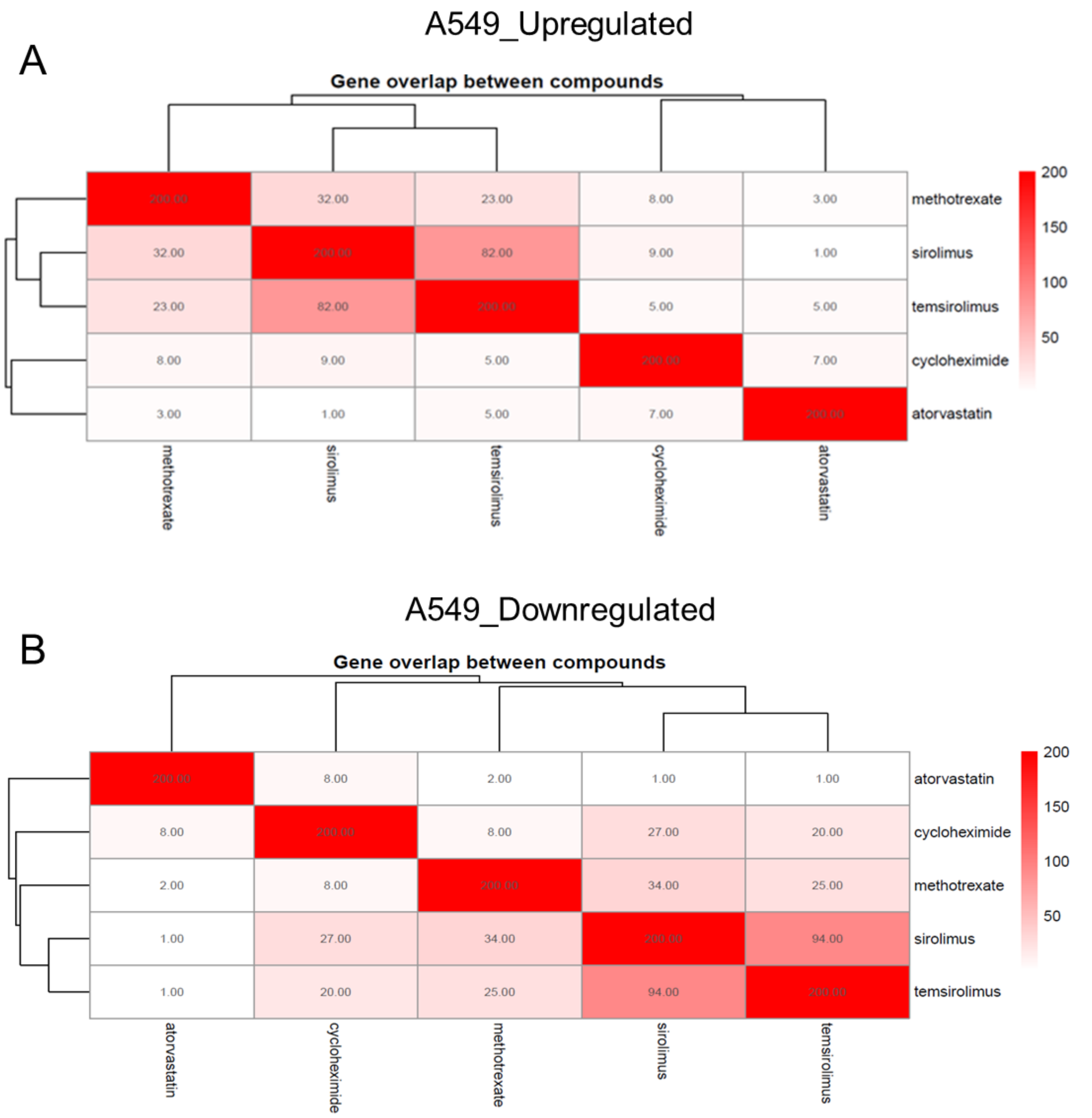


**Figure S33. Gene overlap among compound-induced transcriptional signatures in A549 cells, Related to Figure 6**

Heatmaps showing the overlap of the top 200 genes regulated by sirolimus, temsirolimus, methotrexate, atorvastatin, and cycloheximide based on CMAP analysis. Numbers indicate the count of shared genes between compound pairs. (A) Upregulated genes in A549. (B) Downregulated genes in A549.

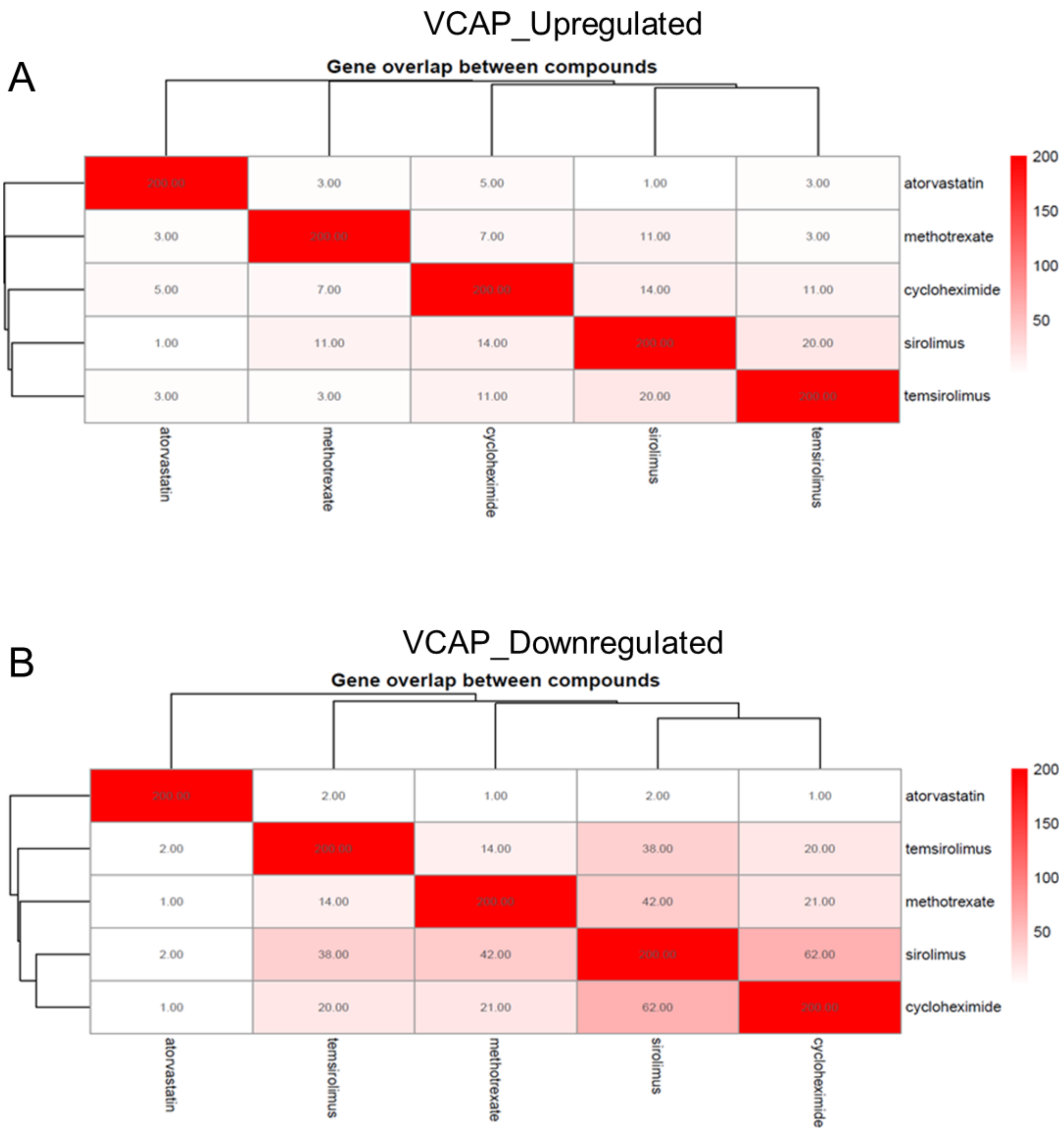


**Figure S34. Gene overlap among compound-induced transcriptional signatures in VCAP cells, Related to Figure 6**

Heatmaps showing the overlap of the top 200 genes regulated by sirolimus, temsirolimus, methotrexate, atorvastatin, and cycloheximide based on CMAP analysis. Numbers indicate the count of shared genes between compound pairs. (A) Upregulated genes in VCAP. (B) Downregulated genes in VCAP.

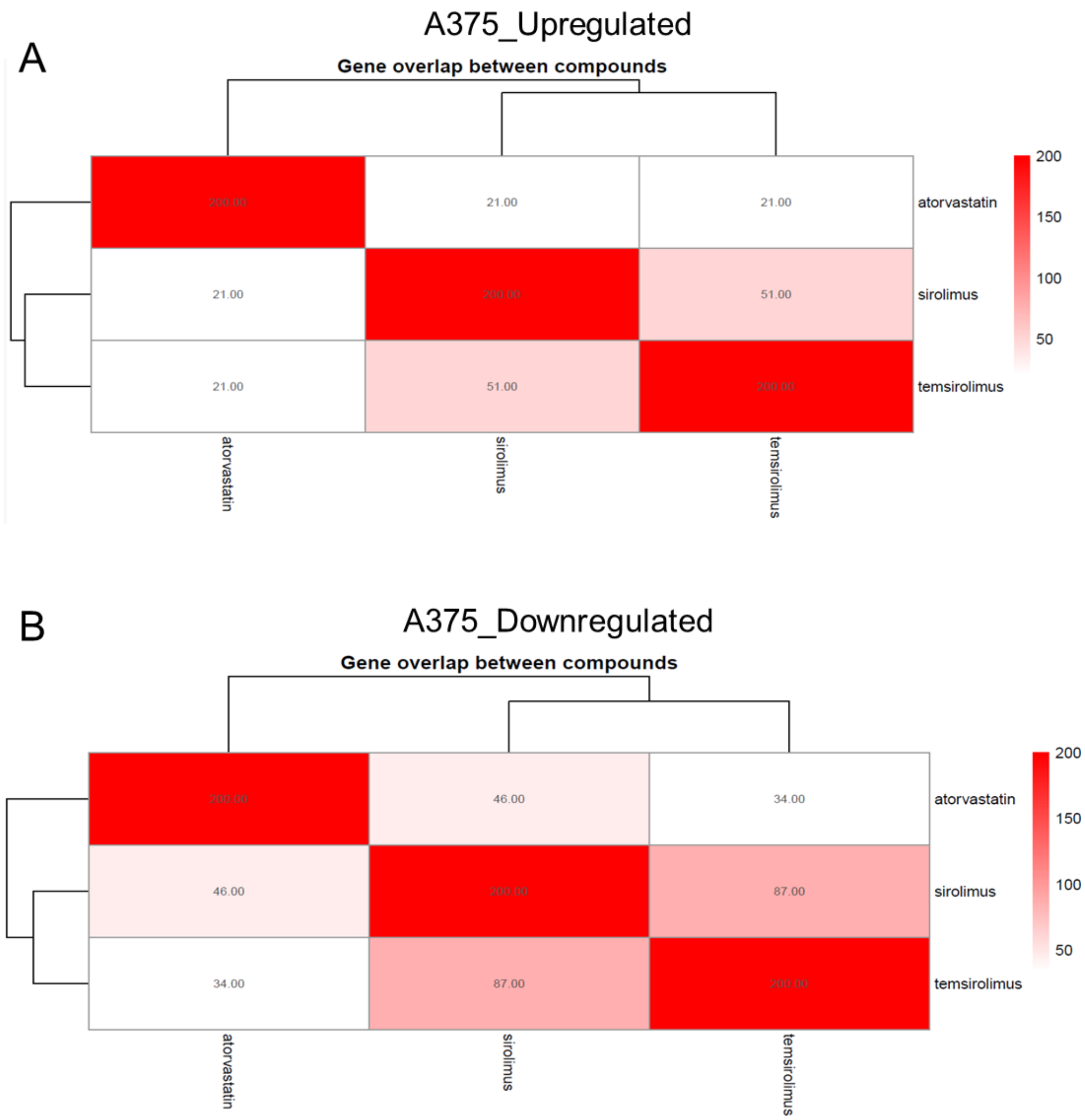


**Figure S35. Gene overlap among compound-induced transcriptional signatures in A375 cells, Related to Figure 6**

Heatmaps showing the overlap of the top 200 genes regulated by sirolimus, temsirolimus, methotrexate, atorvastatin, and cycloheximide based on CMAP analysis. Numbers indicate the count of shared genes between compound pairs. (A) Downregulated genes in A375. (B) Upregulated genes in A375.

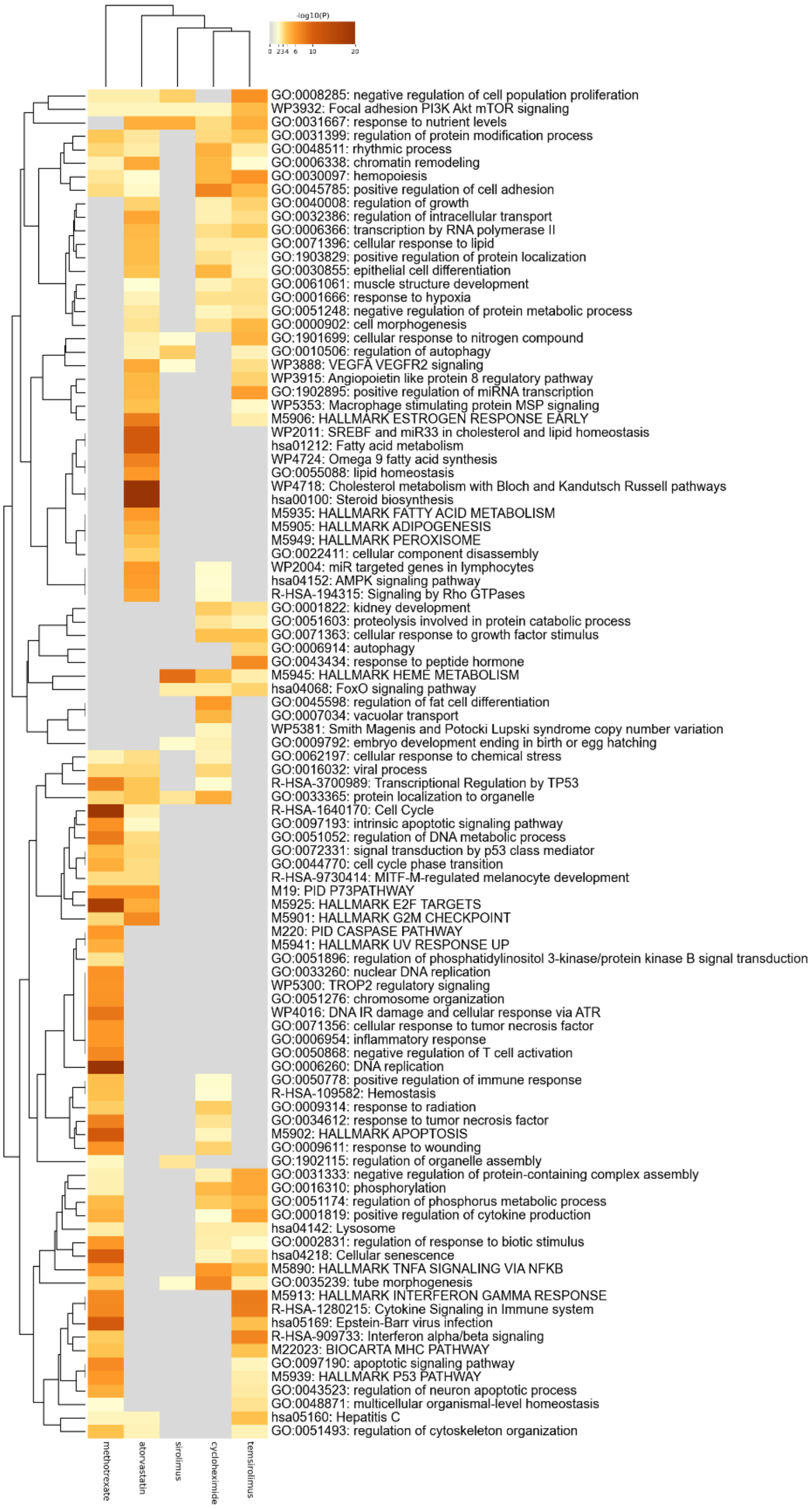


**Figure S36. Convergent pathway responses to metabolic perturbations in VCAP prostate cancer cells, Related to Figure 6**
Heatmap of pathway enrichment derived from the top 200 upregulated genes following treatment with methotrexate, sirolimus, temsirolimus, atorvastatin, or cycloheximide.

Color scale indicates enrichment significance (−log10 P value).

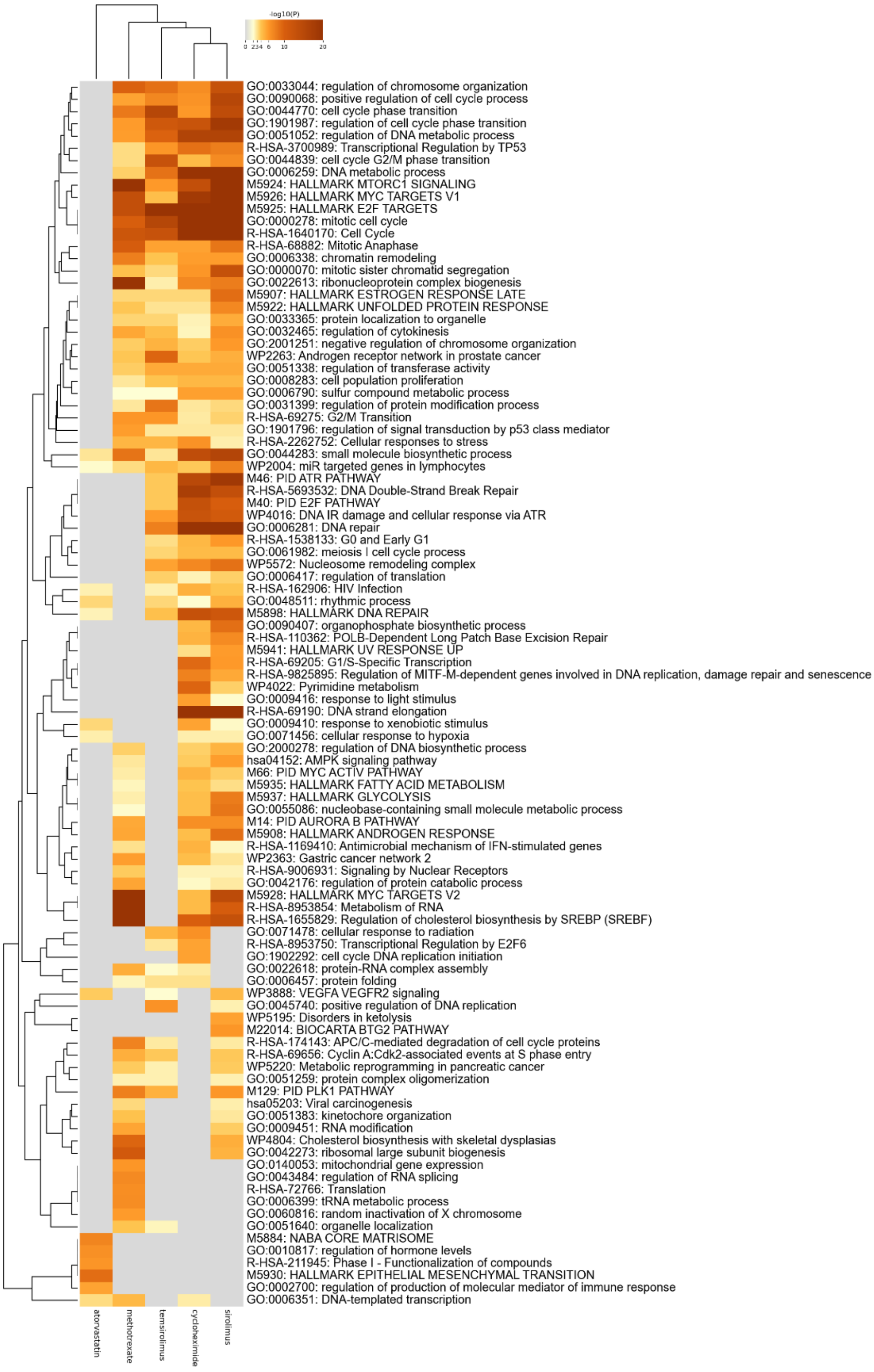


**Figure S37. Convergent pathway responses to metabolic perturbations in VCAP prostate cancer cells, Related to Figure 6**
Heatmap of pathway enrichment derived from the top 200 downregulated genes following treatment with methotrexate, sirolimus, temsirolimus, atorvastatin, or cycloheximide.

Color scale indicates enrichment significance (−log10 P value).

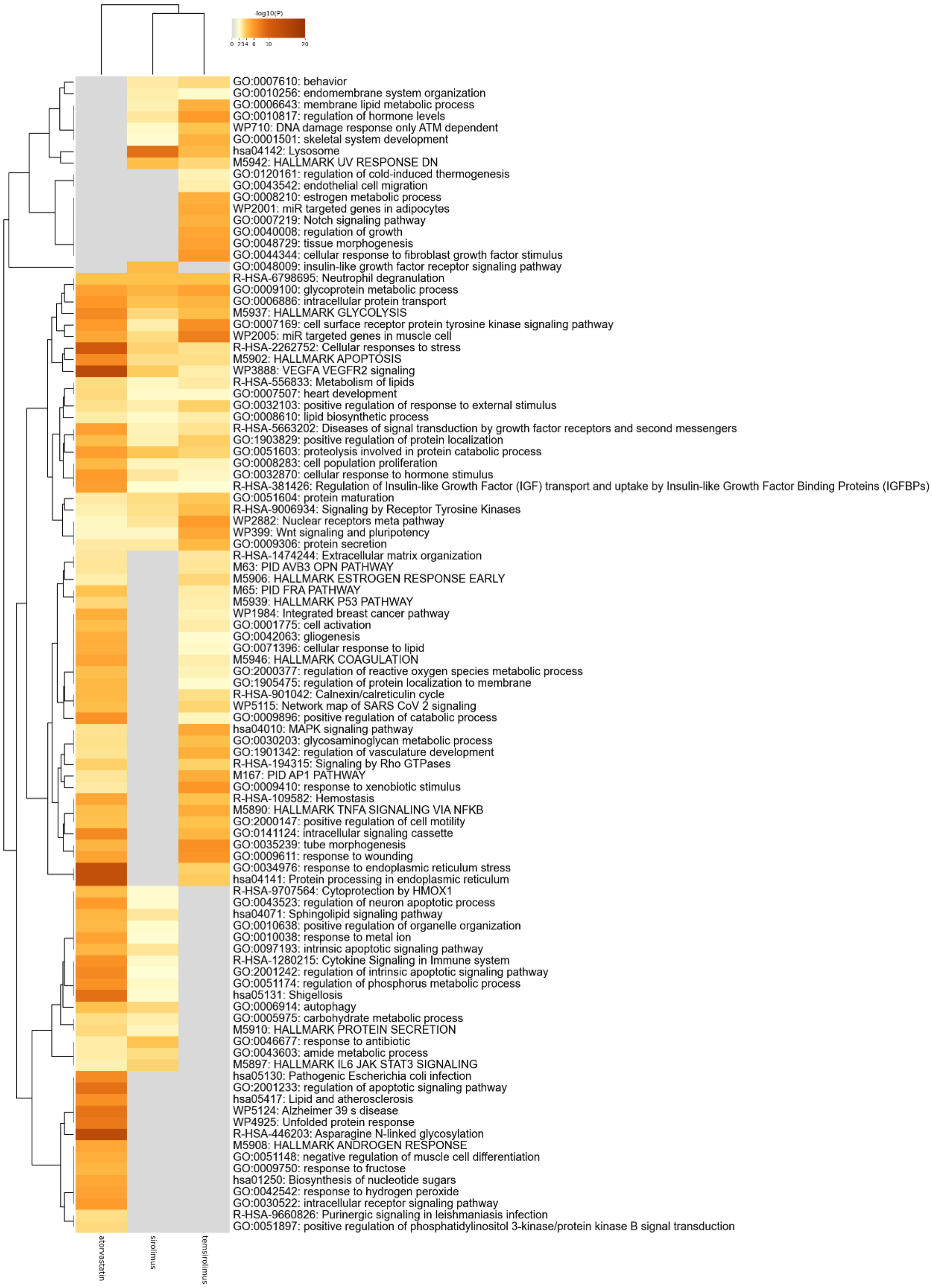


**Figure S38. Convergent pathway responses to metabolic perturbations in A375 melanoma cells, Related to Figure 6**
Heatmap of pathway enrichment derived from the top 200 upregulated genes following treatment with methotrexate, sirolimus, temsirolimus, atorvastatin, or cycloheximide.

Color scale indicates enrichment significance (−log10 P value).

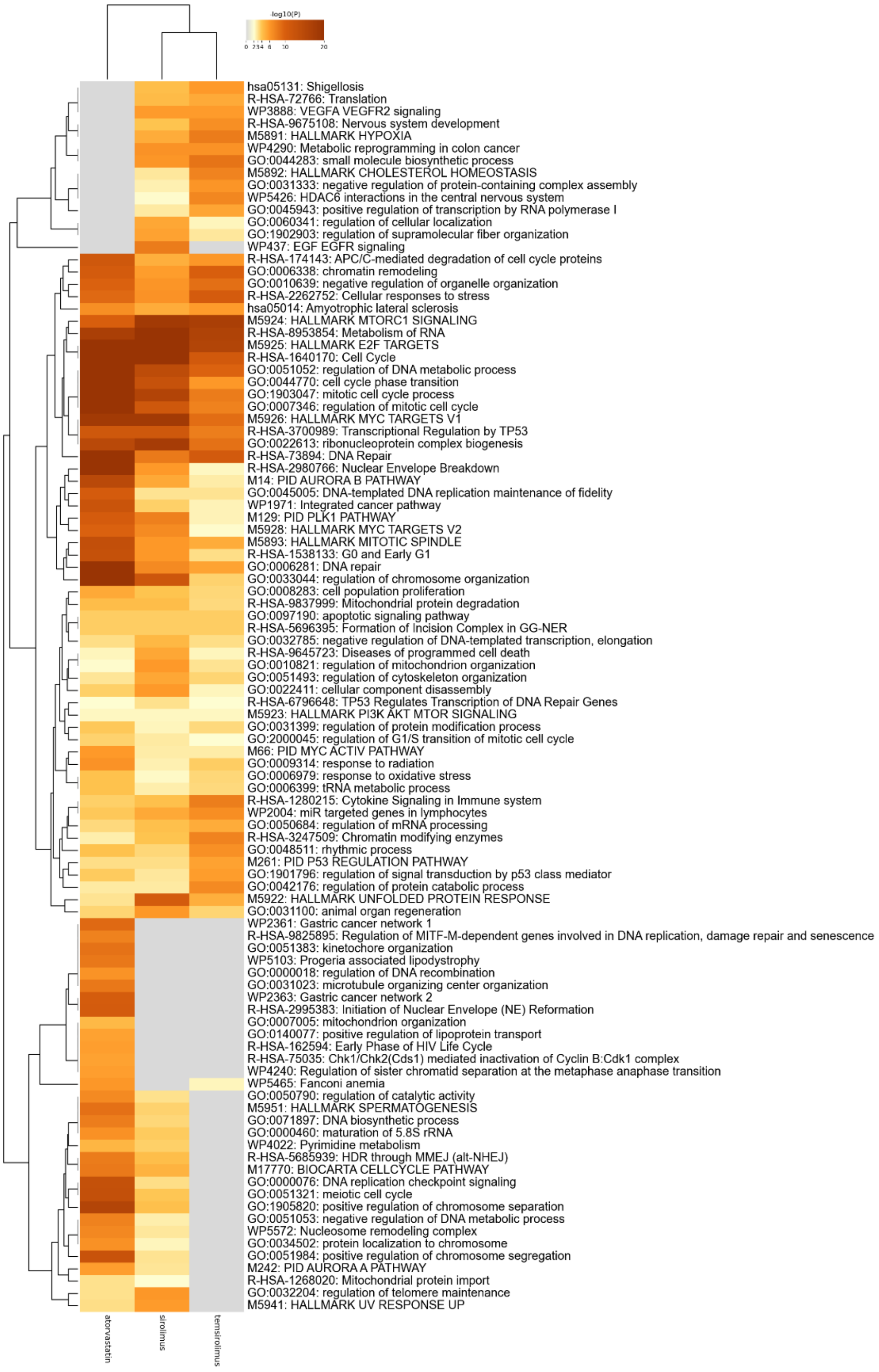


**Figure S39. Convergent pathway responses to metabolic perturbations in A375 melanoma cells, Related to Figure 6**
Heatmap of pathway enrichment derived from the top 200 downregulated genes following treatment with methotrexate, sirolimus, temsirolimus, atorvastatin, or cycloheximide.

Color scale indicates enrichment significance (−log10 P value).

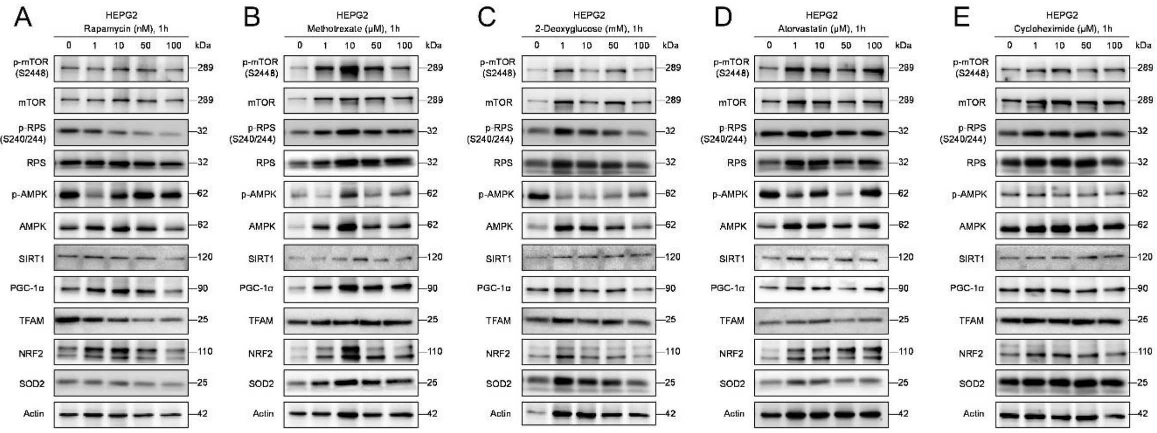


**Figure S40. Metabolic perturbations modulate mTORC1, mitochondrial, and stress signaling in HEPG2 cells, Related to Figure 7**

(A–E) Immunoblot analysis of mTORC1 signaling, metabolic, mitochondrial, and stress-associated markers in HEPG2 hepatocellular carcinoma cells following acute (1 h) treatment with rapamycin (A), methotrexate (B), 2-deoxyglucose (C), atorvastatin (D), and cycloheximide (E) at the indicated concentrations. mTORC1 activity was assessed by phosphorylation of mTOR (S2448) and ribosomal protein S6 (p-RPS, S240/244). Additional markers include p-AMPK/AMPK, SIRT1, PGC-1α, TFAM, NRF2, and SOD2. Actin serves as a loading control.

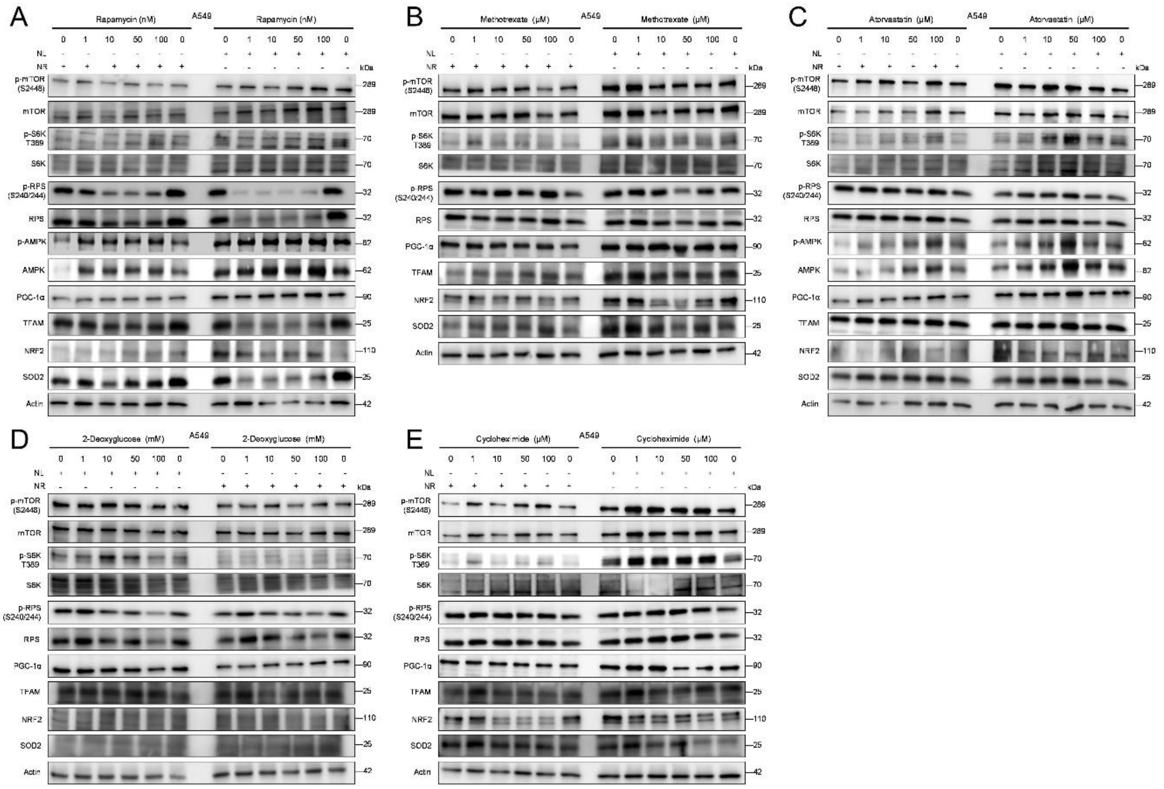


**Figure S41. Nutrient-dependent signaling responses to metabolic perturbations in A549 cells, Related to Figure 7**

(A–E) Immunoblot analysis of mTORC1 signaling, metabolic, mitochondrial, and stress-associated markers in A549 lung adenocarcinoma cells under nutrient-rich (NR; 10% FBS) and nutrient-limited (NL; 1% FBS) conditions following treatment with rapamycin (A), methotrexate (B), atorvastatin (C), 2-deoxyglucose (D), and cycloheximide (E) at the indicated concentrations. mTORC1 activity was assessed by phosphorylation of mTOR (S2448), S6K (T389), and ribosomal protein S6 (p-RPS, S240/244). Additional markers include p-AMPK/AMPK, PGC-1α, TFAM, NRF2, and SOD2. Actin serves as a loading control.

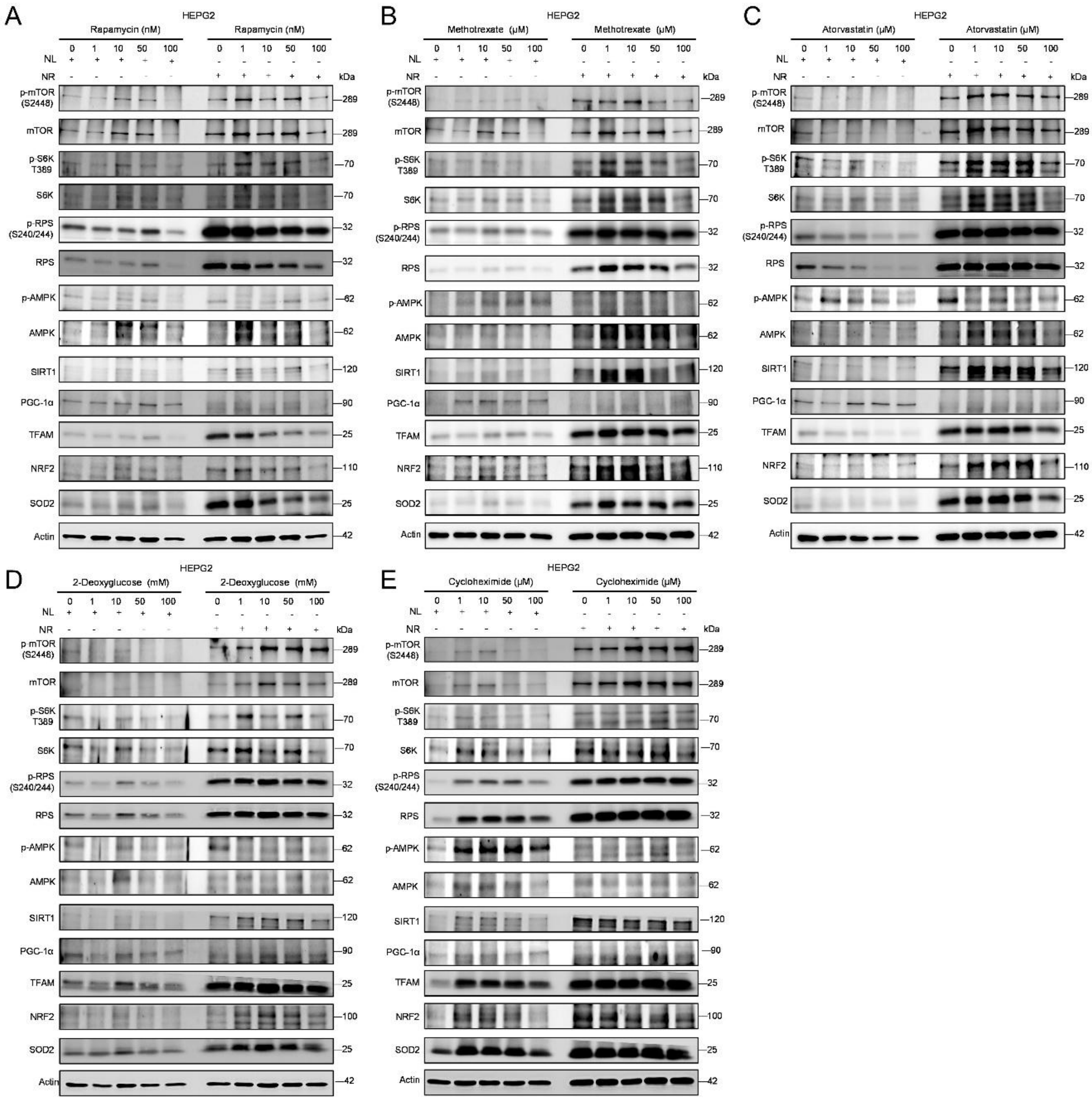


**Figure S42. Nutrient-dependent signaling responses to metabolic perturbations in HEPG2 cells, Related to Figure 7**

(A–E) Immunoblot analysis of mTORC1 signaling, metabolic, mitochondrial, and stress-associated markers in A549 lung adenocarcinoma cells under nutrient-rich (NR; 10% FBS) and nutrient-limited (NL; 1% FBS) conditions following treatment with rapamycin (A), methotrexate (B), atorvastatin (C), 2-deoxyglucose (D), and cycloheximide (E) at the indicated concentrations. mTORC1 activity was assessed by phosphorylation of mTOR (S2448), S6K (T389), and ribosomal protein S6 (p-RPS, S240/244). Additional markers include p-AMPK/AMPK, PGC-1α, TFAM, NRF2, and SOD2. Actin serves as a loading control.

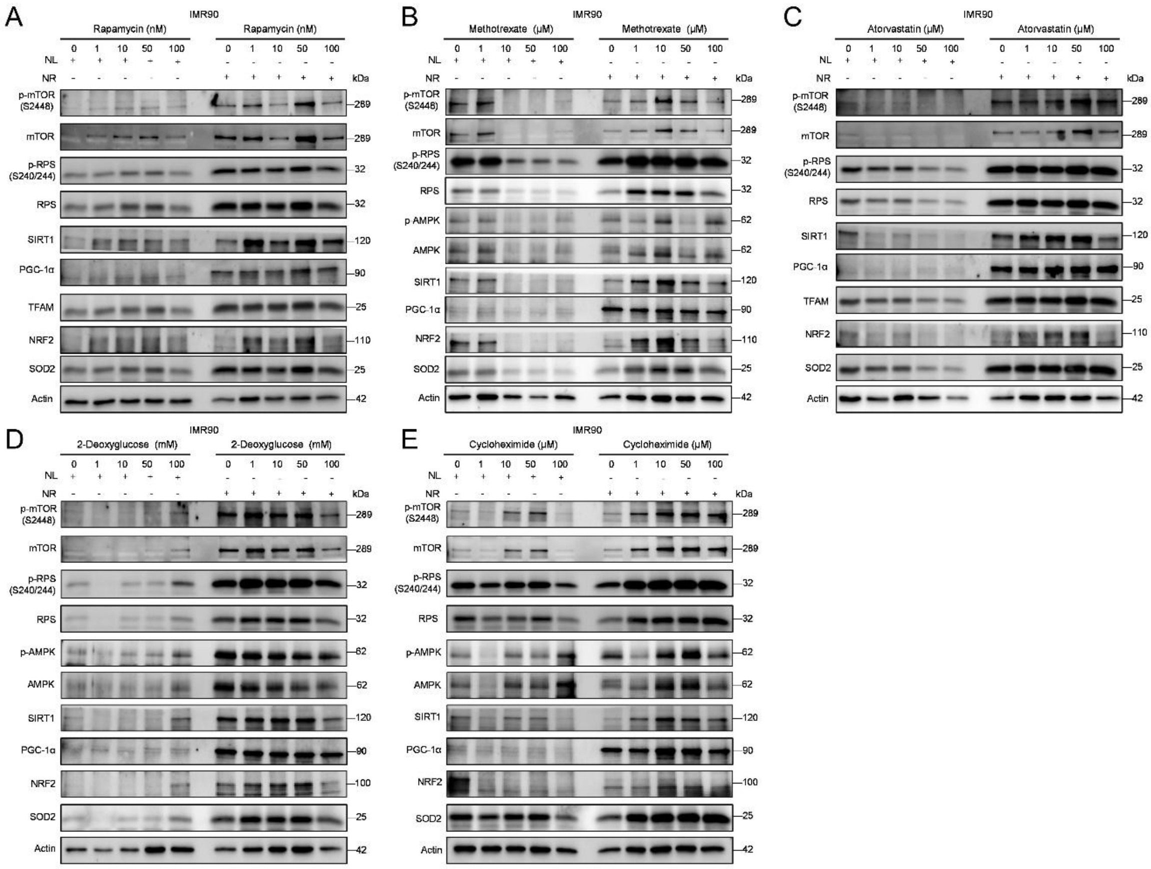


**Figure S43. Nutrient-dependent signaling responses to metabolic perturbations in IMR90 cells, Related to Figure 7**

(A–E) Immunoblot analysis of mTORC1 signaling, metabolic, mitochondrial, and stress-associated markers in A549 lung adenocarcinoma cells under nutrient-rich (NR; 10% FBS) and nutrient-limited (NL; 1% FBS) conditions following treatment with rapamycin (A), methotrexate (B), atorvastatin (C), 2-deoxyglucose (D), and cycloheximide (E) at the indicated concentrations. mTORC1 activity was assessed by phosphorylation of mTOR (S2448), S6K (T389), and ribosomal protein S6 (p-RPS, S240/244). Additional markers include p-AMPK/AMPK, PGC-1α, TFAM, NRF2, and SOD2. Actin serves as a loading control.
